# Selective persistence of HIV-1-infected T cell clones can occur through immune reprogramming driven by defective, transcriptionally active proviruses

**DOI:** 10.1101/2025.09.22.676989

**Authors:** Martin V. Hamann, Lisa Brauckmann, Christoph Schwarz, Michael Spohn, Ramon Stoeck, Sabrina M. Leddy, Julie Frouard, Maisha Adiba, Riekje Winzer, Sanamjeet Virdi, Adam Grundhoff, Cedric Feschotte, Frauke Muecksch, Nadia R. Roan, Eva Tolosa, Ulrike C. Lange

## Abstract

People living with HIV (PLWH) on antiretroviral therapy (ART) accumulate primarily defective proviral sequences in genomes of often clonally expanded CD4+ HIV-1 target cells. The majority of viral-derived DNA is transcriptionally active and preferentially found at distinct genomic loci suggesting a selective process driven by integration site-specific crosstalk between viral and host sequences. Focusing on one of the most prominent selected integration loci, the BTB Domain and CNC Homolog 2 (*BACH2*) gene, we here show mechanistic insights how CD4+ T cells are functionally reprogrammed via exaptation of provirus-derived regulatory sequences during long-term ART. Using a cellular model of *BACH2*-integrated proviruses, we find that proviral transcription drives aberrant BACH2 protein levels that escape autoregulatory feedback and impose BACH2-dependent transcriptomic changes. By mimicking these changes in primary CD4+ T lymphocytes, we observe that BACH2 drives reprogramming of cells toward a proliferative, precursor memory-like type. These reprogrammed CD4+ T cells possess traits of immune evasion and cellular survival that are signatures of persistent HIV reservoir cells in PLWH. Inhibition of provirus transcriptional activity can mitigate exaptation, suggesting a strategy to offset HIV-driven differentiation and expansion of CD4+ T cells. Finally, our data suggest that provirus exaptation at a second prominently selected proviral integration gene, the Signal Transducer And Activator of Transcription 5B (STAT5B) gene, drives a contrary, effector-like T cell fate, suggesting a multifaceted impact of exaptation on immune homeostasis. Overall, our data suggest that transcriptionally active proviruses, even if structurally defective, modulate target cells through insertional activation of integration genes, a process which we postulate to contribute to the complex immune modulation and dysregulation experienced by ART-suppressed PLWH.

## Introduction

People living with HIV-1 (PLWH) harbor chromosomally integrated viral genomes in HIV-1 reservoir cells. These proviruses are remnants of the initial infection and persist despite suppressive antiretroviral therapy (ART) (1,2). One central challenge in current HIV research is understanding how these persistent proviruses lead to immune dysfunction that has been linked to reservoir maintenance and non-AIDS co-morbidities in PLWH and devising strategies how to halt this process (3). The provirus reservoir in PLWH on long-term ART has been extensively characterized, and displays three main features: structural defectiveness, transcriptional activity and clonality. Less than 2% of proviruses can support replication and viral re-bound upon ART cessation. The vast majority are sequence-defective, demonstrating point mutations or deletions (4–8). Although not replication-competent, defective proviruses can impact on immune responses in PLWH: Since ART does not inhibit HIV-1 transcription, defective proviruses can be the source of viral transcripts and peptides that elicit type 1 interferon responses, production of inflammatory cytokines, and deflection of CTL responses (9–13). Defective proviruses have also been suggested as source for non-suppressible residual viremia in PLWH on ART and as driver for HIV-related disease progression and lack of immunological response during therapy (7,14,15). Recent studies suggest that up to 80% of proviruses in PLWH show transcriptional activity (16,17).

The provirus reservoir in PLWH on ART is also highly clonal (18–22). Clonal expansion has been observed for cells with intact as well as defective proviruses. Homeostatic proliferation and antigen-driven stimulation have been suggested as major driving forces for clonal expansion of HIV reservoir cells (23,24). Evidence from longitudinal studies of proviral integration landscapes in PLWH on ART suggest an additional expansion process driven by proviral activity. Time on ART selects for clones with specific integration genes, termed recurrently-<u>d</u>etected integration <u>g</u>enes (RdIGs) (25–31). These RdIGs often appear to be associated with pathways promoting cellular proliferation and/or survival (19,20,30,31). Clonal expansion of cells with proviral integrations at RdIGs has thus been suggested to result from functional crosstalk between provirus and human DNA, in which virus-derived promoter elements cause insertional activation and aberrant expression of integration genes (24,25). While often stated, there is so far little direct evidence and mechanistic understanding of this process of HIV-1 promoter exaptation.

The *BTB And CNC Homolog 2* (*BACH2)* gene has been identified as one of the most prominent RdIGs (30–34). *BACH2* encodes a transcriptional master regulator in B and T lymphocytes, that has been associated with maintaining immune homeostasis by restraining terminal differentiation, promoting memory and regulatory subsets, and ensuring effective, balanced immune responses (35–41). Time on ART selects for *BACH2* provirus integrations that are clustered in congruent transcriptional orientation upstream of the gene’s protein coding exons (29–33). This selection pattern is highly reminiscent of a functional transcriptional crosstalk in terms of promoter exaptation. Indeed, chimeric proviral/*BACH2* transcripts, which are initiated by splicing from the HIV-1 major splice donor site (MSD) into an endogenous splice acceptor site of *BACH2,* have been detected in peripheral blood mononuclear cells (PBMCs) from PLWH (42–44). The selective pressure for clones carrying *BACH2* proviral integrants is remarkably high: *BACH2* integrations and proviral/BACH2 transcripts have been found in up to 40% of long-term suppressed PLWH (32,42,43). Despite this prominence, mechanistic details of how promoter exaptation occurs at the BACH2 locus and how this impacts the host cell are largely unknown. We have previously shown that transcriptional activity of *BACH2*-integrated HIV-1 promoter sequences is not linked to *BACH2* gene promoter activity, indicating that effects driven by promoter-exaptation are potentially governed by distinct regulatory pathways compared to endogenous BACH2 expression (32). It has also been suggested that proviral/BACH2 crosstalk could promote differentiation toward a regulatory T cell (Treg)-like phenotype (43), and that within the Treg population, such crosstalk could contribute to an expanded reservoir population that maintains HIV persistence (42). This conclusion however warrants further investigation, since no sequence-intact *BACH2*-integrated proviruses have so far been reported in PLWH on ART, making their contribution to the replication-competent reservoir little likely (12,23,45–48). Instead, the findings provide a first indication that transcriptional crosstalk between proviruses and their integration gene is a means of how defective proviruses may shape the fate of infected cells during ART.

Building on these observations, we here set out to further define how in particular defective proviruses that are selected to persist at RdIGs during ART could affect CD4+ T cells and in which way this can lead to immune modulation in PLWH. We focus primarily on BACH2-proviral clones given their striking prominence in suppressed PLWH. Using targeted genome engineering, we generated a cellular model system, that replicates *BACH2* proviral integrations observed *in vivo* as unique tool to study transcriptional provirus/human crosstalk at this locus. We further determine how, as a result, aberrant BACH2 expression reprograms primary CD4+ T lymphocytes and thus alters the target cell immune state. We highlight how reprogrammed cells show features of expanded HIV-1 reservoir clones in PLWH, reveal that outcomes of provirus-driven reprogramming are integration-gene dependent and show the potential to halt provirus-mediated insertional activation through compounds that promote HIV-1 latency. Our results emphasize a mechanism by which persistent proviruses, independent of replication competence, have the ability the modulate the immune landscape of PLWH on ART through exaptation of virus-derived regulatory elements.

## Results

### 1. LTR exaptation drives BACH2 protein expression in a cellular model that replicates *in vivo*-selected *BACH2* proviral clones

In order to establish a comprehensive view of BACH2 proviral integrations and the effects of *in vivo* selection, we first undertook comparative profiling of proviral integration sites (ISs) detected in PBMCs of aviremic PLWH on ART and integration sites from *in vitro* infected primary CD4 T cells. Using data from the Retrovirus Integration Database v2.0 (49), we catalogued 11,579 ISs and 3,898 integration genes (IGs) reported *in vivo* (12,30,31,33,34,46,50–54), and 463,206 ISs and 14,192 IGs *in vitro* (26,51,55,56) from a total of 15 studies. Consistent with prior reports (26,30–33), we detected the *BACH2* locus as the most frequent and most *in vivo* selected integration gene (Fig1a, SuppFig1). We also observed as previously described that *in vivo BACH2*-integrations showed significant enrichment towards intron 5 and congruent transcriptional orientation with respect to the gene (Fig1b).

**Figure 1.**
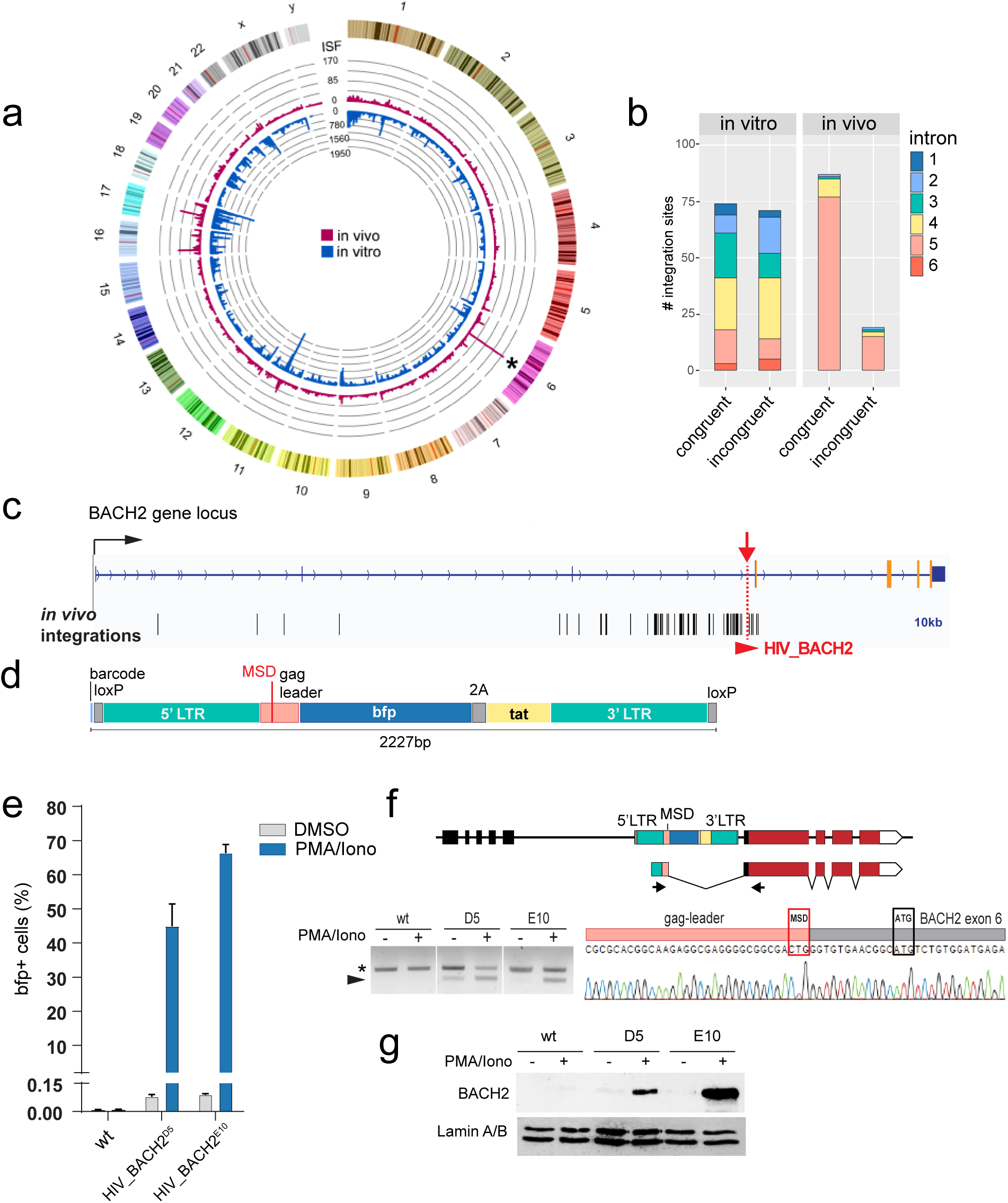
HIV_BACH2 cells model LTR-driven insertional activation of the BACH2 RdIG. (a) Circos plot depicting the genome-wide frequency of proviral integrations per gene locus in PLWH on ART (in vivo) vs HIV-1 infection of CD4+ T cells in vitro. Data on proviral integration sites was extracted from the Retrovirus Integration Database ^45^. Asterisk marks the BACH2 gene locus. ISF: integration site frequency. (b) Frequency of in vivo and in vitro provirus integration sites in the BACH2 gene locus stratified by intron and transcriptional orientation of the provirus in context of the BACH2 gene (congruent, incongruent). (c) Schematic of the BACH2 gene locus. Black bars indicate mapped provirus integrations in PBMCs of PLWH on ART. The site of targeted integration of the HIV-1 reporter (LTRbfp2Atat) 3710bp upstream of BACH2 ATG within intron 5 is marked (red arrow). (d) Schematic of the LTRbfp2Atat reporter that was targeted into the BACH2 locus to generate HIV_BACH2 cell line models (E10 and D5 clones). (e) Percentage of bfp positive HIV_BACH2 cells (E10 and D5 clonal lines) upon exposure to PMA/Ionomycin or DMSO control. Untargeted wild-type parental cells (wt) were used as controls. (f) Detection of LTR/BACH2 chimeric transcripts in HIV_BACH2 cell models (E10 and D5 clonal lines) and untargeted wt cells by non-quantitative RT-PCR. Schematic indicates aberrant splicing from reporter-inherent HIV-1 major splice donor (MSD) site to acceptor site of BACH2 exon 6. Protein-coding sequence of BACH2 is indicated in red. Arrows indicate position of primers used for PCR amplification of chimeric transcripts (arrowhead). Asterisk denotes unspecific PCR product. (g) Western blot analysis of HIV_BACH2^D5/E10^ cells for expression of BACH2 protein. Anti-Lamin A/B antibody was used for blotting and loading control.

While direct phenotyping of clones with BACH2-integrated proviruses from PLWH is currently technically not feasible, we next generated a cellular model to study how proviral sequences interact with the BACH2 integration locus. To replicate integration features of *in vivo*-selected BACH2-proviruses, we used CRISPR/Cas9-mediated genome engineering of a leukemia cell line and targeted a minimal HIV-1 reporter into intron 5 of the *BACH2* gene, the precise region that BACH2 integrants are selected for in PLWH (Fig1c, SuppFig2a). Two clonal lines, HIV_BACH2^D5^ and HIV_BACH2^E10^ were generated, that carry a heterozygous (D5) or homozygous (E10) insertion of the HIV-1 reporter in congruent transcriptional orientation at the targeted site (Fig1c). The reporter consists of an expression cassette for blue fluorescent protein (bfp) and HIV-1 transcriptional activator tat, flanked by two HIV-1 long terminal repeat (LTR) sequences (Fig1d). Of note, the gag leader sequence containing HIV-1 major splice donor site (MSD) is included in the reporter upstream of bfp, allowing for splicing of LTR-driven transcripts. Clonal lines were validated for correct reporter insertion using Northern blotting and Sanger sequencing (SuppFig2b, c).

In both clonal lines, transcriptional activity of the LTR (as read out by bfp expression) was minimal but could be induced by exposure to phorbol myristate acetate (PMA) PMA/Ionomycin (Iono), a potent HIV-1 latency reversing treatment (Fig1e). This is consistent with our previous findings on HIV-1 reporter integrations into *BACH2* intron 5 (32). We next tested whether PMA/Iono stimulation can lead to generation of chimeric LTR-BACH2 transcripts. Using primers targeting the reporter LTR and exon 6 of *BACH2*, we could indeed amplify LTR-BACH2 transcripts in cDNA derived from PMA/Iono-treated HIV_BACH2^D5/E10^ cells (Fig1f). Sanger sequencing confirmed that chimeric transcripts were generated through aberrant splicing between the reporter MSD and the *BACH2* exon 6 splice acceptor site. To assess whether chimeric transcripts were translated to full length BACH2 protein, we carried out Western blot analysis of whole cell lysates of uninduced and PMA/Iono-treated HIV_BACH2^D5/E10^ cells. As shown in Fig1g, transcriptional induction by PMA/Iono indeed resulted in induction of BACH2 protein expression. These results demonstrate that the HIV_BACH2^D5/E10^ lines provide a suitable *in vitro* model to study exaptation of proviral sequences, and insertional activation at the *BACH2* RdIG.

### 2. BACH2 integrated proviral sequences activate the *BACH2* gene expression program

Using our HIV_BACH2 model, we next went on to study downstream effects of LTR exaptation at the *BACH2* locus. To do so, we employed a tripartite CRISPR activator (CRISPRa) system, termed synergistic activation mediator (SAM), which allows for guide (g) RNA-directed recruitment of the VP64, p65 and HSF1 transcriptional activator domains to specified targets (57). Leveraging our previously validated gRNAs for efficient activation of the HIV-1 LTR and the *BACH2* gene promoter (32,58), we set out to assess transcriptomic changes induced by SAM-mediated activation of the *BACH2*-integrated HIV reporter and contrast these to the transcriptomic programme induced by activation of the endogenous *BACH2* promoter. In brief, HIV_BACH2^E10^ cells were transiently transfected with LTR-targeting (LTR-SAM) or *BACH2*-targeting (BACH2-SAM) SAM. Expression of SAM components was linked to a fluorescent marker to allow for selection of successfully transfected cells. Wild-type (wt) cells, which harbor the endogenous *BACH2* locus without HIV-1 reporter integrant were used as controls. 72 hours post transfection SAM-expressing cells were enriched by flow cytometry and subjected to transcriptomic profiling (Fig2a).

**Figure 2.**
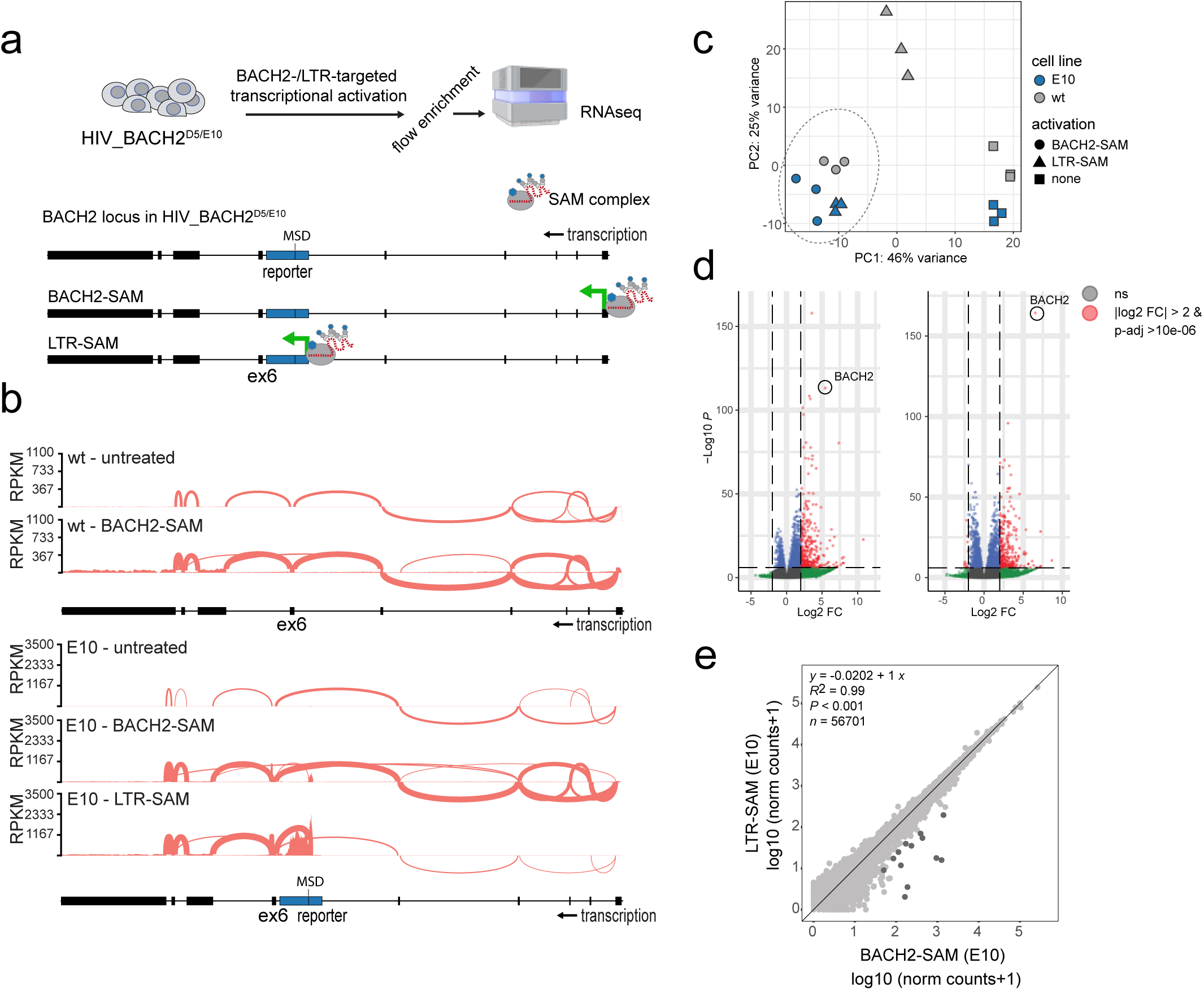
LTR exaptation drives BACH2 gene program in HIV_BACH2 cells. (a) Schematic depicting LTR- (LTR-SAM) and BACH2- (BACH2-SAM) targeted transcriptional activation in HIV_BACH2^D5/E10^ cells and parental wild type cells (wt) using the synergistic activation mediator (SAM) system. SAM-treated cells were subjected to RNAseq analysis. (b) Sashimi plots depicting the BACH2 transcript splicing patterns in BACH2-SAM- and LTR-SAM-treated wt and HIV_BACH2^E10^ cells. (c) Principal component analysis based on RNAseq datasets of SAM-treated wt and HIV_BACH2^E10^ cells as well as untreated controls. (d) Volcano plots depicting differentially expressed genes (DEGs) in HIV_BACH2E10 cells upon SAM activation of endogenous BACH2 promoter (left panel) and SAM activation of BACH2-integrated LTRbfp2Atat reporter (right panel). Cutoffs set at |log2FC|>2 and padj<1x10E-6. (e) Correlation analysis of DEG genes in HIV_BACH2E10 cells upon LTR-SAM and BACH2-SAM activation. Pearson statistical method, R2 and p values indicated.

We first assessed effects of BACH2-SAM treatment and observed that BACH2 gene transcription was induced in HIV_BACH2^E10^ and wt control cells showing the same transcript splicing pattern (Fig2b). Only minimal transcriptional induction of the BACH2-integrated reporter was observed in BACH2-SAM HIV_BACH2^E10^ cells, which confirms our previous results that activity of integrated LTRs within the BACH2 intron 5 is independent of BACH2 locus activity (32). Next, we examined the impact of LTR-SAM treatment: In HIV_BACH2^E10^ cells, LTR-SAM induced HIV reporter transcripts as expected, but additionally a prominent splicing activity from the reporter-inherent MSD to BACH2 exon 6 was observed, giving rise to chimeric LTR/BACH2 transcripts. The splicing pattern of these chimeric transcripts downstream of exon 6 was reminiscent of endogenous BACH2 transcripts (Fig2b).

In order to assess the extent to which *BACH2* transcription induced by the LTR versus the endogenous promoter differentially affects the overall transcriptional profile of the cells, we performed RNA-seq on the SAM-induced HIV_BACH2^E10^ cells. As shown in Fig2c, in principal component space, LTR-SAM and BACH2-SAM-treated HIV-BACH2^E10^ samples segregate close to one another together with BACH2-SAM wt cells and away from LTR-SAM wt cells and non-treated wt or HIV_BACH2^E10^ cells. This indicates that transcriptomic changes induced by LTR-driven insertional activation of BACH2 mirror changes driven by endogenous BACH2. This was confirmed by differential gene expression analysis, which showed a highly similar pattern of differentially transcribed genes driven by LTR-driven versus endogenous BACH2 activation (Fig2d,e). Gene set enrichment analysis of differentially expressed genes (DEGs) in both comparisons further demonstrated that near identical cellular pathways were affected (SuppFig3a,b). Hence, these data indicate that transcriptional activity of *in vivo* selected BACH2-integrated proviruses can result in transcriptomic reprogramming driven by provirus-induced, aberrant BACH2 expression.

### 3. Aberrant BACH2 expression in CD4+ lymphocytes drives cells toward a proliferative precursor memory-like fate

We next set out to explore how aberrant activation of the *BACH2* gene program, as observed via provirus promoter exaptation in our cell line models, could impact primary CD4+ T cells, as main HIV target cells *in vivo*. In this context it is noteworthy that dysregulated BACH2 expression has previously been reported to cause disease: *BACH2* haploinsufficiency can lead to hypogammaglobulinemia and immune dysregulation, manifesting clinically as recurrent infections and chronic intestinal inflammation (59). The *BACH2* gene locus has also been prominently associated with conditions linked to immune imbalance in genome-wide association studies (GWAS) (SuppFig4) (60). All characterized single nucleotide polymorphisms in these studies mapped to non-coding, intronic or upstream regions of *BACH2* with highest occurrence at the 5’ end of the locus and thus coinciding with regions that likely regulate BACH2 transcription and expression levels.

In order to investigate how elevated *BACH2* expression as a result of LTR exaptation could alter functions and differentiation states of primary CD4+ T cells, we used a lentiviral overexpression system (Fig3a). In short, CD4+ lymphocytes isolated from PBMCs of HIV-seronegative donors were transduced with lentiviruses coding for cDNAs of either BACH2 (LV-BACH2) or green fluorescent protein (LV-GFP) as negative control. By linking BACH2- and GFP-cDNA expression to expression of a surface tag (SBP-dLNGFR), transduced cells could be enriched using magnetic beads (Fig3b). Transduced cells showed robust expression of BACH2 or GFP RNA and protein over a time course of 25 days as assessed by RT-qPCR and flow cytometry (SuppFig5a).

**Figure 3.**
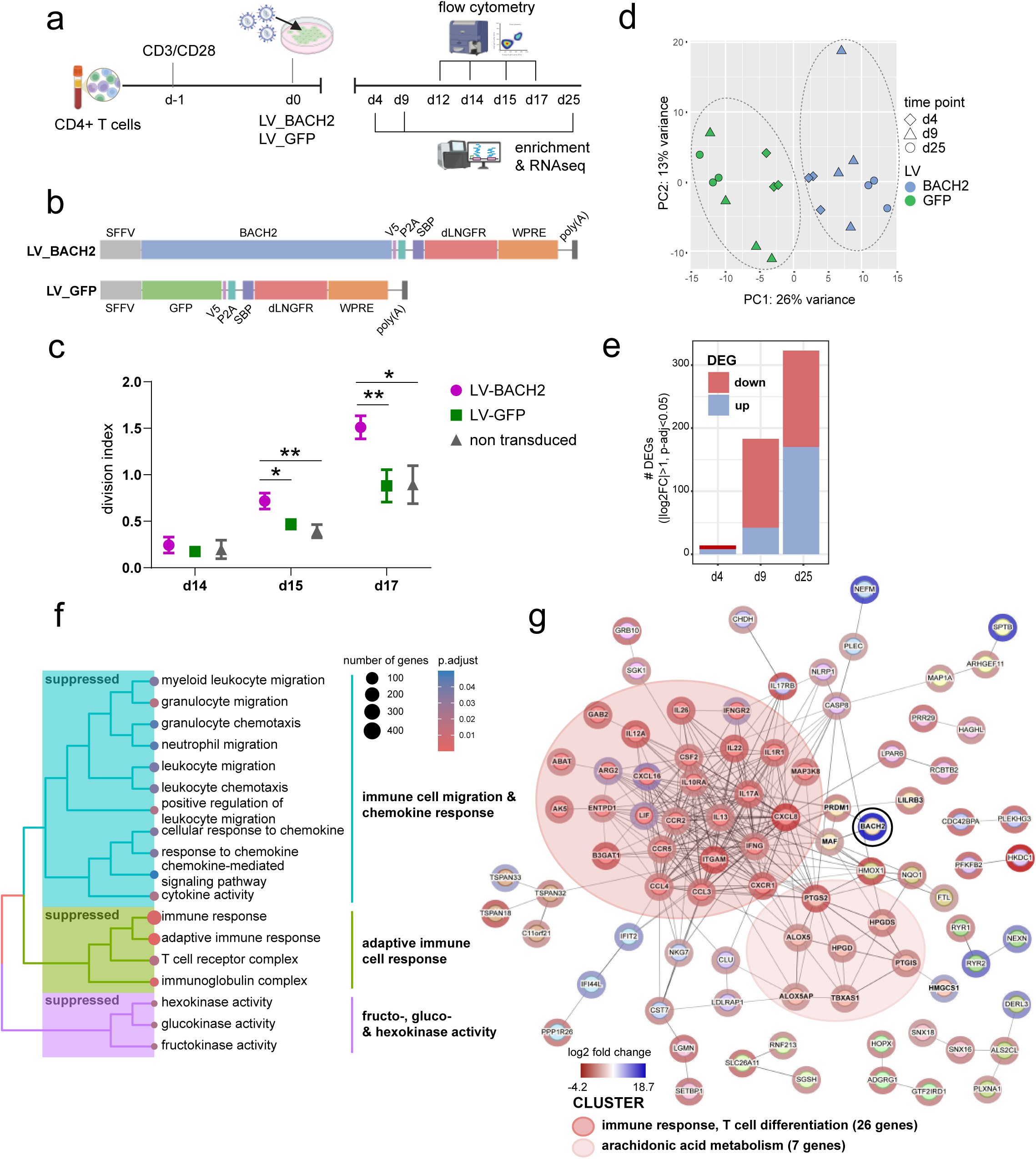
Aberrant BACH2 expression induces proliferation and suppresses effector fates in primary CD4+ T cells. (a) To mimic aberrant BACH2 expression, CD3/CD28-activated CD4+ T cells were transduced with LV-BACH2 and control LV-GFP lentivirus and assessed longitudinally for proliferative behavior (flow-cytometry based) and transcriptomic changes. (b) Schematic of LV-BACH2 and LV-GFP expression cassettes. (c) Cell tracer-based assay for proliferative behavior of LV-BACH2, LV-GFP and non-transduced cells. Cells were stained at d12 post transfection and followed on d14, d15 and d17. Statistical analysis was done using unpaired t-test, *p<0.05, **p<0.03. (d) Principal component analysis based on RNAseq datasets of SPB-dLNGFR-enriched LV-BACH2 and LV-GFP transduced CD4+ T cells at d4 (n=3), d9 (n=4) and d25 (n=3) post transduction. (e) Number of differentially expressed genes (DEG; cutoffs set at |log2FC|>1, padj<0.05) between LV-BACH2 and LV-GFP transduced CD4+ T cells at d4, d9 and d25 post transduction. (f) Gene set enrichment analysis (GSEA) based on DEGs between LV-BACH2 and LV-GFP transduced CD4+ T cells at d9 post transduction delineates suppressed cellular pathways in LV_BACH2 cells. (g) STRING analysis based on DEG between LV-BACH2 and LV-GFP transduced CD4+ T cells at d25 post transduction.

We first assessed whether constitutive *BACH2* expression affects CD4+ T cell proliferation in this system. LV-BACH2 and LV-GFP-transduced cultures were stained with a marker dye 12 days post-transduction, and cell-division-dependent dilution of the dye was measured for up to five days. We observed a significant increase in proliferation of BACH2-transduced cells compared to GFP-transduced or non-transduced control cells as indicated by raised division index (Fig3c). These data suggest that BACH2 drives expansion of primary CD4+ T cells.

We next determined BACH2-induced changes in the transcriptomic profile of primary CD4+ T cells. BACH2- and GFP-transduced cells were enriched at days 4, 9, and 25 post-transduction and analyzed by RNA-seq. In principle component space, LV-BACH2- and LV-GFP cells increasingly segregate away from one another during the 25 days of analysis (Fig3d). In agreement, differentially expressed gene (DEG) analysis revealed that *BACH2* overexpression progressively drives a new transcriptional program: at day 4 post transduction, 14 genes were differentially expressed, and this number increased to 183 at day 9 and 323 at day 25 (SuppTable1, Fig3e). As expected, *BACH2* transcripts were detected as significantly upregulated at all time points in LV-BACH2 cells.

We went on to assay transcriptomic changes using gene set enrichment analysis (GSEA) and STRING functional network profiling based on DEGs between LV-BACH2 and LV-GFP cells. At day 4, no particular differentially regulated pathway was highlighted, likely due to the low numbers of DEGs at this time point. However, we observe that LV-BACH2 cells show upregulation of RAS protein activator like 2 (*RASAL2),* that has been associated with inhibition of T cell activation and cytokine production (61), as well as downregulation of adhesion g protein-coupled receptor G1 (ADGRG1; also known as *GPR56*) and heme oxygenase 1 (*HMOX1)*, two genes linked to effector fates, namely cytotoxic T cells and T regs, respectively (62,63) (SuppTable1).

At d9, GSEA shows downregulation of cellular pathways associated with chemokine signaling, cellular migration and immune response (Fig3f). Most downregulated DEGs belong to two main networks linked to effector T cell differentiation and function (eg. pro-inflammatory cytokine IFN gamma and chemokine receptor IL17RB (64)) as well as arachidonic acid metabolism, necessary for inflammatory signaling (65–67) (Fig3g). In addition, we observe downregulation of *PRDM1* (encoding B-lymphocyte-induced maturation protein-1 (Blimp1)) a direct target of BACH2. PRDM1 is a master transcriptional regulator that promotes terminal effector differentiation of T cells and reduced *PRDM1* expression levels have been shown to support an early memory T cell phenotype (68,69).

At d25, LV-BACH2 cells show further downregulation of a large subset of genes associated with effector T cells and inflammatory cytokine signaling, including CXC-chemokine receptor 6 (*CXCR6*), interleukins 5 and 13 (*IL5/13*) and granzyme B and H (*GZMB/H*) (Fig4a). Furthermore, we observe upregulation of *TCF7* (encoding T cell factor-1 (TCF-1)) and lymphoid enhancer-binding factor 1 (*LEF-1*), that have been reported to synergistically support formation of T cell memory precursors and acquisition of stemness properties for long-term persistence (70,71). We also find positive enrichment for pathways linked to mitotic activity, DNA repair and ribonucleotide complex binding in LV-BACH2 cells at d25, driven for example by upregulation of the mitotic regulators NIMA-related kinase 2 (*NEK2*) and cytoskeleton associated protein 2 (*CKAP2*) (Fig4a, SuppFig5c). This proliferative transcriptomic signature matches our results using cell tracer assays (Fig3c).

**Figure 4.**
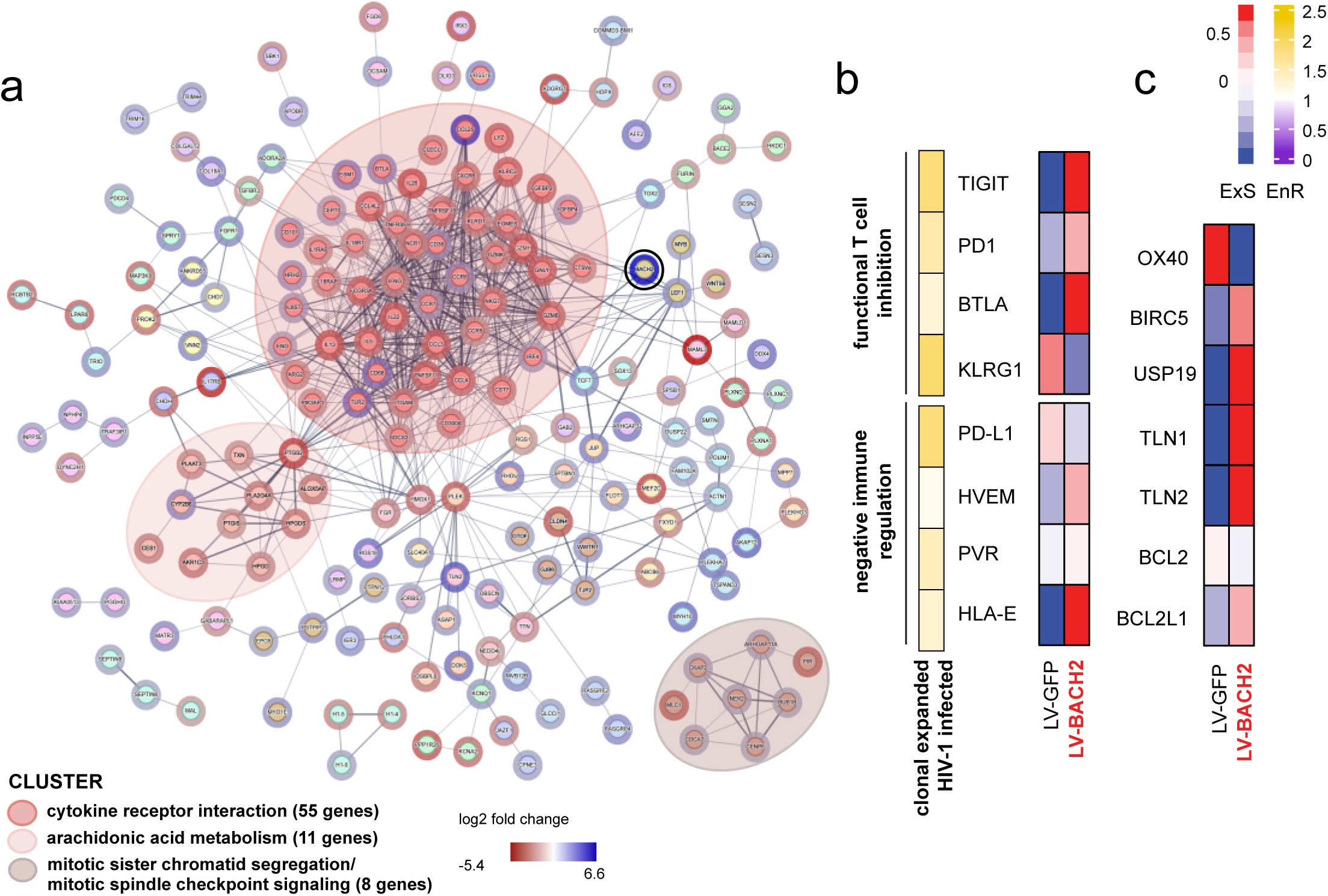
BACH2-reprogrammed CD4+ T cells show precursor memory signature and mimic features of expanded CD4+ T cells in PLWH on ART. (a) STRING analysis based on DEG between LV-BACH2 and LV-GFP transduced CD4+ T cells at d25 post transduction. (b) Heatmap depicting transcript expression scores (ExS) and enrichment scores (EnS) for key genes associated with T cell inhibition and negative immune regulation. ExS are derived from LV-BACH2 and LV-GFP transduced CD4+ T cells (d25 post transduction) and EnS are derived from a published dataset contrasting clonally expanded to non-clonally expanded HIV-1 infected CD4+ T cells from PLWH on ART 44. ExS: scaled normalized transcript levels; EnR: enrichment ratio calculated as the ratio of the proportion of clonal HIV-infected cells divided by the proportion of non-clonal HIV-infected cells (c) Heatmap depicting transcript expression scores (ExS) in LV-BACH2 and LV-GFP transduced CD4+ T cells (d25 post transduction) for key genes associated with anti-apoptotic activity in infected CD4+ T cells in PLWH on ART.

In summary, these data suggest that aberrant BACH2 expression induces a gradual transcriptomic reprogramming, marked by initial inhibition of CD4+ T cell activation and effector type differentiation, followed by initiation of a memory-precursor like program that features a signature of persistence and proliferation.

Finally, we asked whether observed transcriptional reprogramming induced by BACH2 overexpression in primary CD4+ T cells *in vitro* correlates with characteristics of clonally expanded HIV-1 infected cells in PLWH on ART. These cells demonstrate a signature of immunoregulatory markers and immune checkpoint molecules that confer T-cell functional inhibition and resistance to immune-mediated killing (48), coupled with a pro-survival gene expression profile (72–74). As observed for expanded clones *in vivo*, LV-BACH2 cells at d25 show upregulation of genes that protect from CTL or NK cell-mediated killing, such as *HLA-E*, poliovirus receptor (*PVR*), *TNFRSF14* (encoding herpes virus entry mediator (HVEM)) and programmed death ligand 1 (*PDL-1*) (Fig4b). We also detected induction of immune checkpoint genes, similar to expanded clones *in vivo*, such as T-cell immunoreceptor with Ig and ITM domains (*TIGIT*), programmed cell death protein 1 (*PD1*) and B- and T-lymphocyte attenuator (*BTLA*) (Fig4b). Furthermore, BACH2 overexpression activates an intrinsic anti-apoptotic state in CD4+ T cells: as reported for expanded HIV-1 infected clones *in vivo*, LV-BACH2 cells were marked for example by upregulation of apoptotic inhibitor *BCL2L1* (encoding for BCL-XL protein), inhibitor of apoptosis (IAP) family member *BIRC5* and IAP regulator *USP19* (deubiquitinase that stabilizes IAP proteins) (Fig4c). Thus, these data indicate that BACH2-overexpression drives a CD4+ T cell phenotype that overall mimics the clonal expansion immune signature that is selected for in PLWH.

### 4. Aberrant expression of the STAT5B RdIG induces an activated, pro-survival CD4+ T cell state

While *BACH2* is the most prominent RdIG in PLWH on ART, a selection of further proviral integration genes have been proposed to be subject to insertional activation that could be driving cellular expansion (20,25,26,29–31,75). We asked whether insertional activation of these genes by proviral promoter exaptation could redirect CD4+ T cell fate in a similar way as here-demonstrated for BACH2. <u>S</u>ignal transducer and <u>a</u>ctivator of transcription 5B (*STAT5B*) is - like *BACH2* - a highly selected gene for proviral integrations in PLWH on ART (SuppFig1) (26,29–31). STAT5B is a central transcription factor in CD4+ T cells that regulates homeostasis, survival and differentiation (76–79), and has been associated with CD4+ T cell differentiation into Th1, Th2, Th9 and Treg cells (80). In PLWH the *STAT5B* RdIG shows significant clustering bias towards congruent proviral integrations in intron 1 (Fig5a) (26,29) and chimeric provirus/*STAT5B* transcripts (spliced into *STAT5B* exon two which contains the start of the open reading frame) have been detected in PBMCs of PLWH on ART (42), providing evidence for provirus promoter exaptation at this locus. Indeed, using a cell line clone transduced with a near full length HIV reporter that has integrated in transcriptional congruent orientation within intron one of the *STAT5B* gene locus, i.e. the region selected for proviral integrations in PLWH on ART, we observe that activation of LTR transcriptional activity though treatment with PMA/Iono resulted in generation of LTR-STAT5B chimeric transcripts, thereby raising overall levels of STAT5B-coding transcripts (SuppFig6). These results mimic above-described findings on HIV_BACH2 cell models and provide further evidence for HIV provirus-triggered insertional activation of *STAT5B* gene expression.

**Figure 5.**
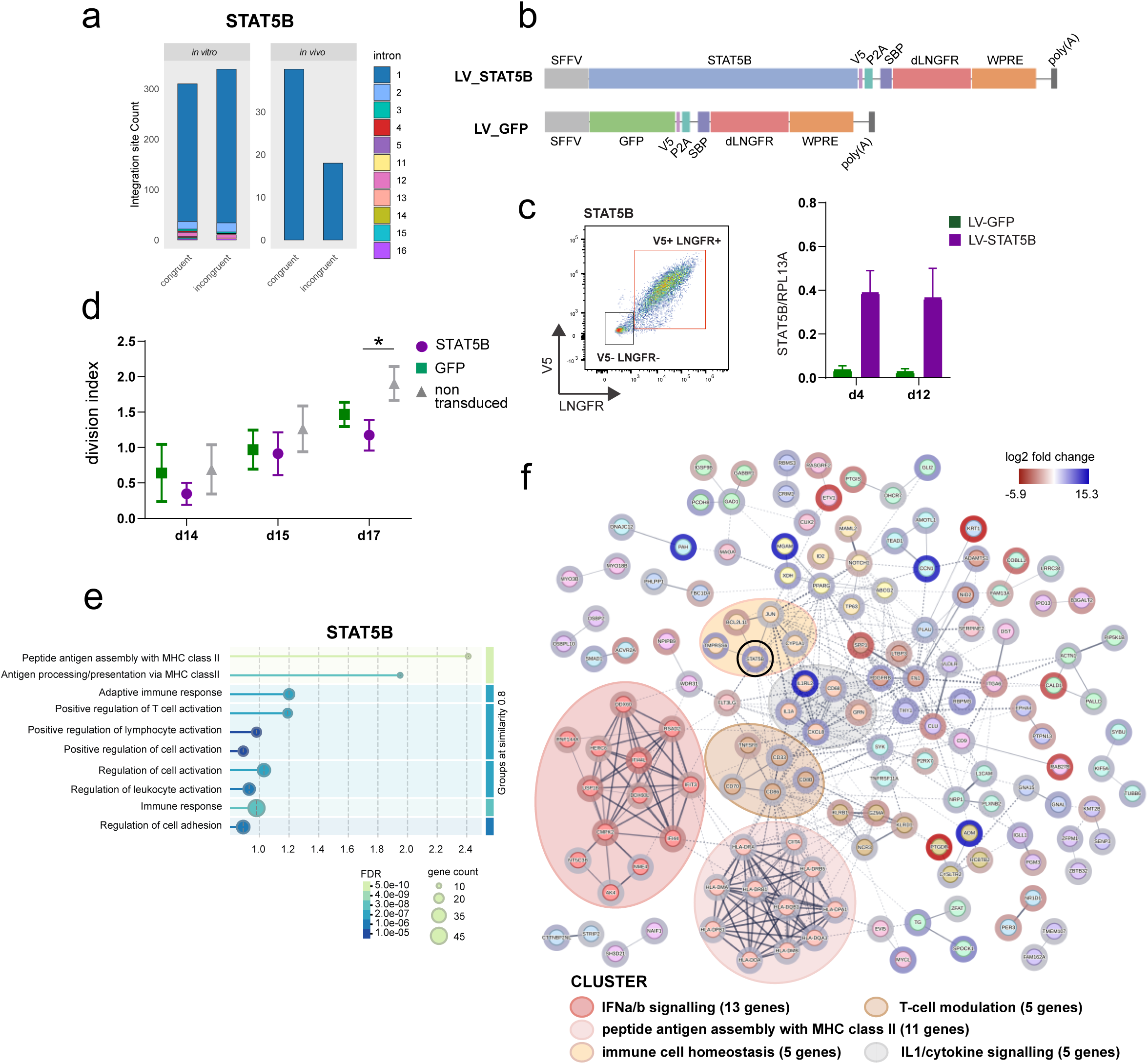
Aberrant expression of STAT5B alters CD4+ T cell phenoytpe towards effector type. (a) Frequency of in vivo and in vitro provirus integration sites in the STAT5B gene locus stratified by intron and transcriptional orientation of the provirus in context of the BACH2 gene (congruent, incongruent). (b) Schematic of LV-STAT5B and LV-GFP expression cassettes used to generate lentiviruses for transduction of primary CD4+ T cells to mimic aberrant expression. (c) Flow cytometric analysis of LV-STAT5B transduced CD4+ T cells stained for V5 and LNGFR expression (left panel). RT-qPCR for normalized STAT5B transcript levels in SPB-dLNGFR-enriched LV-STAT5B and LV-GFP transduced cells, at day (d) 4 and 12 post transduction (right panel). (d) Cell tracer-based assay for proliferative behavior of LV-STAT5B transduced cells as well as LV-GFP and non-transduced control cells. Cells were stained at d12 post transfection and followed on d14, d15 and d17. Statistical analysis was done using unpaired t-test, *p<0.05. (e) Enrichment analysis of functional gene networks stratified by biological processes based on DEG of LV-STAT5B transduced cells compared to LV-GFP controls at d12 post transduction. (f) STRING analysis based on DEG between LV-STAT5B and LV-GFP transduced CD4+ T cells at d12 post transduction.

We went on to address whether exaptation-driven, aberrant STAT5B activation could redirect CD4+ T cell fate in a similar way as BACH2. To do so, we used above-described overexpression assay in primary CD4+ T cells. We transduced activated CD4+ T cells from HIV-seronegative donors with lentiviral SBP-deltaLNGFR bi-cistronic expression construct for STAT5B (LV-STAT5B) (Fig5b). LV-STAT5B cells showed robust expression of STAT5B at RNA and protein level (Fig5c).

Contrary to BACH2, aberrant STAT5B expression did not alter proliferation rates of CD4+ lymphocytes in comparison to controls in a marker dye-based assay (Fig5d). Further, transcriptomic profiling and DEG analysis at 12 day post transduction revealed that STAT5B overexpression upregulated 222 and downregulated 145 genes (SuppTable1). Differing from LV-BACH2-treated cells, these transcriptomic changes positively highlighted pathways of cell activation, immune response and MHC class II antigen presentation as shown by GSEA and STRING analysis (Fig5e, 5f). Functional networks promoting inflammatory signaling and cell survival were also found enriched (Fig5f). Taken together, aberrant STAT5B expression promotes an activated CD4+ T cell state with pro-survival signature. This contrasts BACH2-driven CD4+ T cell reprogramming and suggests that insertional activation of RdIGs through LTR exaptation likely results in clones with different T cell states depending on the proviral integration site.

### 5. A negative feedback loop autoregulates endogenous *BACH2* expression

Our results indicate that an aberrant increase in *BACH2* expression leads to changes in CD4+ T cell state and drives longevity and cell proliferation. Our data also suggest that transcriptional activity of BACH2-integrated proviruses can cause such aberrant, LTR-driven BACH2 expression. Since BACH2, outside the HIV infection context, has to our knowledge however not been associated with T cell transformation or malignant expansion - as might be assumed given the robust proliferative phenotype - we asked whether there might be differences in the regulation of endogenous versus LTR-driven BACH2 expression. In particular we were interested in a potential autoregulatory feedback loop for endogenous BACH2 that had been previously hypothesized (43). We undertook investigation of the *BACH2* promoter region to gain further understanding. Motif scanning of a 40kb region around *BACH2* exon1 identified four T-MARE consensus motifs (TGCTGA(G/C)TCA; Fig6a), which are motifs that have been found to be targeted by BACH2 (81). To understand whether these motifs could have functional relevance, we assessed their epigenetic signature in CD4+ T cells. We found that two motifs in particular had an accessible chromatin profile, enriched for transcriptional regulatory histone marks, namely trimethylation of H3K4 and acetylation of H3K27. Furthermore, these two motifs have been found by chromatin immunoprecipitation to be bound by endogenous BACH2 in B cells (Fig6a). We therefore hypothesized that endogenous *BACH2* expression could be subject to autoregulation through promoter-inherent BACH2 binding sites.

**Figure 6.**
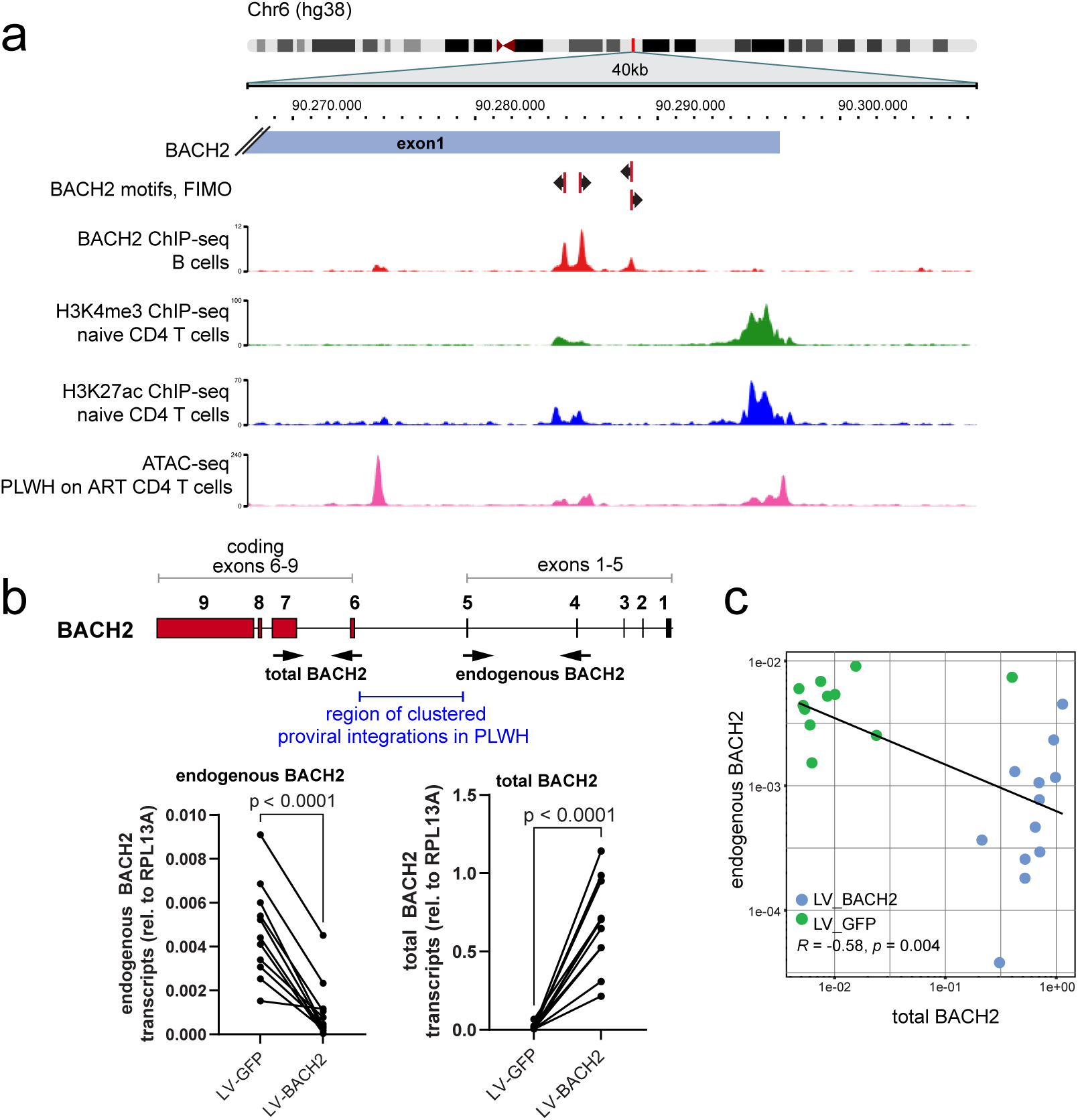
Autoregulation of BACH2 expression through negative feedback loop. (a) BACH2 exon 1 contains four BACH2 binding motifs, of which two are bound by BACH2 (data from B cells) and display cis-regulatory epigenetic marks and open chromatin accessible in CD4+ T cells. (b) Overexpression of BACH2 mimicking LTR-driven insertional BACH2 activation downregulates activity of the endogenous BACH2 gene promoter promoter. RT-qPCR of endogenous and total BACH2 transcript levels in LV-BACH2 and LV-GFP transduced CD4+ T cells. Total BACH2 transcripts derive from endogenous BACH2 locus and lentiviral overexpression construct (containing coding exons marked in red). Positions of primers used for RT-qPCR analysis are marked by arrows. Statistical analysis with paired two-tailed t-test. (c) Pearson correlation analysis of endogenous and total BACH2 transcript levels in LV-BACH2 and LV-GFP transduced CD4+ T cells (R2 and p values indicated in the plot).

To investigate whether BACH2, that has been primarily described as transcriptional repressor (82–84), could restrict its own expression, we took advantage of our aberrant BACH2 expression system in primary CD4+ T cells that have been transduced with LV-BACH2. We assessed whether transcriptional activity from the endogenous *BACH2* locus could be affected by raised BACH2 protein levels. To do so, we quantified *BACH2* transcripts that originated from the non-coding region in exons 1 to 5 and reflect endogenous *BACH2* transcriptional activity, in presence or absence of lentiviral-driven BACH2. Notably the BACH2 lentiviral overexpression constructs only contains coding *BACH2* sequences. Indeed, primary CD4+ T cells that were transduced with lentiviral LV-BACH2 demonstrated significantly reduced transcription of the endogenous *BACH2* locus compared to controls (Fig6b). Total and endogenous BACH2 transcript levels were negatively correlated (Fig6c). These data suggest that BACH2 expression levels are usually restricted through a negative autoregulatory feedback loop likely involving binding of BACH2 to its own promoter region. Outside the HIV-1 infection context, this feedback loop could thus prohibit a continued raise of BACH2 levels, and prevent reprogramming toward the proliferative CD4+ T cell phenotype that we observe upon overexpression. On the other hand, in PLWH on ART the prominent selection of cells harboring *BACH2* proviral integrants could thus be a result of LTR-driven aberrant *BACH2* expression that escapes endogenous regulatory mechanisms.

### 6. Aberrant *BACH2* expression through LTR exaptation can be controlled by HIV-1 latency promoting agents

Our data suggest that LTR exaptation can affect CD4+ T cell profiles, which may contribute to immune modulation and dysregulation observed in PLWH on ART. The immune modulatory activity is dependent on promoter activity of the integrated HIV LTR. To determine whether agents that interfere with HIV transcriptional activity can halt exaptation-dependent effects on the integration site locus, we tested how latency promoting agents (LPAs) impact on LTR-driven *BACH2* expression within our HIV_BACH2 cellular model. We focused on two previously reported LPAs, Spironolactone, an aldosterone receptor antagonist that has been shown to interfere with proviral transcription through degrading components of the TFIIH transcription factor complex (85,86); and KRIBB11, a heat shock factor 1-pTEFb complex inhibitor that has been shown to inhibit HIV-1 latency reversal (87). HIV_BACH2^D5/E10^ cells were pre-treated with Spironolactone, Kribb11, or DMSO control and then stimulated with PMA/Ionomycin to induce transcription of the *BACH2* integrated proviral reporter. Samples were processed for flow cytometry to detect reporter expression and quantitative RT-PCR for LTR/BACH2 chimeric transcript detection. Both LPAs reduced expression of the HIV_BACH2 bfp reporter (Fig7a). In addition, both LPAs also significantly reduced induction of the chimeric LTR/BACH2 transcripts (Fig7b). Thus, treatment with LPAs is able to reduce LTR exaptation activity and block insertional activation of provirus integration genes. These findings show that supplementation of ART treatment with LPAs could provide a means to avoid immune modulation and clonal expansion driven by transcriptionally active provirus sequences.

**Figure 7.**
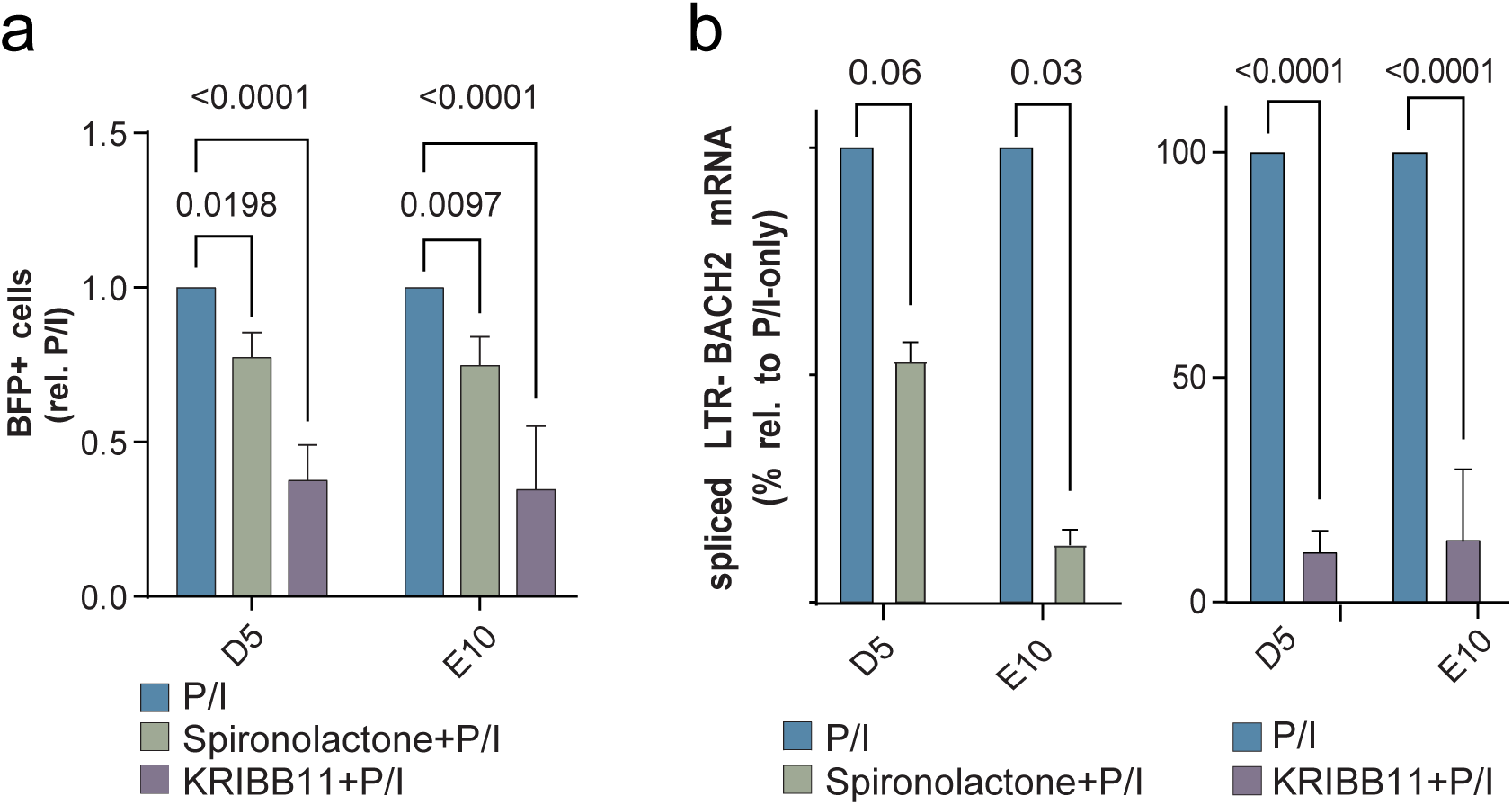
Latency promoting agents reduce LTR exaptation in HIV_BACH2 cells. HIV_BACH2^D5/E10^ cells were treated for 24h with LPAs Spironolactone or KRIBB11 and co-induced for 6hs with PMA/Ionomycine (P/I) prior to readout. Controls were treated with 6h P/I alone. (a) Fraction of cells that express the LTR-driven bfp reporter as measured by flow cytometry (relative to P/I only control). (b) Transcript levels of LTR/BACH2 chimeric transcripts as measured by RT-qPCR relative to P/I only control. Statistics using paired, two-sided t-test.

## Discussion

Expanded clones of HIV-1-infected CD4+ T cells that carry mainly defective proviruses are a prominent finding in PLWH on ART, their contribution to immune modulation in persistent HIV-1 infection is however unclear. In the present study we have used model systems to investigate the most abundant clones so far detected in PLWH, that is CD4+ T cells with proviral integrations in the *BACH2* locus. These cells are selected during ART with a proviral integration pattern that suggests dependence on proviral/host gene transcriptional crosstalk. We report that transcriptionally active, *BACH2*-integrated proviruses can - via LTR exaptation and evasion of autoregulatory feedback - aberrantly raise protein levels of BACH2, leading to induction of a proliferative precursor memory-like state that presumably persists through intrinsic anti-apoptotic activity and shielding against immune-mediated cell clearance. Interference with provirus transcriptional activity through LPA exposure can limit exaptation and thus provides a potential strategy to restrain BACH2 provirus clones. Beyond elucidating mechanisms into how BACH2 provirus clones may persist and influence the immune landscape in PLWH, our study more broadly underscores that the transcriptionally active, structurally defective provirus reservoir is a significant immune modulatory component during suppressive therapy.

Two hypotheses have so far been put forward for cells carrying *BACH2* integrated proviruses in PLWH on ART: First, *BACH2* proviral clones have been proposed to contribute to the replication-competent HIV-1 reservoir (42). However, since most datasets of proviral integrations do not correlate integration site with proviral sequence, it is unclear to date whether *BACH2* proviruses can contribute to persistence of the replication competent HIV-1 reservoir. To our knowledge, six studies have so far reported near full-length sequence information on a total of 14 BACH2 proviral integrants in PBMCs of PLWH on ART (12,23,45–48). In all but two reported cases, integration had occurred congruently, and in all cases, proviruses integrated in the *BACH2* locus carried sequence deletions of proviral genes that imply replication incompetence. Notably, these proviruses did however retain the HIV major splice donor site, which facilitates generation of chimeric LTR-BACH2 transcripts. The low number of *BACH2* integrants with sequence information preclude statistical analysis as for now, but there is currently no indication that *BACH2* clones contribute to the replication competent HIV-1 reservoir.

Second, it has been suggested that cells with *BACH2* integrated proviruses differentiate towards an activated, T-reg like phenotype. This conclusion was based on an *in vitro* model with targeted insertion of a minimal LTR/MSD cassette into *BACH2* (43). In contrast, our data suggest that BACH2-provirus clones do not present effector type-signatures but adopt a long-lived, precursor memory-like fate. Discrepancies could be due to differences in technical approaches (LV overexpression vs HDR manipulation) and time points of analysis (9 to 25 days post overexpression vs 7 weeks post HDR manipulation). In support of our data, BACH2 transcription factor has indeed previously been described to restrain differentiation towards CD4+ effector fates while promoting long-lived memory phenotypes (39,60,88,89). In agreement, two recent reports suggest that BACH2 acts as master regulator in directing CD4+ T cells towards a long-lived memory fate, supporting a cell population essential for HIV-1 persistence (40,41). As our study strongly suggests, this same pathway drives persistence of BACH2-proviral clones through provirus-driven BACH2 dysregulation. Mechanistically, we observe BACH2-induced transcriptomic changes to be gradual, with reprogramming of T cells taking place over several weeks in our experimental set-up. This may reflect that aberrant BACH2-expression indirectly leads to cell fate alteration. This is consistent with the finding, that BACH2 does not actively instruct expression of specific target genes, but rather acts as transcriptional repressor to limit activity of super enhancer (SE) genomic regions, which regulate genes important for T-cell identity and lineage-specific cytokine responses (90–92).

While clonal expansion is a well-described and prominent feature of the HIV-1 reservoir, we currently have very little understanding of how these clones modulate immune responses in PLWH and impact HIV persistence or infection-associated co-morbidities – in particular if clones carry structurally defective and replication-incompetent proviruses such as in the vast majority of cases. As for BACH2-proviral clones, our findings indicate that cells reprogrammed by LTR exaptation and thus aberrant BACH2 levels show upregulated expression of negative immune regulators, such as HLA-E. HLA-E has been shown to inhibit natural killer (NK) cell function through binding lectin-like receptor NKG2A (93,94). NK cell activity is thought to be important in controlling HIV-1 infection and HLA-E expression positively correlates with HIV viremia in PLWH (95). It has also been suggested that prolonged exposure to elevated HLA-E levels could cause NK cell dysfunction and reduced viral control *in vivo* (96). Thus, expansion of T cell clones with *BACH2* proviral integrants, could promote an immune environment in PLWH that is permissive to HIV persistence through loss of NK cell control. Our data also imply that proviral clones with STAT5B integrants, another prominently selected population in PLWH on ART, adopt an activated, inflammatory CD4+ T cell fate. Since inflammation can promote HIV-1 reactivation (97), STAT5B provirus clones could thus also contribute to an immune microenvironment that fosters not only pathogenic inflammation but also maintenance of the HIV-1 reservoir. Our findings provide first indications of different pathways how provirus exaptation-driven, reprogrammed infected CD4+ T cells might shape the HIV-1 reservoir.

Reprogramming of infected CD4+ T cells via exaptation of provirus regulatory elements is dependent on transcriptional activity of the provirus LTR. While proviruses in PLWH on ART have long time been thought of as primarily latent and transcriptionally silent, recent evidence suggest the contrary: Elongated viral transcripts have been detected in over 30% of all infected cells in PLWH, with evidence of LTR transcriptional activity in over 80% of these cells (16,17). This raises the question of potential functional consequences, in particular for active replication-defective proviruses. Our study highlights how these latter can impact host function via LTR exaptation, which can occur as long as LTR-inherent regulatory elements and splice donor sites are functional. Notably this goes beyond so far suggested immune modulatory activity of defective proviruses linked to viral gene transcripts or proteins (9–11,98). Given a recent report, which indicates that insertional activation of proviral integration genes is observed in up to 5% of HIV-1 infection events (99), future work is clearly warranted to further our understanding of how LTR exaptation shapes the immune landscape in PLWH. In this context, our findings also underscore the potential benefit of supplementing current antiretroviral regimens with components that block provirus transcription. As we show, these drugs have the capacity to limit insertional activation and thus can interfere with downstream host cell modulation.

In summary, our data highlight that proviruses – at the BACH2 locus and likely also further recurrently-detected integration genes - have the ability reprogram infected cells and modulate the immune landscape of PLWH on ART through exaptation of virus-derived regulatory elements. We speculate that this could impact HIV-1 persistence and thus should be taken into account in view of optimizing current antiretroviral therapy regimens for healthy remission of PLWH.

## Materials and Methods

### Generation and analysis of provirus integration site datasets

Data on provirus integration sites was extracted from the Retrovirus Integration Database (https://rid.cancer.gov) (49). Datasets of integration sites were generated combining integration site data from 11 studies on in vivo integrations mapped in PBMC samples of PLWH on ART (12,30,31,33,34,46,50–54), and from 4 studies on in vitro integrations mapped in in vitro HIV-1 infected CD4+ T cells (26,51,55,56). The IS were annotated using the ‘closest’ subcommand of BEDTools v2.27.1. For gene annotation, the RefSeq GRCh37 Release 105.20220307 reference was used, selecting only the “RefSeq Select” isoform for each protein-coding transcript. For visualizing of genome-wide integration site frequency, Circos plots were generated using Circos Builder (v069-815) of Galaxy Server, based on human hg19 karyotype ideogram (100). Bar plots depicting integration site distribution with regards to intronic position and provirus orientation were generated using Ggplot2. The integration site (IS) frequency for both in vivo and in vitro datasets was calculated for each gene using the formula: IS_freq = (IS count per gene / total IS count) × 100. To compare the two datasets, a scatter plot was generated where the x-axis represents individual genes of interest, and the y-axis displays the log₁₀-transformed difference in IS frequency between in vivo and in vitro.

### Genome-wide association study analysis

Genome-wide association study (GWAS) data including SNP location and trait information in context with *BACH2* was downloaded from https://www.ebi.ac.uk/gwas/genes/BACH2 (accessed May 2024). SNPs in *BACH2* were visualized in JBrowser2 (101). Trait information was categorized, counted and plotted using Microsoft Excel.

### Cell culture

HAP1 cells (Horizon Discovery) were used for generation of HIV_BACH2 reporter lines and cultured in IMDM medium (Thermo Fisher Scientific) with 10% FBS and 1% Pen/Strep. MOLT4 cells (incl HIV_STAT5B cells) were cultured in X-VIVO^TM^ 15 cell medium (Lonza) with 10% FCS and 0,5% Pen/Strep. Buffy coats for PBMC isolation were obtained from anonymized healthy donors through collaboration with the Institute of Transfusion Medicine, University Medical Center Hamburg-Eppendorf. PBMCs were isolated using density gradient centrifugation. CD4+ T cells were then isolated using the EasySep™ Human CD4+ T Cell Isolation Kit (stem cell technologies). CD4+ T cells were cultured in X-VIVO^TM^ 15 cell medium (Lonza) with 10% FCS, 0,5% Pen/Strep and 200U/mL IL-2 at a density of 1x10^^6^ cells/ml. Upon isolation, cells were activated with CD3/CD28 (ImmunoCult™ Human CD3/CD28 T Cell Activator, Stemcell^TM^).

### Generation of HIV_BACH2 reporter lines

An HIV-1 reporter construct was cloned in pMK backbone (Thermo Fischer Scientific) using sequential Gibson assembly (NEBuilder HiFi DNA assembly). 5’LTR, gag leader region and 3’LTR were derived from pNL4-3 (BEI Resources), bfp 2A sequence was amplified from pNLT2DenvBLB (102). To generate the transfer plasmid for CRISPR/Cas9-mediated knock-in, 5’ and 3’ homology arms of 1000bp length were amplified from genomic DNA extracted from Jurkat cell line and cloned flanking the HIV reporter. An 8bp long barcode sequence was introduced upstream the 5’LTR. Guide RNA for CRISPR/Cas9 targeting of the BACH2_3710 locus (5’-ggagattgtgtgaacccagc-3’) was cloned into pX330-U6-Chimeric_BB-cBh-hSpCas9 (Addgene #42230) utilizing BbsI restriction sites (yielding pX330-U6-Chimeric_3710_BACH2-cBh-hSpCas9). E-CRISP tool was used for gRNA design (http://www.e-crisp.org).

CRIPSR/Cas9-mediated insertion of the HIV reporter into BACH2 intron 5 in HAP1 cells was carried out following a workflow previously described (103,104). In brief, HAP1 cells were co-transfected using Lipofectamine 3000 (Thermo Fisher Scientific) with transfer plasmid and pX330-U6-Chimeric_3710_BACH2-cBh-hSpCas9 plasmid. Transfected cells were plated in limiting dilution for single cell clone generation at a density of 0.5-1 cells/well in flat bottom 96-well format. Screening for targeted clones was carried out in two rounds. First, bfp-expressing clones were selected after induction with 50 ng/ml phorbol 12-myristate 13-acetate (PMA; Sigma Aldrich) and 1 µM Ionomycin (Sigma Aldrich) for 18 hours. Positive clones based on flow cytometry were subsequently screened by PCR on whole cell lysate for presence genomic insertion of the reporter at the targeted locus. To exclude cell clones with an integrated vector backbone, clones were screened by PCR for presence of pMK backbone (negative selection). Site-specific homozygous or heterozygous BACH2 targeting events were assessed by Southern blotting using bfp-specific and genomic probes as described previously (103). In brief, high molecular weight genomic DNA from 3710_BACH2_positive clones was extracted from whole cell lysate by isopropanol precipitation, digested with *ExoRI* and blotted onto Amersham Hybond^TM^-N^+^ blotting membrane (GE Healthcare). The corresponding probes were PCR amplified from genomic DNA or targeting plasmid DNA and labelled with ^32^dCTP using High Prime DNA labeling (Roche). Membranes were hybridized with labelled probes in hybridization buffer at 65°C over night. After stringent washing, membranes were exposed to Amersham Hyperfilm^TM^ MP (GE Healthcare).

### Generation of HIV_STAT5B reporter line

Clones were generated by limiting dilution following transduction of Molt-4 cells with pV1/SBP-P2A-GFP (105), a V1-derived minimal reporter virus carrying authentic NL43 proviral 5ʹ and 3ʹ sequences, at an MOI of 0.3. Successful proviral integration in each single-cell clone was confirmed by amplification of host–virus junctions using ligation-mediated PCR(46), followed by visualization of PCR products via agarose gel electrophoresis. Integration sites of positive clones were determined by sequencing of LM-PCR products. The HIV_STAT5B single-cell clone was selected based on its integration site in STAT5B.

### PMA/Iono and SPA treatment

For PMA/Iono treatment, cells were seeded at 1 x 10^6^ cells/ml and treated with 50 ng/ml phorbol 12-myristate 13-acetate (PMA; Sigma Aldrich) and 1 µM Ionomycin (Sigma Aldrich) or DMSO as a control. Reporter expression was analysed by flow cytometry 16 hours post treatment. The effect of silencing promoting agents (SPAs) KRIBB11 and Spironolactone was assayed in HIV_BACH2 cells as follows: 1 x 10^6^ cells/6-well were seeded and simultaneously treated with KRIBB11 (final concentration 5 uM, Absource Diagnostic) or Spironolactone (final concentration 10 uM, Sigma-Aldrich). 18 h post seeding, cells were induced with PMA/Iono. One day post seeding RNAzol samples were collected and reporter(48) expression was analyzed by flow cytometry.

### Western Blot analysis of HIV_BACH2 cells

HIV_BACH2 cells were activated with PMA and Ionomycin for 16h as described before. Cells were lysed in RIPA buffer (150 mM NaCl, 1 % Triton X-100, 0.25 % NP40, 0.1% SDS, 50 mM Tris HCl pH8.0 in H2O) supplemented with oComplete protease inhibitor cocktail (Millipore) and protein concentration determined by Pierce 660 reagent according to manufacturer’s instructions (Thermo Fisher). Twenty ug of total protein were separated by SDS-PAGE and transferred to nitrocellulose membranes (Whatman Protran). Primary antibodies against BACH2 (clone D3T3G, Cell Signaling) and LaminA/C as control (sc-376248, Santa Cruz) were used in combination with secondary HRP-conjugated antibodies (Jackson Laboratories) for detection.

### SAM-mediated targeted transcriptional activation

SAM-mediated LTR (LTR-SAM) and BACH2 (BACH2-SAM) targeted transcriptional activation of HIV_BACH2 and wt cells was carried out as previously described (32). In brief, 5*10^5^ cells were seeded in a six-well plate and co-transfected with 1.4 µg sgRNA(MS2)-gRNA for either LTR or BACH2 activation, 0,8 µg dCas9-VP64_GFP (Addgene #61422) and 1 µg MS2-P65-HSF1_GFP (Addgene #61423) per well. Transfection was performed using TransIT LT1 reagent (Mirus Bio) according to the manufacturer’s protocol. Three days post transfection cells were harvested, washed with PBS, sorted for BFP/GFP-positive cells (FACS Aria, BD Biosiences), washed once more and stored in RNAzol (Sigma Aldrich) at - 80°C upon further processing. Total RNA was isolated following the RNAzol manufacturer’s protocol. All experiments were conducted in n=3 independent replicates.

### Production of lentivirus and transduction of CD4+ T cells

Lentiviruses were produced by plasmid co-transfection of the following plasmids into LentiX cells (Takara Bio) using TransIT-LT1 (Mirus Bio) according to manufacturer’s instructions: psPAX2 (Addgene #12260), pCMV-VSV-G (Addgene #8454) and lentiviral transfer vectors. Supernatants were collected 72 hours post transfection and filtered through a 0.22 µm filter (Millipore). Virus concentration was conducted by ultracentrifugation at 12,300 × g for 1.5h at 4 °C (Beckman Coulter) and resuspended in X-VIVO 15 medium (Lonza). CD4+ T cells from healthy donors were activated by CD3/CD28 stimulation 24 h prior transduction. Spin-inoculation was used to transduce the cells with lentivirus stocks in X-VIVO 15 medium (Lonza) containing 0.2 % protamine sulfate (Sigma) (850 x g, 1 h, 30 °C). Following centrifugation, the plates were incubated for 24 h at 37 °C before medium was replaced by X-VIVO 15 cell with 10 % FBS, 0.5 % Pen/Strep and 200 U/mL IL-2 (Biomol).

### Magnetic bead-mediated enrichment for SBP-LNGFR + cells

5x10^6^ transduced CD4+ T cells were enriched using magnetic selection of the SBP-ΔLNGFR surface marker with Dynabeads Biotin Binder (Invitrogen). In brief, cells were washed three times with incubation buffer (0.5 M EDTA, 1 mg/L BSA in PBS). Beads were added at a 10:1 ratio (beads:cells) and the mixture was incubated at 4 °C for 30 min. After incubation, the samples were placed on a magnetic stand for 3 minutes before carefully removing the supernatant. Bead-bound cells were washed with cold incubation buffer, placed into the magnetic stand again for 3 minutes before decanting the supernatant. Bead-bound cells were resuspended in pre-warmed release buffer (1 M HEPES, 20 mM biotin in medium) and incubated at room temperature for 15 min. The tube was placed on the magnetic stand for 3 minutes before collecting the released cells in the supernatant.

### Analysis of cell proliferation

Division index of transduced cells was measured using CellTrace™ Far Red Cell Proliferation Kit (Thermo Fisher Scientific) according to the manufacturer’s instructions. In brief, transduced cell populations and non-transduced control cells (n=3) were stained at d12 after transduction. Expression of the Far Red marker dye was measured at d12 and at three following time points (d14, d15 and d17 after transduction) by flow cytometry.

### Quantitative RT PCR assays

RNA was extracted using RNAzol (Sigma Aldrich) according to manufacturer’s instructions. cDNA was generated from 1ug total RNA using SuperScript III Reverse Transcriptase (Invitrogen). Droplet digital PCR with the Bio-Rad QX200 system was used for quantification of i) endogenous *BACH2* (amplicon spanning exon 4/5, upstream of region of recurrently-detected proviral integrations in PLWH) and total BACH2 (exon 6, downstream of region of recurrently-detected proviral integrations in PLWH) transcript levels; and ii) endogenous (exon 1+ exon 2), total (exon 10 + exon 11) and chimeric (LTR + exon 2) STAT5B transcript levels (utilized oligonucleotides are provided in SupplTable2). Each 20 µL reaction contained 10 µL of 2× ddPCR Supermix for probes (No dUTP) (Bio-Rad), 2 µL of each primer (10 µM stock), 1 µL probe (5 µM stock), 2 µL of cDNA template, 1µL of RPL13A primer/probe mastermix for normalization (Bio-Rad #10031255) and 2 µL nuclease-free water. The reaction mix was loaded into the QX200TM Droplet Generator (Bio-Rad) with 70 µL of Droplet Generation Oil, and droplets were generated according to the manufacturer’s guidelines. PCR amplification was performed using a T100TM Thermal Cycler (Bio-Rad) under the following cycling conditions: 95 °C for 10 min, 40 cycles of 94 °C for 30 sec and 57 °C for 1 min, 98 °C for 10 min, hold at 4 °C. Following amplification, droplets were analyzed using the QX200 Droplet Reader (Bio-Rad). Resulting data was visualized using GraphPad Prism v10 software.

### BACH2 binding site and chromatin landscape analysis

The datasets supporting the conclusions of this analysis are publicly available and can be accessed in NCBI repositories under the accession numbers GSE102460 [BACH2 ChIP-seq], GSE122826 [H3 ChIP-seq], and GSE144329 [PLWH-on-ART ATAC-seq]. To identify putative BACH2 binding sites across the human genome, the BACH2 motif was obtained from the JASPAR database (2024 expansion, ID: MA1101) (106). The hg38 human genome assembly was scanned for BACH2 motifs using FIMO, a tool within the MEME suite (v5.5.5) (107). FIMO was run using the MEME suite’s precomputed hg38 background model based on the genome’s nucleotide frequencies. Significant BACH2 motif hits were defined as those with *p* < 1e-4. Raw ChIP-seq and ATAC-seq data were first assessed for quality using FastQC (v0.12.1) (https://www.bioinformatics.babraham.ac.uk/projects/fastqc/). Subsequently, reads were trimmed with Trimmomatic (v0.39) removing adapter sequences and low-quality bases (108). Next, trimmed reads were aligned to the hg38 human reference genome using Bowtie2 (v2.5.1) with the --very-sensitive-local parameter for optimal sensitivity (109). All unmapped reads and reads with a mapping quality score below 10 were excluded from further analysis. PCR duplicates were removed using Picard MarkDuplicates (v2.26.10) (http://broadinstitute.github.io/picard). For peak calling, MACS2 (v2.2.7.1) was used with parameters tailored for each assay — broad peak calling was performed for H3K27ac ChIP to capture large acetylation domains, while narrow peak calling was used for BACH2 ChIP, H3K4me3 ChIP, and ATAC-seq open chromatin regions (110). Peak annotation to hg38 genes was carried out using Homer (v4.11) (https://homer.ucsd.edu/). Sample integration and signal normalization was performed using BEDTools (v2.29.2) and deepTools (v3.5.5) (111,112). Tracks were imported in bigWig format and visualized in JBrowse2 (v2.18.0) (113).

### RNA sequencing and RNAseq dataset analysis

RNA sequencing was carried out on total RNA, isolated from SAM-treated HAP1_BACH2_LTRbfp2Atat cells and LV-transduced CD4+ T lymphocytes. For HAP1 samples, 1 ug of RNA was subjected to library preparation utilizing the NEBNext Ultra II Directional RNA mRNA kit in combination with NEBNext Multiplex Oligos Unique Dual Index UMI Adaptors and sequenced at ∼100M clusters 2x100bp per sample (Illumina NextSeq500). For primary T lymphocytes, about 5 mio transduced CD4+ T cells were enriched using magnetic selection based on SBP-ΔLNGFR surface marker expression (see above). Total RNA was isolated from selected primary CD4 T cells using RNAzol (Sigma Aldrich) and following manufacturer’s instructions. Isolated RNA was subjected to library preparation using CORALL mRNA-Seq Library Prep Kit (Lexogen) and subsequent sequenced at ∼20M reads, 2x50bp per sample (Illumina NextSeq500).

Sequence reads were mapped using HISAT2 (HAP1 RNAseq) or Salmon (CD4 T lymphocyte RNAseq) aligners against the human reference genome (hg38) after adapter trimming and quality filtering using TrimGalore (114–116). Gene count matrices were further analysed in R in terms of differential gene expression (DEG) and gene set enrichment analysis (GSEA) utilizing packages DESeq2 (v1.44.0), EnhancedVolcano (v1.22.0), pheatmap (v1.0.12), ClusterProfiler (v4.12.6) and ggplot2 (117–121). In CD4+ T lymphocyte over expression experiments contrasting LV-GFP vs LV-BACH2 samples, ‘donor_ID’ was considered as covariate. In PCA plots this was achieved utilizing DESeq2 vst transformation of gene counts and limma::removeBatchEffect() (v3.60.3) for ‘donor_ID’ (119,122). Treeplots of significant gene ontology (GO) terms were generated by hierarchical clustering based on Jaccard’s similarity index using ClusterProfiler v(4.12.6) (120). Analysis of enrichment of functional gene networks using MCL clustering (inflation parameter 3) was done using STRING (v12.0) (123).

Single BACH2 exon analysis in the HAP1 reporters was conducted with a filtered genome annotation file (GTF) for BACH2 reference transcript ENST00000257749 containing 9 exons. HTSeq function htseq_count (v2.0.5) with parameters -m union -i exon_number -t exon was used to get read counts for individual BACH2 exons (124). Downstream analysis and plotting in R utilizing packages ggplot2 (v3.5.1) and ggpubr (v0.6.0) (121,125). Statistical method: unpaired t-test.

HIV-to-host transcript splicing in HAP1 reporters was analysed by inserting the HIV reporter sequence into the BACH2 locus (position chr6:90012553-90015110) of the reference genome version GRCh38. Genome annotations (GTF) were accordingly updated using reform (126). Subsequently raw FASTQ files were aligned to the updated reference genome and annotation using STAR (127). Splicing pattern plots were generated using the modified GTF annotation file and the sashimi_plot command of miso with scaling parameters for intron and exon at 50 and 3 respectively (128,129).

## Data sharing

### Materials availability

All new unique reagents generated in this study are available from the corresponding author with a completed materials transfer agreement.

### Datasets availability

The raw RNA-Seq data generated as part of this study have been uploaded in the GEO repository (accession #GSE304581) and are available upon request. The remaining data are available within the article or Supplementary Information. Any additional information required to reanalyze the data reported in this study is available from the corresponding author upon request. This study does not report original code.

## Supporting information

SFig1

SFig2

SFig3

SFig4

SFig5

SFig6

SuppTableLegends

STable1

STable2

## Acknowledgements

We would like to acknowledge technical assistance in the project from Christina Herrde, Susanne Hoibian and Manuela Kolster as well as the flow cytometry and sequencing core facilities at the Leibniz Institute of Virology (LIV). This work was supported by funds from the German Federal Ministry of Research, Technology and Space award number 01KI2105 (U.C.L.), NIAID award number UM1AI164559, co-funded by NHLBI, NIDA, NIMH, NINDS, and NIDDK (U.C.L., C.F., N.R.R.) and the German Center of Infection Research (DZIF) award number TI07.005 (U.C.L.).

## Author information

U.C.L. conceived and designed the project. C.F., N.R., E.T. and U.C.L. secured funding. C.S., M.V., F.M. and R.S. carried out cell line experiments. L.B. and M.K. carried out primary cell experiments. Computational data analysis was carried out by: M.S. and M.A. (integration site database), S.M.L. (promoter characterization), M.V.H., J.F., S.V. and A.G. (RNAseq). M.V.H., L.B. and U.C.L. interpreted the data and discussed the results with all authors. U.C.L. wrote the manuscript, in consultation with M.V.H., F.M., C.F., N.R.R. and E.T. with comments from all authors. M.V.H. and L.B. contributed equally to this work.

