## Supplementary material for "Selective persistence of HIV-1-infected T cell clones can occur through immune reprogramming driven by defective, transcriptionally active proviruses": SFig1

**SUPPLEMENTAL FIGURE 1**
**Hamann et al**

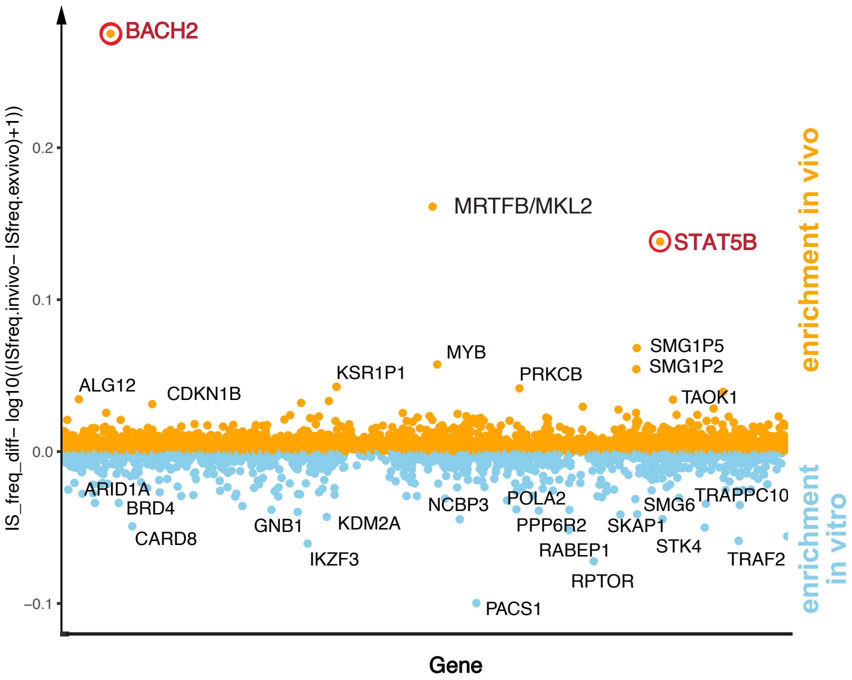


**Suupplemental Figure 1**
**Reshaping of provirus integration landscape in PLWH on ART.** Manhattan plot showing enrichment of proviral integration genes in PLWH on ART (*in vivo*) versus HIV-1 infection of CD4+ T cells *in vitro* as calculated by frequency of integration sites per gene locus. Data on proviral integration sites was extracted from Retrovirus Integration Database [45].
