## Supplementary material for "Selective persistence of HIV-1-infected T cell clones can occur through immune reprogramming driven by defective, transcriptionally active proviruses": SFig2

### SUPPLEMENTAL FIGURE 2
#### Hamann et al

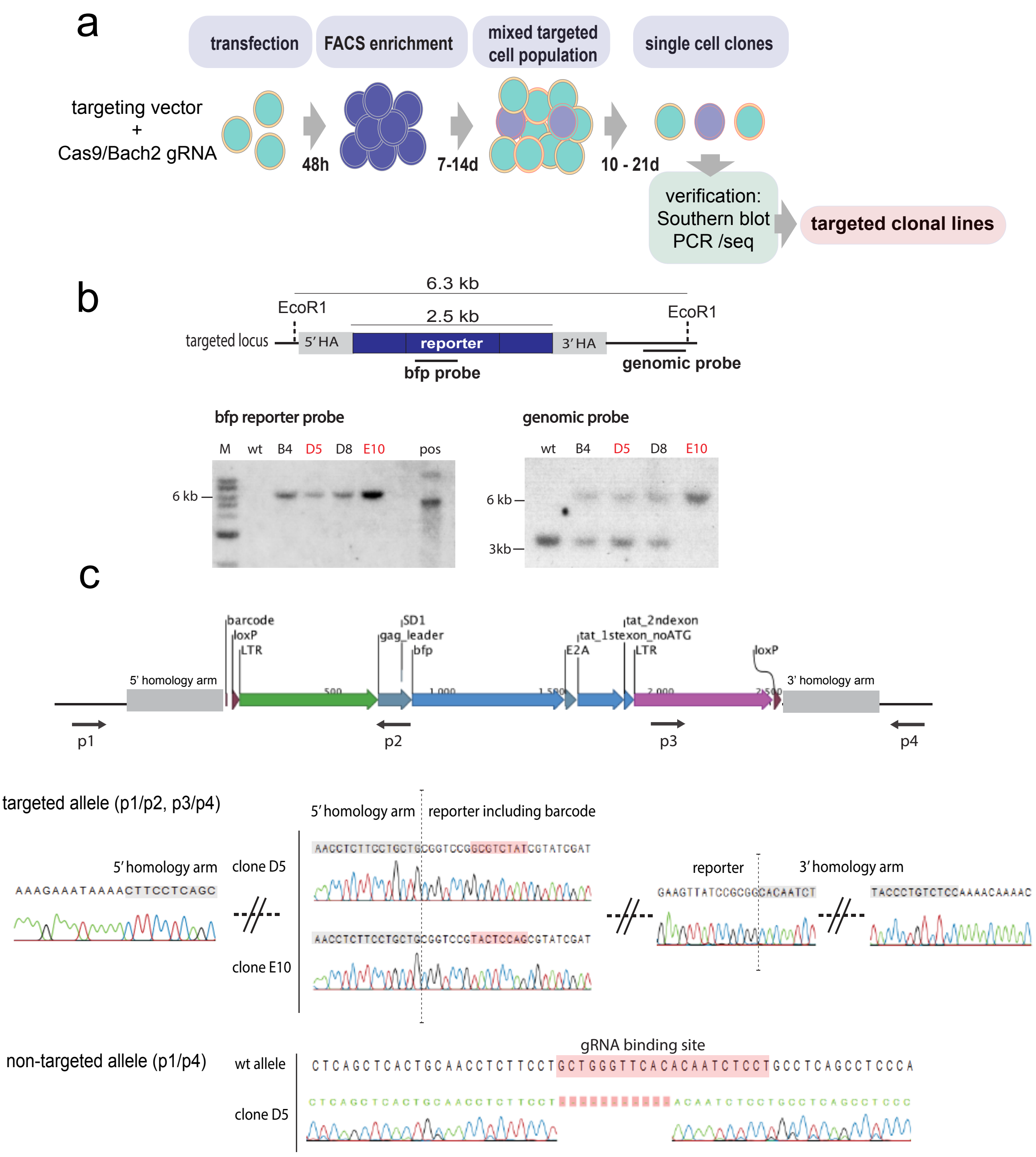


**Supplemental Figure 2**

**Generation of HIV_BACH2 cell line models. (a) Workflow for generating HIV_BACH2 clonal cell lines.** (b) Verification of HIV_BACH2 clonal lines for correct reporter targeting into the BACH2 locus (3710bp upstream of BACH2 open reading frame start) using Southern blotting. Genomic DNA of targeted clones was extracted, digested with EcoR1 and submitted to gel electrophoresis for size-separation of fragments. Blotted DNA was probed against bfp (bfp probe) and a genomic region outside the homology arm (HA) sequence employed for targeting (genomic probe). Clonal lines B4, D5, D8 and E10 were tested and showed heterozygous (B4, D5, D8) or homozygous (E10) pattern of targeted reporter insertion. Lines D5 and E10 were selected for further analysis. (c) Verification of HIV_BACH2 clonal lines for correct reporter targeting by Sanger sequencing. Genomic DNA of HIV_BACH2 D5 and E10 clonal lines was extracted and genomic regions containing 5' (primer p1 and p2) and 3' (p3 and p4) genome/reporter junctions were amplified by PCR and products submitted to Sanger sequencing. For D5 lines, the region spanning the non-targeted allele, containing the complementary sequence to the gRNA used for Cas9 targeting, was amplified using primers p1 and p4. Chromatograms show correct genome/reporter junctions and an 11bp deletion in the gRNA-complementary sequence on the non-targeted allele for D5 lines.
