## Supplementary material for "Selective persistence of HIV-1-infected T cell clones can occur through immune reprogramming driven by defective, transcriptionally active proviruses": SFig3

### SUPPLEMENTAL FIGURE 3

**Hamann et al**

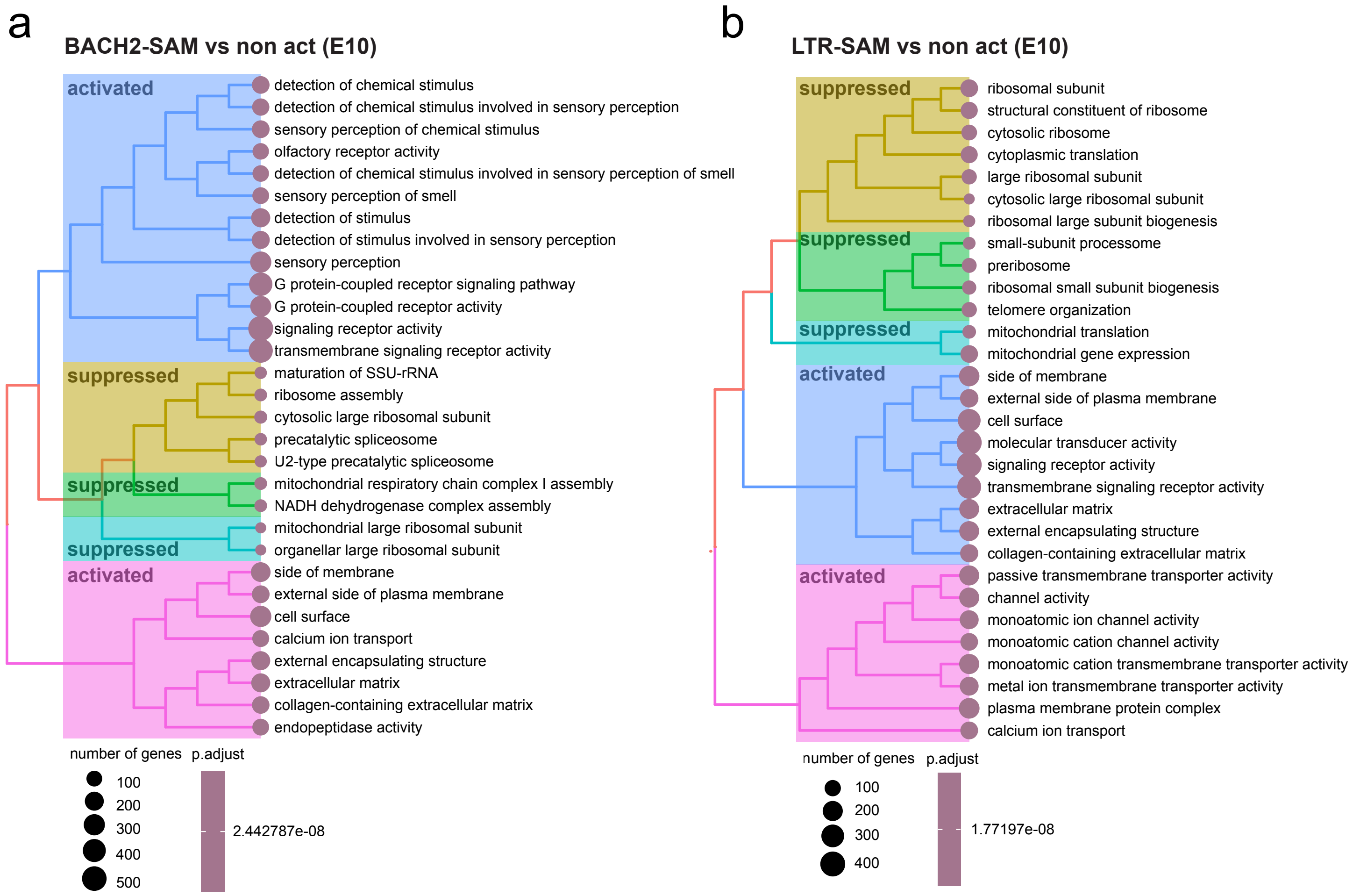


**Supplemental Figure 3**

**BACH2-induced transcriptomic changes in HIV_BACH2 cells.** HIV_BACH2$^{E10}$ clonal lines were submitted to BACH2-targeted (BACH2-SAM) or LTR-targeted (LTRT-SAM) transcriptional activation and transcriptomic changes analyzed by RNAseq. Treeplots depicting GSEA of differentially expressed genes upon (a) BACH2-targeted activation (BACH2-SAM) and (b) LTR-targeted activation (LTR-SAM) are shown.
