## Supplementary material for "Selective persistence of HIV-1-infected T cell clones can occur through immune reprogramming driven by defective, transcriptionally active proviruses": SFig4

### SUPPLEMENTAL FIGURE 4

#### Hamann et al

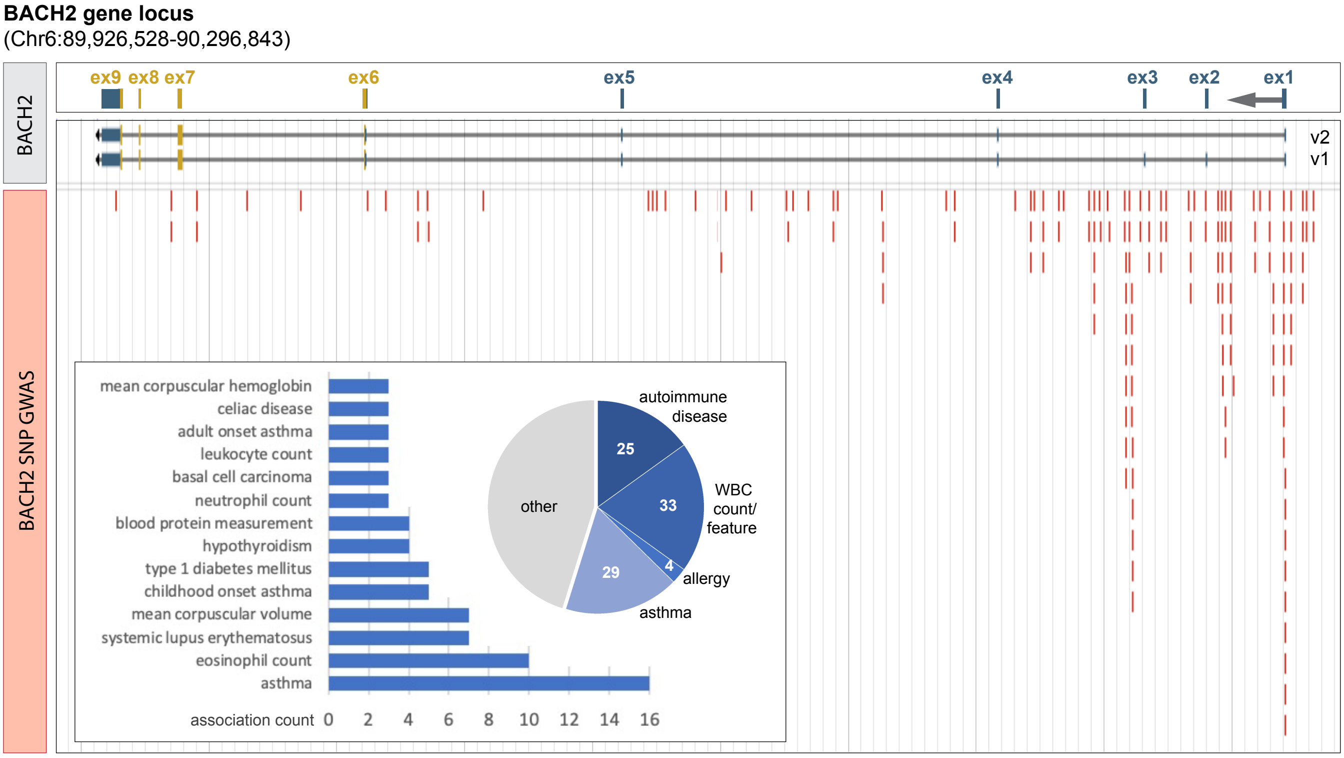


**Supplemental Figure 4**

**Single nucleotide polymorphisms (SNPs) in the BACH2 locus have been associated with immune-related traits in genome-wide association studies (GWAS).** The NHGRI-EBI GWAS catalog was used to retrieve information on BACH2 SNP-associated traits [131]. 84 EFO traits were associated with 166 SNPs within the BACH2 locus. Positions of SNPs relative to BACH2 gene and main BACH2 transcript variants (v1, v2) are marked (red bars). Exons coding for BACH2 open reading frame are highlighted in yellow. Bar graph insert depicts the 14 traits with association counts >3. Pie chart based on all BACH2 associations highlights associations with immune-related traits (blue).
