## Supplementary material for "Selective persistence of HIV-1-infected T cell clones can occur through immune reprogramming driven by defective, transcriptionally active proviruses": SFig5

### SUPPLEMENTAL FIGURE 5
#### Hamann et al

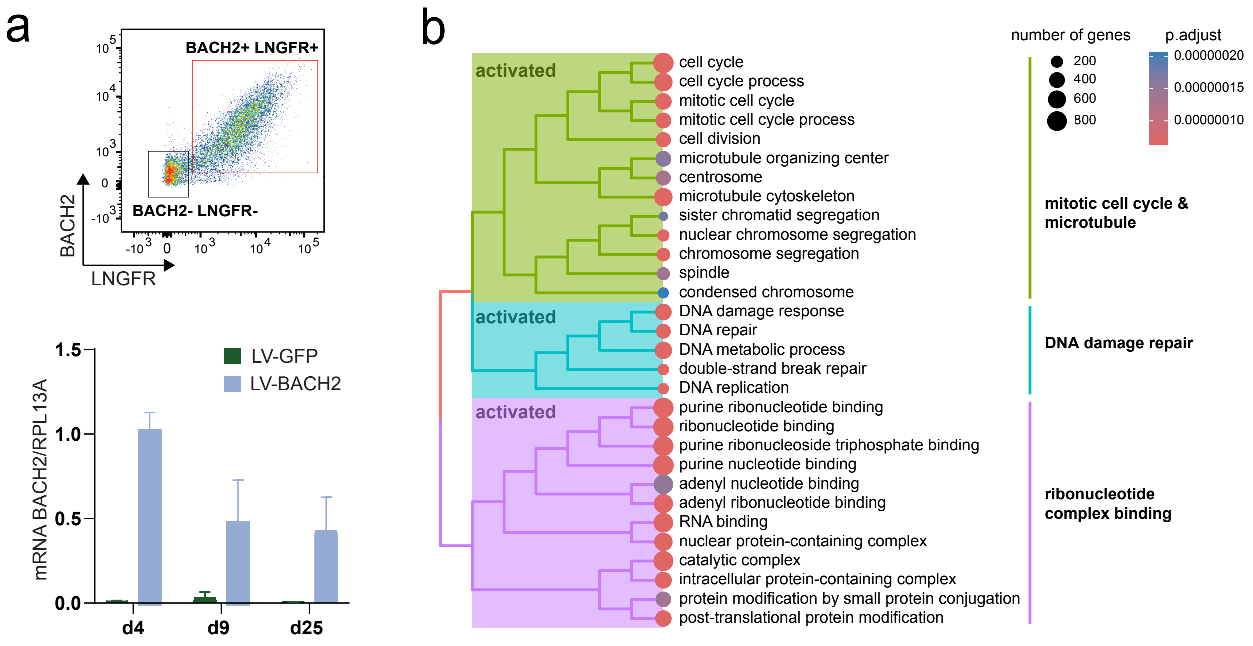


**Supplemental Figure 5**

**Transcriptomic changes induced by aberrant BACH2 expression in CD4+ T cells.** (a) Flow cytometric analysis of LV-BACH2 transduced CD4+ T cells stained for BACH2 and LNGFR expression (upper panel). RT-qPCR for normalized BACH2 transcript levels in SPB-dLNGFR-enriched LV-BACH2 and LV-GFP transduced cells at day (d) 4, 12, 25 post transduction (lower panel). (b) Gene set enrichment analysis (GSEA) based on DEG between LV-BACH2 and LV-GFP transduced CD4+ T cells at d25 post transduction delineates activated cellular pathways in LV_BACH2 cells.
