## Supplementary material for "Selective persistence of HIV-1-infected T cell clones can occur through immune reprogramming driven by defective, transcriptionally active proviruses": SFig6

### SUPPLEMENTAL FIGURE 6
#### Hamann et al

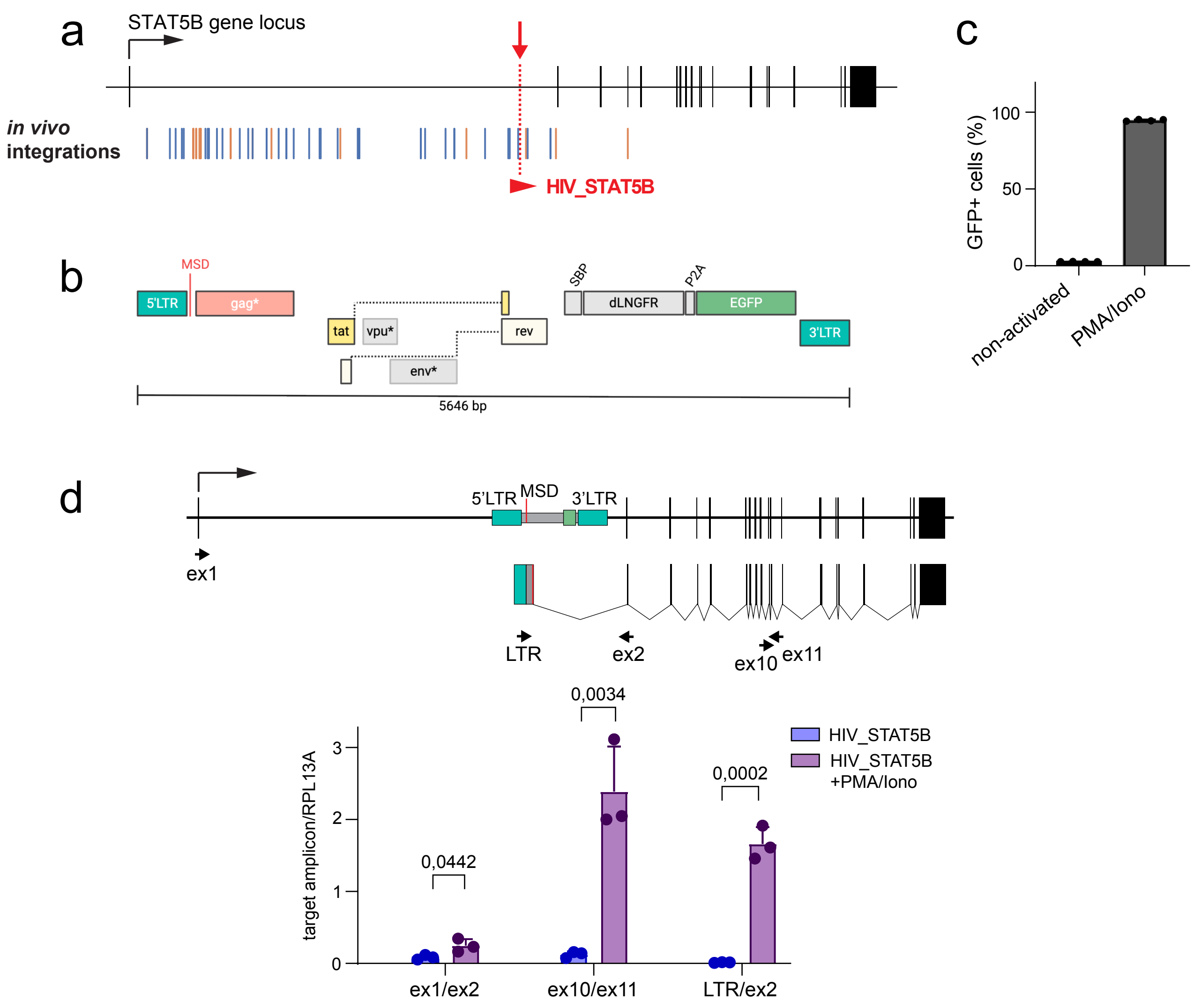


**Supplemental Figure 6**

**LTR-driven insertional activation of the STAT5B RdIG in HIV_STAT5B cells.** (a) Schematic of the STAT5B gene locus. Blue/orange bars indicate mapped provirus integrations in PBMCs of PLWH on ART (blue=congruent integration, orange= incongruent integration). The site of integration of the HIV-1 reporter in the HIV_STAT5B cell line within intron 1 4545bp upstream of STAT5B ATG is marked (red arrow). (b) Schematic of the HIV-1 reporter (V1/SBP-P2A-GFP) in the HIV_STAT5B cell line. MSD: major splice donor site; SBP: Streptavidin binding protein; dLNGFR: truncated low-affinity nerve growth factor receptor; *denotes presence of inactivating mutations. (c) The HIV-1 reporter in HIV_STAT5B cells is transcriptionally inactive but can be induced with PMA/Iono. Percentage of GFP positive HIV_STAT5B cells without and with exposure to PMA/Iono. (d) Detection of different STAT5B transcripts in HIV_STAT5B cells without and with PMA/Iono exposure by RT-qPCR. Schematic indicates the STAT5B locus in HIV_STAT5B cells (upper panel) and chimeric LTR-driven LTR/STAT5 transcripts generated by aberrant splicing from the reporter-inherent HIV-1 MSD to acceptor site of STAT5B exon 2. Arrows indicate position of primers used for target amplification of endogenous (ex1/ex2), chimeric (LTR/ex2) or total (ex10/ex11) STAT5B transcripts (arrowheads). PMA/Iono exposure induces LTR transcriptional activity leading to generation of chimeric LTR/ex2 transcripts that raise total STAT5B transcript levels. Statistics using unpaired, two-sided t-test.
