## Supplementary material for "Selective persistence of HIV-1-infected T cell clones can occur through immune reprogramming driven by defective, transcriptionally active proviruses": SuppTableLegends

Hamann et al.

### Supplemental Table legends

**Supplemental Table 1**

Differentially expressed genes (DEGs) from RNAseq assays of HIV_BACH2 SAM-treated cell lines and LV-BACH2/LV-STAT5B-treated CD4+ T cells (in comparison to controls).

**Supplemental Table 2**

Oligonucleotides used in this study.
