## Supplementary material for "Selective persistence of HIV-1-infected T cell clones can occur through immune reprogramming driven by defective, transcriptionally active proviruses": STable1

LV_BACH2_vs_GFP_d4

| baseMean | log2FoldChange | lfcSE | stat | pvalue | padj | ENSEMBL | SYMBOL |
|---|---|---|---|---|---|---|---|
| 132,3226787 | -1,34533462 | 0,332479653 | -4,046366766 | 5,20187E-05 | 0,034398565 | ENSG00000087842 | PIR |
| 31,97700974 | 3,924563643 | 0,970519567 | 4,043775907 | 5,25972E-05 | 0,034398565 | ENSG00000284294 | LOC122455341 |
| 83,31786793 | -1,638653025 | 0,398985997 | -4,107043948 | 4,00755E-05 | 0,030577607 | ENSG00000100292 | HMOX1 |
| 64,14997557 | -2,034695066 | 0,488884041 | -4,161917541 | 3,15586E-05 | 0,025495662 | ENSG00000185304 | RGPD2 |
| 198,5645871 | -1,184873724 | 0,276210478 | -4,28974937 | 1,78875E-05 | 0,015354171 | ENSG00000126822 | PLEKHG3 |
| 169,4540464 | 1,605085215 | 0,371454882 | 4,32107718 | 1,55269E-05 | 0,014216461 | ENSG00000015568 | RGPD5 |
| 115,9304322 | 1,510924286 | 0,342193726 | 4,415406163 | 1,00821E-05 | 0,010651299 | ENSG00000131401 | ENSG00000131401 |
| 330,8513753 | -1,114017698 | 0,238575398 | -4,669457572 | 3,01996E-06 | 0,00368999 | ENSG00000205336 | ADGRG1 |
| 65,16185864 | 2,244297214 | 0,479129466 | 4,684114367 | 2,81173E-06 | 0,00368999 | ENSG00000075391 | RASAL2 |
| 325,519042 | 1,060407617 | 0,223306617 | 4,748661862 | 2,04767E-06 | 0,003515337 | ENSG00000134852 | CLOCK |
| 322,7546624 | 1,224972866 | 0,247659867 | 4,946190433 | 7,56799E-07 | 0,001732314 | ENSG00000170100 | ZNF778 |
| 232,5551839 | 1,483782351 | 0,2913395 | 5,092966634 | 3,52504E-07 | 0,000968257 | ENSG00000162614 | NEXN |
| 174,2403185 | -1,521411869 | 0,283946313 | -5,358096941 | 8,41031E-08 | 0,000552605 | ENSG00000138678 | GPAT3 |
| 11238,08548 | 6,645958138 | 0,149218337 | 44,53848148 | 0 | 0 | ENSG00000112182 | BACH2 |

LV_BACH2_vs_GFP_d9

| baseMean | log2FoldChange | lfcSE | stat | pvalue | padj | ENSEMBL | SYMBOL |
|---|---|---|---|---|---|---|---|
| 44,13857273 | 18,71703136 | 3,456370371 | 5,41522735 | 6,12109E-08 | 1,63157E-05 | ENSG00000244398 | RPL36AP37 |
| 5815,33448 | 4,297471781 | 0,862358084 | 4,983395947 | 6,24779E-07 | 0,000111023 | ENSG00000112182 | BACH2 |
| 8,791258653 | 3,551002791 | 0,915024507 | 3,880773424 | 0,000104125 | 0,007751201 | ENSG00000148143 | ZNF462 |
| 15,10702187 | 3,327938892 | 0,791330237 | 4,205499468 | 2,60506E-05 | 0,002547412 | ENSG00000227165 | WDR11-DT |
| 16,30520908 | 3,306090298 | 0,862339436 | 3,833861889 | 0,000126147 | 0,008871086 | ENSG00000070182 | SPTB |
| 16,17256435 | 3,251547738 | 0,634923315 | 5,121166072 | 3,03652E-07 | 6,1197E-05 | ENSG00000104722 | NEFM |
| 12,90066212 | 3,22376499 | 0,839724455 | 3,839074795 | 0,000123499 | 0,008787942 | ENSG00000156011 | PSD3 |
| 16,38587118 | 2,834031525 | 0,853681028 | 3,319778034 | 0,00090089 | 0,038670431 | ENSG00000269896 | ENSG00000269896 |
| 8,668090011 | 2,759932412 | 0,80488447 | 3,42897958 | 0,000605855 | 0,029798695 | ENSG00000240591 | ENSG00000240591 |
| 18,1024471 | 2,352688763 | 0,597199971 | 3,939532617 | 8,16405E-05 | 0,006404942 | ENSG00000181577 | LINC03040 |
| 33,13946661 | 2,245308759 | 0,521326831 | 4,306911953 | 1,65549E-05 | 0,001776539 | ENSG00000198626 | RYR2 |
| 23,88802206 | 2,073128382 | 0,492577026 | 4,208739488 | 2,56799E-05 | 0,00252611 | ENSG00000175841 | ENSG00000175841 |
| 20,41718282 | 2,060189809 | 0,583447629 | 3,531062097 | 0,000413895 | 0,021714354 | ENSG00000159871 | LYPD5 |
| 18,19544252 | 1,942145831 | 0,602906114 | 3,221307243 | 0,001276073 | 0,04985432 | ENSG00000136630 | HLX |
| 34,60957483 | 1,798777441 | 0,543149157 | 3,311755929 | 0,000927124 | 0,039590827 | ENSG00000248503 | ENSG00000248503 |
| 37,89304733 | 1,765992435 | 0,44365189 | 3,98058134 | 6,87469E-05 | 0,005624316 | ENSG00000126878 | AIF1L |
| 185,1634291 | 1,661663464 | 0,334117493 | 4,973290833 | 6,58258E-07 | 0,000115727 | ENSG00000162614 | NEXN |
| 19,59493704 | 1,649494032 | 0,509602171 | 3,236826932 | 0,001208667 | 0,048481624 | ENSG00000161921 | CXCL16 |
| 32,86940289 | 1,644952333 | 0,435074026 | 3,780856214 | 0,00015629 | 0,010500654 | ENSG00000141622 | ARK2C |
| 86,59759961 | 1,598312145 | 0,345880817 | 4,620991004 | 3,81911E-06 | 0,000553637 | ENSG00000099958 | DERL3 |
| 41,29847897 | 1,575973298 | 0,377831163 | 4,171104593 | 3,03127E-05 | 0,00283021 | ENSG00000008441 | NFIX |
| 102,9529174 | 1,560996567 | 0,276031517 | 5,655138894 | 1,5572E-08 | 5,04595E-06 | ENSG00000237989 | LINC01679 |
| 25,78725365 | 1,384336913 | 0,421809588 | 3,281900065 | 0,001031101 | 0,043139195 | ENSG00000167034 | NKX3-1 |
| 199,6800934 | 1,370195433 | 0,26348416 | 5,200295287 | 1,98972E-07 | 4,3266E-05 | ENSG00000066923 | STAG3 |
| 140,1555988 | 1,367868334 | 0,214432104 | 6,379027716 | 1,78216E-10 | 9,50063E-08 | ENSG00000160190 | SLC37A1 |
| 105,1001655 | 1,364962265 | 0,313011631 | 4,360739761 | 1,29623E-05 | 0,001509023 | ENSG00000204219 | TCEA3 |
| 33,5045346 | 1,357650996 | 0,401114738 | 3,384694867 | 0,000712574 | 0,03326554 | ENSG00000102996 | MMP15 |
| 50,34883678 | 1,322789648 | 0,352624366 | 3,751271251 | 0,00017594 | 0,011677061 | ENSG00000214595 | EML6 |
| 516,9257723 | 1,287937889 | 0,333012944 | 3,867531019 | 0,000109943 | 0,008004033 | ENSG00000137959 | IFI44L |
| 40,81415639 | 1,272952238 | 0,346927962 | 3,669211994 | 0,000243299 | 0,01505904 | ENSG00000119922 | IFIT2 |
| 518,0824024 | 1,227500625 | 0,22779226 | 5,388684513 | 7,09753E-08 | 1,8474E-05 | ENSG00000145287 | PLAC8 |
| 169,6710014 | 1,214229265 | 0,339546271 | 3,576034754 | 0,000348845 | 0,019485718 | ENSG00000164187 | LMBRD2 |
| 80,15230359 | 1,168863564 | 0,246057468 | 4,75036817 | 2,03047E-06 | 0,000313603 | ENSG00000139832 | RAB20 |
| 47,75580475 | 1,148755849 | 0,320289374 | 3,586618665 | 0,000334994 | 0,019090018 | ENSG00000140479 | PCSK6 |
| 240,015253 | 1,128804411 | 0,237296666 | 4,756933285 | 1,96556E-06 | 0,000309361 | ENSG00000270055 | ENSG00000270055 |
| 129,9173224 | 1,127898513 | 0,332861095 | 3,388496076 | 0,00070277 | 0,032994269 | ENSG00000157483 | MYO1E |
| 4981,928425 | 1,068926972 | 0,203640473 | 5,24908903 | 1,52853E-07 | 3,66094E-05 | ENSG00000128342 | LIF |
| 92,6102906 | 1,061415169 | 0,246606286 | 4,30408805 | 1,67675E-05 | 0,00178774 | ENSG00000081181 | ARG2 |
| 103,0732471 | 1,040274777 | 0,253685265 | 4,100651167 | 4,11989E-05 | 0,003602399 | ENSG00000159128 | IFNGR2 |
| 693,9494153 | 1,039479754 | 0,299288112 | 3,473174212 | 0,000514341 | 0,026153863 | ENSG00000171316 | CHD7 |
| 1125,226742 | 1,032024575 | 0,26372559 | 3,913251556 | 9,10616E-05 | 0,006967054 | ENSG00000112972 | HMGCS1 |
| 59,05535073 | 1,01201789 | 0,30345095 | 3,335029564 | 0,000852903 | 0,037092316 | ENSG00000158457 | TSPAN33 |
| 84,37139726 | -1,001779757 | 0,264963201 | -3,780825988 | 0,000156309 | 0,010500654 | ENSG00000196387 | ZNF140 |
| 112,8151397 | -1,006685789 | 0,300330566 | -3,35192585 | 0,000802515 | 0,035797137 | ENSG00000198964 | SGMS1 |
| 211,9669699 | -1,023498174 | 0,265272815 | -3,858285193 | 0,000114185 | 0,008240294 | ENSG00000138185 | ENTPD1 |
| 1118,040745 | -1,029588059 | 0,145184912 | -7,091563771 | 1,32605E-12 | 9,96104E-10 | ENSG00000132965 | ALOX5AP |
| 325,2048504 | -1,029820941 | 0,191834005 | -5,368291936 | 7,94858E-08 | 2,0209E-05 | ENSG00000105374 | NKG7 |
| 4940,392438 | -1,038485534 | 0,163208267 | -6,362946893 | 1,97919E-10 | 1,02213E-07 | ENSG00000178209 | PLEC |
| 1785,320807 | -1,04786729 | 0,186354844 | -5,622967809 | 1,87704E-08 | 5,96539E-06 | ENSG00000178573 | MAF |
| 71,56146681 | -1,049134797 | 0,276898957 | -3,788872334 | 0,000151333 | 0,010334394 | ENSG00000168672 | LRATD2 |
| 121,4243083 | -1,052132353 | 0,303276589 | -3,469217179 | 0,000521977 | 0,026460727 | ENSG00000238121 | LINC00426 |
| 1352,242598 | -1,055738707 | 0,181928478 | -5,803042607 | 6,51222E-09 | 2,33959E-06 | ENSG00000185989 | RASA3 |
| 467,1279788 | -1,075555706 | 0,284059992 | -3,786368148 | 0,000152865 | 0,010396088 | ENSG00000111145 | ELK3 |
| 407,8068259 | -1,086876307 | 0,190773231 | -5,697216025 | 1,2178E-08 | 4,02506E-06 | ENSG00000092871 | RFFL |
| 146,3092451 | -1,107554542 | 0,340598241 | -3,251791722 | 0,0011468 | 0,046679854 | ENSG00000178163 | ZNF518B |
| 8463,248238 | -1,110340101 | 0,146429545 | -7,58276003 | 3,3828E-14 | 4,30032E-11 | ENSG00000087086 | FTL |
| 252,9308555 | -1,128598518 | 0,217941702 | -5,17844226 | 2,23746E-07 | 4,71559E-05 | ENSG00000121807 | CCR2 |
| 167,3973015 | -1,131237047 | 0,27325176 | -4,139907628 | 3,47446E-05 | 0,003141398 | ENSG00000178996 | SNX18 |
| 681,5577511 | -1,1346807 | 0,245955678 | -4,613354354 | 3,96222E-06 | 0,00056448 | ENSG00000133816 | MICAL2 |
| 128,6043548 | -1,136139761 | 0,266124636 | -4,269201756 | 1,96174E-05 | 0,002051878 | ENSG00000121039 | RDH10 |
| 1071,526465 | -1,138258147 | 0,223562575 | -5,091452131 | 3,55332E-07 | 6,90848E-05 | ENSG00000091592 | NLRP1 |
| 89,61667066 | -1,14450837 | 0,274192793 | -4,174100848 | 2,99165E-05 | 0,002809093 | ENSG00000059377 | TBXAS1 |
| 506,599375 | -1,147474878 | 0,165774443 | -6,921904601 | 4,45611E-12 | 3,06841E-09 | ENSG00000120885 | CLU |
| 149,5084438 | -1,154191222 | 0,316679053 | -3,644671826 | 0,000267733 | 0,016148026 | ENSG00000163867 | ZMYM6 |
| 178,3257652 | -1,158963044 | 0,318607127 | -3,637592968 | 0,000275198 | 0,016477971 | ENSG00000132359 | RAP1GAP2 |
| 312,8498236 | -1,162211067 | 0,314895843 | -3,690779319 | 0,000223568 | 0,014210328 | ENSG00000139278 | GLIPR1 |
| 96,42440607 | -1,176695223 | 0,312247433 | -3,768470447 | 0,000164251 | 0,010945205 | ENSG00000101842 | VSIG1 |

| | | | | | | | |
|---|---|---|---|---|---|---|---|
| 166,851967 | -1,179606879 | 0,214655278 | -5,495354651 | 3,89926E-08 | 1,1507E-05 | ENSG00000016391 | CHDH |
| 392,6864253 | -1,197249221 | 0,278741133 | -4,295201105 | 1,74535E-05 | 0,001848953 | ENSG00000136111 | TBC1D4 |
| 92,81231688 | -1,199885757 | 0,286609751 | -4,186479182 | 2,83315E-05 | 0,002722129 | ENSG00000012779 | ALOX5 |
| 354,3565056 | -1,218846146 | 0,243680109 | -5,001828625 | 5,67891E-07 | 0,000104277 | ENSG00000157978 | LDLRAP1 |
| 831,2996755 | -1,225370632 | 0,178294232 | -6,872744125 | 6,29785E-12 | 4,00301E-09 | ENSG00000064012 | CASP8 |
| 40,72242247 | -1,226681319 | 0,344907701 | -3,556549523 | 0,000375758 | 0,020359906 | ENSG00000179104 | TMTC2 |
| 93,83917832 | -1,22933486 | 0,334852148 | -3,671276613 | 0,000241342 | 0,015050631 | ENSG00000183044 | ABAT |
| 1877,812044 | -1,250768804 | 0,213056902 | -5,870585705 | 4,34258E-09 | 1,7087E-06 | ENSG00000110324 | IL10RA |
| 321,8905651 | -1,256011299 | 0,258915529 | -4,851046612 | 1,22812E-06 | 0,000200949 | ENSG00000114554 | PLXNA1 |
| 254,6136419 | -1,258245449 | 0,202819825 | -6,20375967 | 5,513E-10 | 2,60308E-07 | ENSG00000166963 | MAP1A |
| 36,34619421 | -1,262076481 | 0,368844907 | -3,421699628 | 0,00062231 | 0,030426919 | ENSG00000284634 | ENSG00000284634 |
| 75,53847343 | -1,264611463 | 0,32947382 | -3,838276026 | 0,000123901 | 0,008787942 | ENSG00000267364 | ENSG00000267364 |
| 72,32461973 | -1,268619222 | 0,391192284 | -3,242955636 | 0,001182966 | 0,047682198 | ENSG00000104447 | TRPS1 |
| 50,77130273 | -1,272205827 | 0,351701468 | -3,617288926 | 0,000297705 | 0,017446353 | ENSG00000161551 | ZNF577 |
| 62,39979072 | -1,272438576 | 0,316819576 | -4,016287728 | 5,91221E-05 | 0,004934602 | ENSG00000164849 | GPR146 |
| 53,21830826 | -1,275658907 | 0,322626832 | -3,953976487 | 7,6863E-05 | 0,006166201 | ENSG00000259436 | POU2F2-AS2 |
| 111,8692555 | -1,27780781 | 0,387210544 | -3,300033615 | 0,000966732 | 0,0408599 | ENSG00000282508 | ENSG00000282508 |
| 473,1150843 | -1,278408772 | 0,18555011 | -6,889830324 | 5,5859E-12 | 3,6925E-09 | ENSG00000169194 | IL13 |
| 101,860909 | -1,279761468 | 0,375032532 | -3,412401218 | 0,000643933 | 0,031003711 | ENSG00000262580 | ENSG00000262580 |
| 67,64158294 | -1,283674469 | 0,359142984 | -3,574271327 | 0,000351205 | 0,019542112 | ENSG00000267041 | ZNF850 |
| 1210,062938 | -1,287990926 | 0,243043007 | -5,29943627 | 1,16161E-07 | 2,84946E-05 | ENSG00000165272 | AQP3 |
| 154,3280967 | -1,295840213 | 0,248072614 | -5,223632684 | 1,75447E-07 | 4,12125E-05 | ENSG00000177034 | MTX3 |
| 135,1042019 | -1,296550952 | 0,257205464 | -5,040915273 | 4,63311E-07 | 8,66863E-05 | ENSG00000164120 | HPGD |
| 35,6019444 | -1,297211078 | 0,388369337 | -3,34014804 | 0,000837337 | 0,036900902 | ENSG00000284879 | ENSG00000284879 |
| 68,54190975 | -1,297576762 | 0,28476613 | -4,556640086 | 5,19784E-06 | 0,000704094 | ENSG00000102349 | KLF8 |
| 221,3079071 | -1,306943155 | 0,251023119 | -5,206465284 | 1,92472E-07 | 4,29836E-05 | ENSG00000110665 | C11orf21 |
| 47,33535919 | -1,314527673 | 0,38475247 | -3,416554218 | 0,00063419 | 0,030825381 | ENSG00000167094 | TTC16 |
| 100,2121555 | -1,314757226 | 0,338301592 | -3,886346549 | 0,000101764 | 0,007609752 | ENSG00000118515 | SGK1 |
| 142,9536394 | -1,319912276 | 0,292627508 | -4,510554338 | 6,46584E-06 | 0,000825262 | ENSG00000111537 | IFNG |
| 526,8199341 | -1,320719872 | 0,20353556 | -6,488890052 | 8,6471E-11 | 4,92766E-08 | ENSG00000160791 | CCR5 |
| 236,8407704 | -1,336547515 | 0,265216364 | -5,039460975 | 4,66845E-07 | 8,66863E-05 | ENSG00000113369 | ARRDC3 |
| 71,19542894 | -1,363842614 | 0,396559931 | -3,439184112 | 0,00058347 | 0,028828342 | ENSG00000103253 | HAGHL |
| 13155,0133 | -1,385929536 | 0,207693817 | -6,672945575 | 2,50719E-11 | 1,53459E-08 | ENSG00000173821 | RNF213 |
| 227,2513863 | -1,386654409 | 0,2405833 | -5,763718471 | 8,22806E-09 | 2,89313E-06 | ENSG00000245164 | LINC00861 |
| 78,4298876 | -1,395511793 | 0,302375531 | -4,615161115 | 3,9279E-06 | 0,000564457 | ENSG00000132694 | ARHGEF11 |
| 227,9545778 | -1,400230731 | 0,26845389 | -5,215907778 | 1,82919E-07 | 4,19449E-05 | ENSG00000181523 | SGSH |
| 807,9090356 | -1,403932137 | 0,238375443 | -5,88958376 | 3,8717E-09 | 1,56058E-06 | ENSG00000169756 | LIMS1 |
| 102,1601069 | -1,414575506 | 0,29668625 | -4,767917302 | 1,8614E-06 | 0,000298656 | ENSG00000124212 | PTGIS |
| 109,1188809 | -1,42457599 | 0,234946562 | -6,063404269 | 1,3327E-09 | 5,95249E-07 | ENSG00000275302 | CCL4 |
| 64,61917636 | -1,439038302 | 0,396914153 | -3,625565611 | 0,00028833 | 0,016957068 | ENSG00000177337 | ENSG00000177337 |
| 164,8496964 | -1,440826019 | 0,22513791 | -6,399748576 | 1,55633E-10 | 8,5733E-08 | ENSG00000178860 | MSC |
| 169,9425458 | -1,454418399 | 0,28535541 | -5,096866394 | 3,45322E-07 | 6,87565E-05 | ENSG00000141540 | TTYH2 |
| 97,03919296 | -1,474995717 | 0,296892726 | -4,968109982 | 6,76086E-07 | 0,000117611 | ENSG00000171621 | SPSB1 |
| 82,40956536 | -1,478380094 | 0,269834935 | -5,478831329 | 4,28144E-08 | 1,24132E-05 | ENSG00000164400 | CSF2 |
| 478,6034415 | -1,481350639 | 0,199192781 | -7,436768697 | 1,03178E-13 | 1,0657E-10 | ENSG00000077984 | CST7 |
| 82,74910346 | -1,48391576 | 0,275449601 | -5,387249617 | 7,1544E-08 | 1,8474E-05 | ENSG00000107968 | MAP3K8 |
| 26,65965449 | -1,515778215 | 0,414907563 | -3,653291359 | 0,0002589 | 0,015788134 | ENSG00000178038 | ALS2CL |
| 108,5400923 | -1,523401166 | 0,421821011 | -3,611487159 | 0,000304446 | 0,017778365 | ENSG00000064201 | TSPAN32 |
| 263,5330376 | -1,530155885 | 0,229829654 | -6,657782651 | 2,77989E-11 | 1,64073E-08 | ENSG00000164483 | SAMD3 |
| 156,4450422 | -1,531990196 | 0,350513604 | -4,370701098 | 1,23848E-05 | 0,001472457 | ENSG00000196218 | RYR1 |
| 22,1719237 | -1,538378621 | 0,461197693 | -3,335616467 | 0,000851105 | 0,037092316 | ENSG00000154027 | AK5 |
| 418,7222962 | -1,538691556 | 0,205875104 | -7,473907831 | 7,78477E-14 | 8,57674E-11 | ENSG00000048471 | SNX29 |
| 90,42639405 | -1,547513094 | 0,325503139 | -4,754218654 | 1,99215E-06 | 0,000310588 | ENSG00000073910 | FRY |
| 39,29625821 | -1,563842509 | 0,470152826 | -3,32624292 | 0,000880252 | 0,037882919 | ENSG00000104497 | SNX16 |
| 47,25841295 | -1,565513751 | 0,32083972 | -4,879426247 | 1,06395E-06 | 0,000175828 | ENSG00000230438 | ENSG00000230438 |
| 98,85276317 | -1,576419925 | 0,309453639 | -5,094203868 | 3,5021E-07 | 6,88996E-05 | ENSG00000180549 | FUT7 |
| 282,8700113 | -1,582298196 | 0,215922857 | -7,328071771 | 2,33488E-13 | 2,26977E-10 | ENSG00000171476 | HOPX |
| 50,86261052 | -1,589379251 | 0,383591939 | -4,143411495 | 3,42177E-05 | 0,003141398 | ENSG00000248323 | LUCAT1 |
| 129,3910187 | -1,600866349 | 0,363772636 | -4,400733285 | 1,07886E-05 | 0,0013014 | ENSG00000171246 | NPTX1 |
| 556,6763638 | -1,607735808 | 0,204241178 | -7,871751563 | 3,4971E-15 | 4,81609E-12 | ENSG00000172965 | MIR4435-2HG |
| 39,75714613 | -1,630169152 | 0,456775603 | -3,568862129 | 0,000358535 | 0,019816555 | ENSG00000006704 | GTF2IRD1 |
| 541,3178522 | -1,638199602 | 0,170592146 | -9,60301892 | 7,76365E-22 | 3,20755E-18 | ENSG00000057657 | PRDM1 |
| 19,67134573 | -1,641847326 | 0,507928947 | -3,232435042 | 0,0012274 | 0,04875966 | ENSG00000271361 | ENSG00000271361 |
| 30,73537779 | -1,675834996 | 0,397413391 | -4,216855886 | 2,47732E-05 | 0,002466278 | ENSG00000172348 | RCAN2 |
| 151,9297589 | -1,684769588 | 0,310642769 | -5,423495276 | 5,84448E-08 | 1,58337E-05 | ENSG00000059728 | MXD1 |
| 66,26644782 | -1,727295674 | 0,332500507 | -5,194866286 | 2,04867E-07 | 4,39693E-05 | ENSG00000013619 | MAMLD1 |
| 22,28621106 | -1,732563622 | 0,501202826 | -3,456811358 | 0,000546607 | 0,027624566 | ENSG00000204577 | LILRB3 |
| 126,8362688 | -1,733556905 | 0,240173012 | -7,217950475 | 5,27769E-13 | 4,59048E-10 | ENSG00000277632 | CCL3 |
| 131,2402435 | -1,738793355 | 0,335865664 | -5,177050054 | 2,25422E-07 | 4,71559E-05 | ENSG00000181019 | NQO1 |

| | | | | | | | |
|---|---|---|---|---|---|---|---|
| 33,19544873 | -1,740256252 | 0,496064217 | -3,508126958 | 0,000451274 | 0,023305463 | ENSG00000163106 | HPGDS |
| 345,504096 | -1,781666403 | 0,187118022 | -9,521618403 | 1,70503E-21 | 5,63546E-18 | ENSG00000222041 | CYTOR |
| 23,29182627 | -1,851208283 | 0,477569482 | -3,876311934 | 0,000106052 | 0,007824154 | ENSG00000196422 | PPP1R26 |
| 54,47457758 | -1,890260479 | 0,374932822 | -5,041597772 | 4,61661E-07 | 8,66863E-05 | ENSG00000205336 | ADGRG1 |
| 470,5437404 | -1,907597377 | 0,231777376 | -8,230300182 | 1,86745E-16 | 3,08615E-13 | ENSG00000139679 | LPAR6 |
| 524,1211841 | -1,908840774 | 0,185181986 | -10,30791824 | 6,48917E-25 | 3,57467E-21 | ENSG00000033327 | GAB2 |
| 18,84003797 | -1,91100536 | 0,568279383 | -3,362791994 | 0,000771585 | 0,034935358 | ENSG00000184731 | FAM110C |
| 194,4843739 | -1,911949746 | 0,265777329 | -7,193803006 | 6,30111E-13 | 4,95867E-10 | ENSG00000136161 | RCBTB2 |
| 56,52260023 | -1,913056493 | 0,39125807 | -4,889500407 | 1,01092E-06 | 0,000168753 | ENSG00000115594 | IL1R1 |
| 56,82871009 | -1,918833604 | 0,323420251 | -5,932942036 | 2,97554E-09 | 1,22934E-06 | ENSG00000164530 | PI16 |
| 95,31302809 | -1,926554125 | 0,330505191 | -5,829119108 | 5,57207E-09 | 2,09243E-06 | ENSG00000186594 | MIR22HG |
| 30,90899801 | -1,932081442 | 0,426072828 | -4,534627221 | 5,77053E-06 | 0,00076291 | ENSG00000152217 | SETBP1 |
| 366,4839823 | -1,937341739 | 0,258155931 | -7,504540889 | 6,16443E-14 | 7,27667E-11 | ENSG00000139289 | PHLDA1 |
| 136,4824054 | -1,996704546 | 0,247287257 | -8,074433631 | 6,77906E-16 | 1,01846E-12 | ENSG00000111536 | IL26 |
| 39,56762463 | -2,003474518 | 0,426910318 | -4,692963448 | 2,69276E-06 | 0,000408261 | ENSG00000269001 | ENSG00000269001 |
| 133,9789227 | -2,005904873 | 0,278554625 | -7,201118521 | 5,97206E-13 | 4,93471E-10 | ENSG00000188322 | SBK1 |
| 28,85750427 | -2,033423202 | 0,630018212 | -3,227562576 | 0,001248497 | 0,049125396 | ENSG00000229212 | ENSG00000229212 |
| 18,91028224 | -2,083111146 | 0,581037507 | -3,585157792 | 0,000336874 | 0,019131228 | ENSG00000112115 | IL17A |
| 34,01744177 | -2,086234669 | 0,623986439 | -3,343397451 | 0,000827593 | 0,036667018 | ENSG00000106070 | GRB10 |
| 150,1263163 | -2,092494046 | 0,382907898 | -5,46474506 | 4,63573E-08 | 1,32086E-05 | ENSG00000181045 | SLC26A11 |
| 109,344027 | -2,122209469 | 0,338859104 | -6,262807894 | 3,78106E-10 | 1,83782E-07 | ENSG00000100600 | LGMN |
| 23,44646447 | -2,131814257 | 0,500196528 | -4,261953324 | 2,02648E-05 | 0,002080098 | ENSG00000224383 | PRR29 |
| 84,63912429 | -2,156434126 | 0,297386346 | -7,251288284 | 4,12826E-13 | 3,7902E-10 | ENSG00000134531 | EMP1 |
| 12,06436884 | -2,187492833 | 0,651473708 | -3,357760727 | 0,000785766 | 0,035383021 | ENSG00000163464 | CXCR1 |
| 24,95542206 | -2,199806206 | 0,502522048 | -4,37753172 | 1,20031E-05 | 0,001437413 | ENSG00000008226 | DLEC1 |
| 241,9148783 | -2,209864439 | 0,317403195 | -6,962325753 | 3,347E-12 | 2,4049E-09 | ENSG00000126822 | PLEKHG3 |
| 21,2150224 | -2,215674295 | 0,509592995 | -4,347929263 | 1,37429E-05 | 0,001545 | ENSG00000198848 | CES1 |
| 55,62621736 | -2,245634228 | 0,430732174 | -5,213527948 | 1,85283E-07 | 4,19449E-05 | ENSG00000143776 | CDC42BPA |
| 25,78131236 | -2,292088795 | 0,52628716 | -4,355205612 | 1,32942E-05 | 0,001515173 | ENSG00000279765 | ENSG00000279765 |
| 27,41919022 | -2,324347304 | 0,588873931 | -3,947105115 | 7,91018E-05 | 0,006254722 | ENSG00000062524 | LTK |
| 51,17980613 | -2,32857413 | 0,576277537 | -4,040716459 | 5,32882E-05 | 0,004518853 | ENSG00000123836 | PFKFB2 |
| 93,34250186 | -2,366804338 | 0,287254769 | -8,239390931 | 1,7309E-16 | 3,08615E-13 | ENSG00000127318 | IL22 |
| 22,96744126 | -2,47297472 | 0,547597435 | -4,51604511 | 6,30052E-06 | 0,000819862 | ENSG00000168811 | IL12A |
| 18,80214147 | -2,49963631 | 0,576187504 | -4,338234152 | 1,43632E-05 | 0,001603827 | ENSG00000243836 | WDR86-AS1 |
| 209,1673577 | -2,524715787 | 0,302485241 | -8,346575133 | 7,02709E-17 | 1,45162E-13 | ENSG00000157570 | TSPAN18 |
| 14,37087873 | -2,598633142 | 0,643374213 | -4,03906947 | 5,36637E-05 | 0,004518853 | ENSG00000109956 | B3GAT1 |
| 13,09590847 | -2,720175126 | 0,749651298 | -3,628587229 | 0,000284976 | 0,016819718 | ENSG00000073756 | PTGS2 |
| 9,307887484 | -2,863450413 | 0,885387324 | -3,234121763 | 0,001220174 | 0,04875966 | ENSG00000251301 | ENSG00000251301 |
| 40,21721324 | -2,874339281 | 0,479936901 | -5,988994126 | 2,11143E-09 | 8,94704E-07 | ENSG00000138678 | GPAT3 |
| 236,8783193 | -2,880267508 | 0,253519137 | -11,36114434 | 6,52838E-30 | 1,07888E-25 | ENSG00000056736 | IL17RB |
| 16,31175126 | -2,896375742 | 0,640777633 | -4,520094951 | 6,18119E-06 | 0,000810717 | ENSG00000066735 | KIF26A |
| 2632,364376 | -2,923473778 | 0,267388434 | -10,93343393 | 7,97714E-28 | 6,59151E-24 | ENSG00000225783 | MIAT |
| 12,02255615 | -2,926526242 | 0,761875965 | -3,841210874 | 0,000122429 | 0,0087587 | ENSG00000100292 | HMOX1 |
| 199,6756305 | -3,067641399 | 0,356220195 | -8,611643705 | 7,20204E-18 | 1,7003E-14 | ENSG00000169896 | ITGAM |
| 9,903145885 | -3,203935385 | 0,944650665 | -3,391661598 | 0,000694702 | 0,032801836 | ENSG00000277693 | ENSG00000277693 |
| 14,24182581 | -3,508183428 | 0,739587192 | -4,743434532 | 2,10125E-06 | 0,00032153 | ENSG00000169429 | CXCL8 |
| 60,71657136 | -4,208421562 | 0,466267605 | -9,025764424 | 1,78445E-19 | 4,91497E-16 | ENSG00000156510 | HKDC1 |

LV_BACH2_vs_GFP_d25

| baseMean | log2FoldChange | lfcSE | stat | pvalue | padj | ENSEMBL | SYMBOL |
|---|---|---|---|---|---|---|---|
| 42,10690893 | 1,766621692 | 0,558178465 | 3,164976442 | 0,001550955 | 0,049289469 | ENSG00000119514 | GALNT12 |
| 56,31125049 | -1,537697839 | 0,485334562 | -3,168325441 | 0,001533198 | 0,048909013 | ENSG00000156510 | HKDC1 |
| 85,19183722 | -1,322279564 | 0,416944436 | -3,171356778 | 0,001517287 | 0,048676975 | ENSG00000273338 | ENSG00000273338 |
| 49,01901219 | -1,72559844 | 0,543536454 | -3,174761194 | 0,001499599 | 0,048292788 | ENSG00000153814 | JAZF1 |
| 135,9908463 | -1,385774497 | 0,435871625 | -3,179317985 | 0,00147622 | 0,047904913 | ENSG00000198756 | COLGALT2 |
| 105,1859207 | -1,199274457 | 0,37658464 | -3,184608 | 0,001449502 | 0,04752033 | ENSG00000274591 | ENSG00000274591 |
| 48,15837758 | 1,961979942 | 0,615724335 | 3,18645834 | 0,001440262 | 0,047374542 | ENSG00000112812 | PRSS16 |
| 153,5863128 | 1,2514568 | 0,392417196 | 3,189097754 | 0,001427176 | 0,047035606 | ENSG00000236552 | RPL13AP5 |
| 18,96884816 | 2,650480305 | 0,830457472 | 3,191590653 | 0,001414917 | 0,04672266 | ENSG00000164283 | ESM1 |
| 44,51445629 | 1,88025097 | 0,583641761 | 3,22158402 | 0,001274841 | 0,043021423 | ENSG00000164056 | SPRY1 |
| 106,6350924 | -1,303632926 | 0,404215166 | -3,225096519 | 0,001259302 | 0,042703183 | ENSG00000155629 | PIK3AP1 |
| 28,2338041 | -2,276561567 | 0,705430143 | -3,227196328 | 0,001250097 | 0,04252592 | ENSG00000163421 | PROK2 |
| 148,3843419 | 1,12700835 | 0,34907949 | 3,228514942 | 0,001244348 | 0,042501383 | ENSG00000114107 | CEP70 |
| 233,0969496 | 1,044099172 | 0,323175378 | 3,230750993 | 0,001234654 | 0,042341387 | ENSG00000107438 | PDLIM1 |
| 99,36782456 | 1,536978177 | 0,475683687 | 3,231092886 | 0,001233179 | 0,042341387 | ENSG00000138449 | SLC40A1 |
| 135,0584419 | -1,685246711 | 0,519014704 | -3,247011493 | 0,001166237 | 0,04032222 | ENSG00000184357 | H1-5 |
| 138,13001 | -1,269850121 | 0,390338438 | -3,253202853 | 0,00114112 | 0,039651572 | ENSG00000180611 | MB21D2 |
| 253,6138419 | -1,160807027 | 0,356847846 | -3,252946711 | 0,001142149 | 0,039651572 | ENSG00000121039 | RDH10 |
| 24,20670755 | 2,526398225 | 0,775936087 | 3,255935981 | 0,001130192 | 0,039398271 | ENSG00000166689 | PLEKHA7 |
| 187,2436526 | 1,285785378 | 0,392304518 | 3,277518662 | 0,001047238 | 0,037118772 | ENSG00000211785 | TRAV12-1 |
| 120,7774068 | -1,316016531 | 0,398560947 | -3,301920426 | 0,000960253 | 0,034764451 | ENSG00000177468 | OLIG3 |
| 278,513808 | 1,099061212 | 0,332597787 | 3,304475417 | 0,000951543 | 0,034597292 | ENSG00000186399 | GOLGA8R |
| 40,82223165 | -2,920814642 | 0,883007481 | -3,307802826 | 0,00094031 | 0,034310929 | ENSG00000205809 | KLRC2 |
| 46,98360716 | 1,833277842 | 0,554294507 | 3,30740756 | 0,000941638 | 0,034310929 | ENSG00000178115 | GOLGA8Q |
| 228,8652534 | 1,032585392 | 0,311872125 | 3,310925569 | 0,000929879 | 0,03427446 | ENSG00000147124 | ZNF41 |
| 36,71482003 | 2,264531082 | 0,683133454 | 3,314917559 | 0,000916702 | 0,033988319 | ENSG00000255036 | SUGT1P4-STRA6LP-CCDC180 |
| 27,21820414 | 3,3288017 | 1,002996628 | 3,318856323 | 0,000903869 | 0,033809108 | ENSG00000173947 | CIMAP3 |
| 102,6458007 | 1,542067574 | 0,463768704 | 3,32507899 | 0,000883935 | 0,033210416 | ENSG00000018408 | WWTR1 |
| 280,370548 | -1,271975662 | 0,380027571 | -3,347061526 | 0,000816731 | 0,031194121 | ENSG00000182240 | BACE2 |
| 18,69514074 | -2,957911504 | 0,880903692 | -3,357814858 | 0,000785612 | 0,03036683 | ENSG00000189143 | CLDN4 |
| 25,72299389 | 2,523934215 | 0,751722887 | 3,357532752 | 0,000786414 | 0,03036683 | ENSG00000155966 | AFF2 |
| 127,0609481 | -1,756823734 | 0,520701375 | -3,373956395 | 0,000740961 | 0,028931705 | ENSG00000000938 | FGR |
| 105,5938684 | 1,353477788 | 0,398710381 | 3,394638945 | 0,000687192 | 0,027401765 | ENSG00000113319 | RASGRF2 |
| 47,26882122 | -2,452128128 | 0,720526656 | -3,40324415 | 0,000665907 | 0,026678904 | ENSG00000222328 | ENSG00000222328 |
| 210,1358943 | -1,375642732 | 0,404055188 | -3,404591186 | 0,000662632 | 0,026610724 | ENSG00000111537 | IFNG |
| 722,5443779 | -2,688381509 | 0,78392711 | -3,429376882 | 0,000604969 | 0,025169124 | ENSG00000277632 | CCL3 |
| 176,5574694 | 1,114798935 | 0,323735705 | 3,443546444 | 0,000574138 | 0,024328197 | ENSG00000221926 | TRIM16 |
| 250,9625581 | -1,210857671 | 0,351478648 | -3,445039058 | 0,000570977 | 0,02425504 | ENSG00000206652 | RNU1-1 |
| 359,9081929 | 1,082423597 | 0,313750732 | 3,449947639 | 0,000560695 | 0,023878275 | ENSG00000133985 | TTC9 |
| 283,7207669 | -1,118724309 | 0,323474023 | -3,458467233 | 0,000543258 | 0,02331185 | ENSG00000124212 | PTGIS |
| 229,5934686 | 1,073516058 | 0,310209089 | 3,46062091 | 0,000538931 | 0,023209757 | ENSG00000008300 | CELSR3 |
| 32,73318603 | 2,904953127 | 0,839483095 | 3,46040694 | 0,00053936 | 0,023209757 | ENSG00000187416 | LHFPL3 |
| 762,9261156 | 1,148320185 | 0,331197298 | 3,467178603 | 0,000525952 | 0,02285932 | ENSG00000117650 | NEK2 |
| 188,4909356 | 1,095686032 | 0,315939236 | 3,468027734 | 0,000524293 | 0,02284594 | ENSG00000166780 | BMERB1 |
| 196,5575037 | 1,110675301 | 0,320009196 | 3,470760579 | 0,000518987 | 0,022731878 | ENSG00000256988 | ENSG00000256988 |
| 277,5449294 | 1,07449833 | 0,308817992 | 3,479390311 | 0,000502556 | 0,022285565 | ENSG00000224429 | LINC00539 |
| 153,0538037 | 1,46348974 | 0,419701775 | 3,486975344 | 0,000488516 | 0,021850123 | ENSG00000053918 | KCNQ1 |
| 200,4639095 | -1,070013506 | 0,306637218 | -3,489509573 | 0,000483908 | 0,0217014 | ENSG00000224259 | LINC01133 |
| 55,9706593 | -1,714482892 | 0,490638637 | -3,494390298 | 0,000475146 | 0,021664815 | ENSG00000267452 | LINC02072 |
| 180,2508516 | 1,378351989 | 0,394433942 | 3,494506539 | 0,000474939 | 0,021664815 | ENSG00000119408 | NEK6 |
| 38,59375677 | 2,311463656 | 0,661095104 | 3,496416236 | 0,000471553 | 0,021664815 | ENSG00000182871 | COL18A1 |
| 52,02167432 | 1,968907988 | 0,561460537 | 3,506761133 | 0,000453596 | 0,021068543 | ENSG00000188385 | JAKMIP3 |
| 102,7019853 | 1,402091294 | 0,399291014 | 3,511452161 | 0,000445666 | 0,020757216 | ENSG00000174500 | GCSAM |
| 268,6009611 | -1,030101651 | 0,293229614 | -3,512952314 | 0,000443157 | 0,020697391 | ENSG00000269028 | ENSG00000269028 |
| 104,0079938 | -2,134783197 | 0,606944434 | -3,517262994 | 0,000436022 | 0,020446801 | ENSG00000114554 | PLXNA1 |
| 214,7391681 | -1,556835174 | 0,441999056 | -3,52225905 | 0,000427886 | 0,020207445 | ENSG00000228049 | POLR2J2 |
| 18,06724646 | 4,604615129 | 1,306410217 | 3,524631903 | 0,000424072 | 0,020089204 | ENSG00000143341 | HMCN1 |
| 20,52376033 | 3,205435313 | 0,90777361 | 3,531095505 | 0,000413842 | 0,019709385 | ENSG00000137462 | TLR2 |
| 65,04318631 | -1,807515376 | 0,511196065 | -3,535855418 | 0,000406457 | 0,019437461 | ENSG00000152217 | SETBP1 |
| 207,5047055 | -1,113019989 | 0,31329893 | -3,552581526 | 0,000381471 | 0,018533123 | ENSG00000177337 | ENSG00000177337 |
| 130,5451973 | 1,280520828 | 0,360122407 | 3,555793263 | 0,00037684 | 0,018360921 | ENSG00000172828 | CES3 |
| 61,25320681 | 1,700419095 | 0,47795289 | 3,55771276 | 0,000374098 | 0,018279988 | ENSG00000038382 | TRIO |
| 207,3151306 | 1,392468502 | 0,390391896 | 3,566847865 | 0,000361301 | 0,017809089 | ENSG00000135709 | KIAA0513 |
| 28,84608058 | -2,470123146 | 0,691152052 | -3,573921451 | 0,000351674 | 0,017436247 | ENSG00000169583 | CLIC3 |
| 69,82018374 | -1,701529876 | 0,475153807 | -3,581008615 | 0,00034227 | 0,017070104 | ENSG00000187240 | DYNC2H1 |
| 55,48495921 | 1,900306898 | 0,530114932 | 3,584707357 | 0,000337456 | 0,016929896 | ENSG00000196205 | ENSG00000196205 |
| 286,6495202 | 1,10049702 | 0,306896241 | 3,585892796 | 0,000335927 | 0,016903324 | ENSG00000240350 | ENSG00000240350 |
| 203,6826983 | -1,181295266 | 0,329295767 | -3,587338147 | 0,000334071 | 0,016889929 | ENSG00000049759 | NEDD4L |
| 64,3464902 | 1,812186538 | 0,505226804 | 3,586877266 | 0,000334662 | 0,016889929 | ENSG00000109265 | CRACD |
| 99,65156127 | -1,536670715 | 0,425945382 | -3,607670798 | 0,000308958 | 0,015781134 | ENSG00000115607 | IL18RAP |
| 117,7600727 | -1,335176923 | 0,369193962 | -3,616464681 | 0,000298654 | 0,015455419 | ENSG00000284634 | ENSG00000284634 |
| 52,00955656 | 2,078113912 | 0,574126512 | 3,619609738 | 0,000295048 | 0,015396206 | ENSG00000186265 | BTLA |
| 40,44094042 | -2,057091689 | 0,567944159 | -3,62199638 | 0,000292338 | 0,01530205 | ENSG00000116711 | PLA2G4A |

| | | | | | | | |
|---|---|---|---|---|---|---|---|
| 283,6113674 | 1,21207605 | 0,33461982 | 3,622248228 | 0,000292054 | 0,01530205 | ENSG00000064886 | CHI3L2 |
| 390,6153107 | 1,179529902 | 0,325316843 | 3,625787988 | 0,000288082 | 0,015173197 | ENSG00000118513 | MYB |
| 22,65911769 | 2,831499725 | 0,778637737 | 3,63647893 | 0,00027639 | 0,014694748 | ENSG00000103888 | CEMIP |
| 333,4196998 | 1,1557746 | 0,31468936 | 3,672747625 | 0,000239956 | 0,012961481 | ENSG00000285565 | ENSG00000285565 |
| 133,2077034 | -1,439197608 | 0,391185006 | -3,679071499 | 0,000234085 | 0,012802607 | ENSG00000196083 | IL1RAP |
| 282,0353553 | 1,054890185 | 0,286894104 | 3,676932254 | 0,000236056 | 0,012802607 | ENSG00000106991 | ENG |
| 273,7443378 | 1,13334615 | 0,308226975 | 3,6769856 | 0,000236006 | 0,012802607 | ENSG00000115155 | OTOF |
| 35,62475084 | 2,403025903 | 0,653579829 | 3,676713688 | 0,000236258 | 0,012802607 | ENSG00000273540 | AGBL1 |
| 79,97504573 | -1,645588322 | 0,447011781 | -3,681308622 | 0,00023204 | 0,012778824 | ENSG00000217825 | ENSG00000217825 |
| 27,88889648 | -3,355519451 | 0,911097046 | -3,682944059 | 0,000230556 | 0,012760497 | ENSG00000100427 | MLC1 |
| 256,5805358 | -1,132764207 | 0,307011691 | -3,689645183 | 0,000224567 | 0,012530544 | ENSG00000143842 | SOX13 |
| 51,68650294 | -1,919521611 | 0,519925777 | -3,691914686 | 0,000222572 | 0,012460354 | ENSG00000232022 | ENSG00000232022 |
| 55,01157277 | -2,099217914 | 0,568319073 | -3,693731237 | 0,000220987 | 0,012412736 | ENSG00000283554 | LINC02341 |
| 30,47395085 | -2,883025412 | 0,778212712 | -3,704675299 | 0,000211662 | 0,011928547 | ENSG00000127318 | IL22 |
| 116,0985172 | -1,448396603 | 0,389220092 | -3,721279127 | 0,000198216 | 0,011245776 | ENSG00000238741 | SCARNA7 |
| 33,11165484 | 3,021040983 | 0,808979011 | 3,734387345 | 0,000188173 | 0,010784528 | ENSG00000260784 | ENSG00000260784 |
| 20,26197923 | 3,609531598 | 0,966242504 | 3,73563736 | 0,00018724 | 0,010767593 | ENSG00000171914 | TLN2 |
| 4658,301778 | 1,002994148 | 0,267749985 | 3,746010104 | 0,000179669 | 0,010510968 | ENSG00000117724 | CENPF |
| 78,05079125 | -1,62401871 | 0,432475625 | -3,755168188 | 0,000173225 | 0,010312384 | ENSG00000266964 | FXYD1 |
| 44,80861812 | -2,222013627 | 0,590747428 | -3,7613598 | 0,000168992 | 0,010131738 | ENSG00000196139 | AKR1C3 |
| 988,039341 | 1,030968001 | 0,273409804 | 3,770779204 | 0,000162739 | 0,009859663 | ENSG00000156970 | BUB1B |
| 1214,894517 | 1,095464458 | 0,290578862 | 3,769938567 | 0,000163288 | 0,009859663 | ENSG00000131242 | RAB11FIP4 |
| 203,8928337 | -1,538270542 | 0,407396072 | -3,77586002 | 0,000159457 | 0,009732605 | ENSG00000230562 | ENSG00000230562 |
| 532,3020725 | 1,208546906 | 0,319529511 | 3,782270073 | 0,000155405 | 0,009554276 | ENSG00000183963 | SMTN |
| 25,82751505 | -3,047233377 | 0,800002768 | -3,80902854 | 0,000139514 | 0,008700563 | ENSG00000196422 | PPP1R26 |
| 508,7419695 | 1,002434066 | 0,262880087 | 3,813275012 | 0,000137137 | 0,008619262 | ENSG00000112679 | DUSP22 |
| 44,67969643 | -2,897065388 | 0,757653173 | -3,823735569 | 0,000131445 | 0,008395757 | ENSG00000004399 | PLXND1 |
| 626,6851966 | 1,192229923 | 0,311895827 | 3,822526041 | 0,000132092 | 0,008395757 | ENSG00000126353 | CCR7 |
| 74,08113889 | 1,903923558 | 0,497091724 | 3,830125238 | 0,000128078 | 0,008264945 | ENSG00000280135 | ENSG00000280135 |
| 115,4073723 | -1,490207184 | 0,388731808 | -3,833509772 | 0,000126328 | 0,008183233 | ENSG00000081181 | ARG2 |
| 230,5577132 | 1,239994563 | 0,323024233 | 3,838704459 | 0,000123685 | 0,008042868 | ENSG00000136295 | TTYH3 |
| 64,67593711 | -1,859420022 | 0,484178651 | -3,840359375 | 0,000122854 | 0,008019684 | ENSG00000180263 | FGD6 |
| 2770,96015 | 1,018260567 | 0,263320542 | 3,86700011 | 0,000110182 | 0,007334069 | ENSG00000271383 | NBPF19 |
| 339,9661139 | 1,023421898 | 0,264545806 | 3,868599971 | 0,000109462 | 0,007314921 | ENSG00000280893 | ENSG00000280893 |
| 131,0915536 | 1,364896983 | 0,352737473 | 3,86944141 | 0,000109085 | 0,007314921 | ENSG00000083454 | P2RX5 |
| 3035,192427 | -1,13078044 | 0,290741286 | -3,889301223 | 0,000100533 | 0,006826169 | ENSG00000090104 | RGS1 |
| 657,7207801 | 1,051221653 | 0,270173266 | 3,890916633 | 9,98662E-05 | 0,00680822 | ENSG00000120896 | SORBS3 |
| 272,0220799 | -1,467912826 | 0,376626825 | -3,897525951 | 9,71804E-05 | 0,00665194 | ENSG00000049249 | TNFRSF9 |
| 988,7516523 | 1,144382744 | 0,292988705 | 3,905893729 | 9,38778E-05 | 0,006451997 | ENSG00000171316 | CHD7 |
| 257,3041018 | -1,231891548 | 0,313523066 | -3,929189524 | 8,52326E-05 | 0,005930158 | ENSG00000115604 | IL18R1 |
| 420,0617848 | 1,029093878 | 0,261603673 | 3,93378986 | 8,36169E-05 | 0,00586602 | ENSG00000130766 | SESN2 |
| 565,640631 | 1,047346337 | 0,264983295 | 3,95249949 | 7,73391E-05 | 0,005517182 | ENSG00000234664 | ENSG00000234664 |
| 60,72703079 | 1,925832985 | 0,486563799 | 3,958027684 | 7,55712E-05 | 0,005413908 | ENSG00000280987 | MATR3 |
| 185,4244407 | 1,308587022 | 0,33043886 | 3,960148698 | 7,49031E-05 | 0,005400945 | ENSG00000121742 | GJB6 |
| 124,737879 | 1,4562855 | 0,366441245 | 3,974130968 | 7,06367E-05 | 0,005147649 | ENSG00000274422 | ENSG00000274422 |
| 1521,387559 | -1,265031705 | 0,317362095 | -3,986083172 | 6,7173E-05 | 0,00495936 | ENSG00000184557 | SOCS3 |
| 325,792391 | -1,43573014 | 0,359911618 | -3,989118633 | 6,63192E-05 | 0,004939469 | ENSG00000211829 | ENSG00000211829 |
| 195,590455 | 1,550041226 | 0,388293753 | 3,991929341 | 6,55379E-05 | 0,004902873 | ENSG00000269897 | COMMD3-BMI1 |
| 83,78910456 | 2,756980405 | 0,682656033 | 4,038608421 | 5,37692E-05 | 0,004113468 | ENSG00000101134 | DOK5 |
| 120,7717549 | -1,614754296 | 0,398260318 | -4,054519675 | 5,02374E-05 | 0,003878375 | ENSG00000176485 | PLAAT3 |
| 82,76658048 | -1,793518479 | 0,440909253 | -4,067772376 | 4,74647E-05 | 0,003681128 | ENSG00000115657 | ABCB6 |
| 363,9963122 | 1,232739528 | 0,302983354 | 4,068670806 | 4,72821E-05 | 0,003681128 | ENSG00000066923 | STAG3 |
| 98,45348849 | 2,388966908 | 0,586928618 | 4,07028527 | 4,69556E-05 | 0,003675363 | ENSG00000184602 | SNN |
| 491,2312525 | 1,087783498 | 0,266656629 | 4,079341668 | 4,51634E-05 | 0,003551525 | ENSG00000072110 | ACTN1 |
| 602,6446005 | 1,120995143 | 0,274675051 | 4,081168414 | 4,48099E-05 | 0,003551525 | ENSG00000127947 | PTPN12 |
| 319,2915606 | 1,262798929 | 0,309469 | 4,080534489 | 4,49323E-05 | 0,003551525 | ENSG00000163814 | CDCP1 |
| 132,5049956 | 2,071979312 | 0,50787326 | 4,079717278 | 4,50905E-05 | 0,003551525 | ENSG00000158457 | TSPAN33 |
| 481,1687755 | 1,01523449 | 0,248344884 | 4,088002436 | 4,35104E-05 | 0,003486396 | ENSG00000137824 | RMDN3 |
| 21,27702026 | -3,703933638 | 0,904814518 | -4,093583339 | 4,24757E-05 | 0,003426117 | ENSG00000226928 | ENSG00000226928 |
| 291,2609398 | 1,165830908 | 0,284824945 | 4,093148888 | 4,25554E-05 | 0,003426117 | ENSG00000198520 | ARMH1 |
| 637,0336167 | 1,089159529 | 0,265494565 | 4,102379758 | 4,08922E-05 | 0,003339928 | ENSG00000189060 | H1-0 |
| 403,0072737 | -1,056318547 | 0,257219264 | -4,106685219 | 4,01378E-05 | 0,003310289 | ENSG00000124588 | NQO2 |
| 44,39179784 | 3,05166663 | 0,742862739 | 4,107981825 | 3,99132E-05 | 0,0033079 | ENSG00000258830 | ENSG00000258830 |
| 669,7475554 | 1,057169598 | 0,255537486 | 4,137043121 | 3,5181E-05 | 0,002974026 | ENSG00000152223 | EPG5 |
| 229,9044687 | 1,630287638 | 0,393443909 | 4,143634204 | 3,41845E-05 | 0,002918975 | ENSG00000113749 | HRH2 |
| 51,3068531 | -2,350551249 | 0,566419112 | -4,149844522 | 3,32701E-05 | 0,002869887 | ENSG00000287061 | ENSG00000287061 |
| 746,0597344 | 1,128462043 | 0,271885115 | 4,150510564 | 3,31734E-05 | 0,002869887 | ENSG00000103381 | CPPED1 |
| 69,94041494 | 2,479945234 | 0,597394842 | 4,151266568 | 3,3064E-05 | 0,002869887 | ENSG00000144785 | ENSG00000144785 |
| 117,4201206 | 1,826699956 | 0,439708502 | 4,154343044 | 3,26223E-05 | 0,002857751 | ENSG00000150054 | MPP7 |
| 1854,652946 | 1,015444523 | 0,243659291 | 4,167477122 | 3,07989E-05 | 0,002712071 | ENSG00000172794 | RAB37 |
| 5317,401888 | 1,016742341 | 0,243394678 | 4,177340073 | 2,94938E-05 | 0,002610741 | ENSG00000143401 | ANP32E |
| 245,9883686 | -1,468285738 | 0,351320091 | -4,179338939 | 2,92358E-05 | 0,002601522 | ENSG00000164120 | HPGD |
| 358,454178 | 1,205368874 | 0,28794945 | 4,186043324 | 2,83859E-05 | 0,00255277 | ENSG00000104081 | BMF |
| 37,85729728 | -3,380638241 | 0,80571775 | -4,195809561 | 2,71899E-05 | 0,002458284 | ENSG00000197057 | DTHD1 |
| 480,523254 | -1,150323976 | 0,272892525 | -4,215300419 | 2,49446E-05 | 0,002292055 | ENSG00000178996 | SNX18 |

| | | | | | | | |
|---|---|---|---|---|---|---|---|
| 33,09334825 | 3,792528182 | 0,898393928 | 4,22145349 | 2,42732E-05 | 0,002254874 | ENSG00000164932 | CTHRC1 |
| 177,8637539 | 2,093518247 | 0,494569173 | 4,233014029 | 2,3058E-05 | 0,002177887 | ENSG00000197408 | CYP2B6 |
| 76,40471647 | 2,237826447 | 0,527898644 | 4,239121417 | 2,24396E-05 | 0,002143428 | ENSG00000089199 | CHGB |
| 698,0190864 | 1,065227803 | 0,249497779 | 4,269488117 | 1,95922E-05 | 0,001882077 | ENSG00000167220 | HDHD2 |
| 183,9172734 | -1,377463526 | 0,322491857 | -4,271312588 | 1,94326E-05 | 0,001877409 | ENSG00000171621 | SPSB1 |
| 1087,179319 | 1,06049583 | 0,248260278 | 4,271709678 | 1,9398E-05 | 0,001877409 | ENSG00000006715 | VPS41 |
| 351,9129926 | -1,19944211 | 0,279784104 | -4,287027372 | 1,8108E-05 | 0,001790362 | ENSG00000174792 | ODAPH |
| 1107,752808 | 1,245862751 | 0,290529056 | 4,288255259 | 1,80082E-05 | 0,001790362 | ENSG00000166326 | TRIM44 |
| 302,3195984 | 1,597676679 | 0,372641501 | 4,287436245 | 1,80747E-05 | 0,001790362 | ENSG00000134256 | CD101 |
| 115,5938352 | -1,840162727 | 0,428260288 | -4,29683251 | 1,73256E-05 | 0,001746516 | ENSG00000198848 | CES1 |
| 1719,026666 | -1,060357611 | 0,246414687 | -4,3031429 | 1,68392E-05 | 0,001715064 | ENSG00000010404 | IDS |
| 238,5440168 | 1,643457228 | 0,38179512 | 4,304552731 | 1,67323E-05 | 0,001714507 | ENSG00000137331 | IER3 |
| 732,511088 | -1,21669874 | 0,28198203 | -4,314809486 | 1,59741E-05 | 0,001646789 | ENSG00000069702 | TGFBR3 |
| 568,7216454 | 1,144573287 | 0,264718588 | 4,323735989 | 1,53409E-05 | 0,001591217 | ENSG00000159496 | RGL4 |
| 1892,609947 | -1,400146142 | 0,323562394 | -4,327283297 | 1,5096E-05 | 0,001579095 | ENSG00000172543 | CTSW |
| 253,4706154 | 1,381197641 | 0,319220814 | 4,326778146 | 1,51306E-05 | 0,001579095 | ENSG00000204104 | TRAF3IP1 |
| 812,6800109 | 1,103909796 | 0,25447581 | 4,337975363 | 1,43801E-05 | 0,00151953 | ENSG00000130787 | HIP1R |
| 305,9262257 | 1,195540237 | 0,275462219 | 4,340124178 | 1,42402E-05 | 0,00151421 | ENSG00000148384 | INPP5E |
| 22,69743377 | 4,698898338 | 1,078533798 | 4,35674649 | 1,3201E-05 | 0,001412591 | ENSG00000263647 | BPTFP1 |
| 247,1029679 | -1,450844483 | 0,330491204 | -4,389963985 | 1,13369E-05 | 0,001220852 | ENSG00000205002 | AARD |
| 54,8866102 | -2,415492414 | 0,54989514 | -4,392641868 | 1,11982E-05 | 0,001213636 | ENSG00000100292 | HMOX1 |
| 115,8943692 | 1,768361179 | 0,402180669 | 4,396932307 | 1,09792E-05 | 0,001197578 | ENSG00000124191 | TOX2 |
| 1869,173821 | -1,002701962 | 0,227597013 | -4,405602462 | 1,0549E-05 | 0,001158132 | ENSG00000143013 | LMO4 |
| 1425,866408 | 1,119548852 | 0,254020788 | 4,407311934 | 1,04661E-05 | 0,001156543 | ENSG00000103365 | GGA2 |
| 5181,431396 | 1,124721033 | 0,255161974 | 4,407870873 | 1,04392E-05 | 0,001156543 | ENSG00000150593 | PDCD4 |
| 67,52978299 | -2,046909119 | 0,462522547 | -4,425533705 | 9,62041E-06 | 0,001077168 | ENSG00000079156 | OSBPL6 |
| 852,9524927 | 1,01548811 | 0,228616188 | 4,441890656 | 8,91719E-06 | 0,001010444 | ENSG00000181827 | RFX7 |
| 242,0384771 | 1,32944248 | 0,299373637 | 4,440746672 | 8,96473E-06 | 0,001010444 | ENSG00000205476 | CCDC85C |
| 882,3472869 | 1,015134055 | 0,226167534 | 4,488416344 | 7,17546E-06 | 0,000825275 | ENSG00000164402 | SEPTIN8 |
| 25,14360921 | -4,78401979 | 1,064126702 | -4,495723845 | 6,93337E-06 | 0,000802893 | ENSG00000253230 | MIR124-1HG |
| 1909,421588 | -1,324951884 | 0,293667371 | -4,511743606 | 6,42969E-06 | 0,000749702 | ENSG00000136810 | TXN |
| 266,9585911 | 1,333072209 | 0,295457585 | 4,511890284 | 6,42524E-06 | 0,000749702 | ENSG00000141753 | IGFBP4 |
| 93,18962711 | 2,112045485 | 0,467681929 | 4,515986942 | 6,30225E-06 | 0,00074512 | ENSG00000164512 | ANKRD55 |
| 246,3870503 | 1,631425517 | 0,361093439 | 4,518014846 | 6,24221E-06 | 0,000743219 | ENSG00000154358 | OBSCN |
| 47,06021121 | -2,448548843 | 0,53887412 | -4,54382341 | 5,52429E-06 | 0,000662406 | ENSG00000168994 | PXDC1 |
| 6891,684546 | 1,118660564 | 0,245851955 | 4,550138974 | 5,36105E-06 | 0,000647423 | ENSG00000167106 | EEIG1 |
| 367,3744686 | 1,265674101 | 0,277951874 | 4,553572825 | 5,27424E-06 | 0,000641522 | ENSG00000136040 | PLXNC1 |
| 718,1288156 | -1,156571933 | 0,253029465 | -4,570898236 | 4,85638E-06 | 0,000596849 | ENSG00000160791 | CCR5 |
| 804,0805829 | 1,055132592 | 0,230870274 | 4,57024014 | 4,87166E-06 | 0,000596849 | ENSG00000119139 | TJP2 |
| 150,7201942 | 1,879968697 | 0,40964427 | 4,589271318 | 4,44796E-06 | 0,000552953 | ENSG00000171777 | RASGRP4 |
| 1352,459974 | 1,169387156 | 0,253815322 | 4,607236256 | 4,08056E-06 | 0,000511038 | ENSG00000136108 | CKAP2 |
| 177,4230202 | -1,578467753 | 0,342013889 | -4,615215355 | 3,92688E-06 | 0,000496411 | ENSG00000188322 | SBK1 |
| 202,7576103 | -1,571512505 | 0,340536236 | -4,61481727 | 3,93441E-06 | 0,000496411 | ENSG00000111186 | WNT5B |
| 770,5292558 | -1,629760299 | 0,352794011 | -4,619580405 | 3,84517E-06 | 0,000492502 | ENSG00000134539 | KLRD1 |
| 668,7616886 | 1,104143496 | 0,238572695 | 4,628121826 | 3,68997E-06 | 0,000477551 | ENSG00000118308 | IRAG2 |
| 367,5298545 | 1,199759158 | 0,259264526 | 4,627548458 | 3,7002E-06 | 0,000477551 | ENSG00000173208 | ABCD2 |
| 235,9903897 | 1,641205188 | 0,354424571 | 4,630619098 | 3,64574E-06 | 0,000477551 | ENSG00000128271 | ADORA2A |
| 95,15307419 | 2,523546971 | 0,545177733 | 4,628851877 | 3,67699E-06 | 0,000477551 | ENSG00000134909 | ARHGAP32 |
| 516,9563524 | 1,361553964 | 0,291081651 | 4,677567133 | 2,90299E-06 | 0,000386463 | ENSG00000153317 | ASAP1 |
| 506,8741008 | 1,277657838 | 0,272762783 | 4,68413551 | 2,81144E-06 | 0,000377246 | ENSG00000137265 | IRF4 |
| 1588,588816 | 1,279445632 | 0,272408435 | 4,696791542 | 2,6428E-06 | 0,000357454 | ENSG00000184730 | APOBR |
| 235,2011055 | -1,411257926 | 0,299686203 | -4,709118775 | 2,4879E-06 | 0,000339217 | ENSG00000139112 | GABARAPL1 |
| 416,7403988 | -1,561508312 | 0,331443118 | -4,711240709 | 2,46213E-06 | 0,000338433 | ENSG00000200312 | ENSG00000200312 |
| 115,5339318 | -1,901020263 | 0,40184309 | -4,730752648 | 2,23689E-06 | 0,000309993 | ENSG00000180549 | FUT7 |
| 225,2753306 | -1,627773148 | 0,343452106 | -4,739447281 | 2,14302E-06 | 0,000299438 | ENSG00000013619 | MAMLD1 |
| 469,3979426 | -1,568953233 | 0,330743427 | -4,743717046 | 2,09832E-06 | 0,000295636 | ENSG00000113088 | GZMK |
| 1474,830549 | 1,030290384 | 0,217085911 | 4,746002997 | 2,07476E-06 | 0,000294772 | ENSG00000144354 | CDCA7 |
| 357,5739952 | 1,672679985 | 0,348894214 | 4,794232519 | 1,63299E-06 | 0,000233974 | ENSG00000157483 | MYO1E |
| 2769,841565 | -1,428623033 | 0,29763089 | -4,799982404 | 1,5868E-06 | 0,000229299 | ENSG00000202198 | RN7SK |
| 43,61147676 | -3,244867136 | 0,674960081 | -4,807494885 | 1,52833E-06 | 0,000222755 | ENSG00000108924 | HLF |
| 817,2625188 | -1,25918159 | 0,261817566 | -4,809385447 | 1,51395E-06 | 0,000222577 | ENSG00000133816 | MICAL2 |
| 254,4986254 | 1,955620965 | 0,406341079 | 4,812757231 | 1,48862E-06 | 0,000220773 | ENSG00000141469 | SLC14A1 |
| 1438,356876 | 1,290777942 | 0,267770983 | 4,820454876 | 1,43231E-06 | 0,000214302 | ENSG00000142102 | PGGHG |
| 1353,62226 | -1,043877661 | 0,216464698 | -4,822392154 | 1,41847E-06 | 0,000214125 | ENSG00000172005 | MAL |
| 689,8028282 | 1,314698351 | 0,272467008 | 4,825165291 | 1,39887E-06 | 0,00021307 | ENSG00000134352 | IL6ST |
| 207,3580632 | 1,838548883 | 0,379747874 | 4,841498813 | 1,28863E-06 | 0,000198063 | ENSG00000138119 | MYOF |
| 53,30768581 | 2,816338411 | 0,580870564 | 4,848478453 | 1,24412E-06 | 0,000192976 | ENSG00000150681 | RGS18 |
| 722,5969984 | 1,220013201 | 0,249788784 | 4,884179278 | 1,03861E-06 | 0,00016259 | ENSG00000106415 | GLCCI1 |
| 221,9205588 | -1,545815371 | 0,316317951 | -4,886903721 | 1,02434E-06 | 0,000161856 | ENSG00000155657 | TTN |
| 784,0605026 | 1,119891097 | 0,228927852 | 4,891895367 | 9,98695E-07 | 0,000159292 | ENSG00000109171 | SLAIN2 |
| 35,12422779 | -3,218339849 | 0,649679697 | -4,953733147 | 7,2803E-07 | 0,000117227 | ENSG00000087842 | PIR |
| 644,1893265 | 1,165749938 | 0,235039524 | 4,95980386 | 7,05644E-07 | 0,000114715 | ENSG00000085719 | CPNE3 |
| 1564,279246 | 1,147920408 | 0,230791322 | 4,973845632 | 6,56376E-07 | 0,000107741 | ENSG00000198826 | ARHGAP11A |
| 5780,518718 | 1,083036235 | 0,217068538 | 4,989374522 | 6,05751E-07 | 0,000100406 | ENSG00000115306 | SPTBN1 |
| 12028,44123 | 1,00553774 | 0,199092487 | 5,050606163 | 4,4041E-07 | 7,37229E-05 | ENSG00000125354 | SEPTIN6 |

| | | | | | | | |
|---|---|---|---|---|---|---|---|
| 143,9093559 | -2,663921549 | 0,526121387 | -5,063321151 | 4,12015E-07 | 6,96594E-05 | ENSG00000164530 | PI16 |
| 229,2100569 | -1,730540172 | 0,341032718 | -5,074410987 | 3,88698E-07 | 6,6381E-05 | ENSG00000189430 | NCR1 |
| 1661,868819 | 1,261217866 | 0,24829354 | 5,07954361 | 3,78343E-07 | 6,52718E-05 | ENSG00000129250 | KIF1C |
| 138,854804 | 1,945493006 | 0,382047399 | 5,092281778 | 3,5378E-07 | 6,16635E-05 | ENSG00000196814 | MVB12B |
| 69,72693333 | -2,36741053 | 0,464482141 | -5,096881707 | 3,45294E-07 | 6,08113E-05 | ENSG00000177301 | KCNA2 |
| 119,2671693 | -2,712264287 | 0,526911164 | -5,147479254 | 2,6401E-07 | 4,69855E-05 | ENSG00000203747 | FCGR3A |
| 935,9483004 | 1,256198169 | 0,243670995 | 5,155304475 | 2,53219E-07 | 4,55444E-05 | ENSG00000149212 | SESN3 |
| 1088,257131 | -1,410529857 | 0,270230838 | -5,219722017 | 1,79192E-07 | 3,25763E-05 | ENSG00000033327 | GAB2 |
| 565,8095462 | 1,618885031 | 0,309779485 | 5,225927187 | 1,73285E-07 | 3,19765E-05 | ENSG00000004468 | CD38 |
| 26,36437896 | 5,734113436 | 1,09740365 | 5,225163445 | 1,74001E-07 | 3,19765E-05 | ENSG00000250548 | LINC01303 |
| 253,5448879 | -1,808562458 | 0,345384158 | -5,236379312 | 1,63757E-07 | 3,07627E-05 | ENSG00000169896 | ITGAM |
| 41,80069812 | -3,39509784 | 0,6469919 | -5,247512129 | 1,54167E-07 | 2,92865E-05 | ENSG00000081189 | MEF2C |
| 121,5180149 | 2,025504412 | 0,384178437 | 5,272301135 | 1,34724E-07 | 2,58838E-05 | ENSG00000266208 | ENSG00000266208 |
| 660,8223629 | -1,734134232 | 0,326817589 | -5,306122713 | 1,11982E-07 | 2,17618E-05 | ENSG00000168298 | H1-4 |
| 1400,294975 | 1,179731072 | 0,218013876 | 5,411265978 | 6,25807E-08 | 1,23029E-05 | ENSG00000100628 | ASB2 |
| 401,8347249 | 1,424215363 | 0,262541885 | 5,42471675 | 5,80465E-08 | 1,15458E-05 | ENSG00000106723 | SPIN1 |
| 172,3487418 | -1,779731933 | 0,327729053 | -5,430497891 | 5,6197E-08 | 1,1311E-05 | ENSG00000114757 | PEX5L |
| 1101,702951 | 1,720476851 | 0,316558321 | 5,434944332 | 5,48135E-08 | 1,11655E-05 | ENSG00000138795 | LEF1 |
| 335,7377749 | 1,714219281 | 0,312059844 | 5,493238913 | 3,94628E-08 | 8,13656E-06 | ENSG00000179934 | CCR8 |
| 417,1584389 | 2,007472019 | 0,36519052 | 5,497054023 | 3,86188E-08 | 8,06085E-06 | ENSG00000234857 | HNRNPUL2-BSCL2 |
| 150,177025 | -1,945932181 | 0,353408366 | -5,506185959 | 3,66691E-08 | 7,74956E-06 | ENSG00000248323 | LUCAT1 |
| 170,6537015 | -1,998214128 | 0,359452433 | -5,559050221 | 2,71247E-08 | 5,83413E-06 | ENSG00000259001 | ENSG00000259001 |
| 4708,31755 | -1,141780696 | 0,20542359 | -5,558177102 | 2,72607E-08 | 5,83413E-06 | ENSG00000059804 | SLC2A3 |
| 32,77527808 | 5,511457546 | 0,986709545 | 5,585693959 | 2,32769E-08 | 5,11094E-06 | ENSG00000131142 | CCL25 |
| 71,9458765 | 2,724432265 | 0,485392182 | 5,612847433 | 1,99024E-08 | 4,4275E-06 | ENSG00000152670 | DDX4 |
| 21,78509216 | 7,862438423 | 1,399519617 | 5,617955139 | 1,93231E-08 | 4,35593E-06 | ENSG00000284829 | ENSG00000284829 |
| 201,1486363 | -1,782620576 | 0,316823931 | -5,626533856 | 1,83867E-08 | 4,20086E-06 | ENSG00000157570 | TSPAN18 |
| 494,7273231 | -1,43075316 | 0,253176528 | -5,651207755 | 1,59324E-08 | 3,69E-06 | ENSG00000146859 | TMEM140 |
| 94,54945859 | -2,731816224 | 0,482347723 | -5,663582711 | 1,48245E-08 | 3,48107E-06 | ENSG00000137441 | FGFBP2 |
| 152,0177237 | 2,211852026 | 0,390211431 | 5,668342469 | 1,44186E-08 | 3,43344E-06 | ENSG00000173801 | JUP |
| 98,66745283 | -2,782503544 | 0,490271558 | -5,675433341 | 1,38338E-08 | 3,34126E-06 | ENSG00000120659 | TNFSF11 |
| 301,5761769 | 1,608831601 | 0,280564829 | 5,734259728 | 9,79392E-09 | 2,39979E-06 | ENSG00000151136 | ABTB3 |
| 85,26074223 | -2,880315027 | 0,501647681 | -5,74170904 | 9,37258E-09 | 2,33033E-06 | ENSG00000276070 | CCL4L2 |
| 632,951699 | 1,581895283 | 0,275465002 | 5,742636165 | 9,32139E-09 | 2,33033E-06 | ENSG00000228168 | HNRNPA1P21 |
| 5193,230461 | -1,273475201 | 0,220890506 | -5,765187585 | 8,15671E-09 | 2,08948E-06 | ENSG00000140564 | FURIN |
| 366,761458 | 1,62937736 | 0,28254024 | 5,766886022 | 8,07496E-09 | 2,08948E-06 | ENSG00000146285 | SCML4 |
| 141,9920778 | -2,006872416 | 0,346257908 | -5,795889041 | 6,79602E-09 | 1,79532E-06 | ENSG00000184293 | ENSG00000184293 |
| 456,4243955 | 1,530413186 | 0,261201912 | 5,859119396 | 4,65328E-09 | 1,24878E-06 | ENSG00000131697 | NPHP4 |
| 84,67215757 | -2,873498592 | 0,487548 | -5,89377578 | 3,77469E-09 | 1,03525E-06 | ENSG00000090382 | LYZ |
| 359,0300605 | -1,586962508 | 0,26908741 | -5,897572493 | 3,68888E-09 | 1,03525E-06 | ENSG00000121064 | SCPEP1 |
| 649,745211 | 1,988239044 | 0,337399729 | 5,892829409 | 3,79638E-09 | 1,03525E-06 | ENSG00000145287 | PLAC8 |
| 360,7992974 | -1,64902695 | 0,277541885 | -5,941542654 | 2,82352E-09 | 8,09107E-07 | ENSG00000107281 | NPDC1 |
| 722,5968356 | -1,571401926 | 0,258653315 | -6,075321043 | 1,2374E-09 | 3,60702E-07 | ENSG00000171476 | HOPX |
| 37,29474033 | -5,404966934 | 0,884834175 | -6,108451827 | 1,00602E-09 | 2,984E-07 | ENSG00000196782 | MAML3 |
| 470,5156967 | -1,760029543 | 0,284755385 | -6,180847266 | 6,37585E-10 | 1,92494E-07 | ENSG00000235831 | BHLHE40-AS1 |
| 1244,903018 | -2,150987244 | 0,347790489 | -6,184721301 | 6,22123E-10 | 1,91241E-07 | ENSG00000139289 | PHLDA1 |
| 6487,560897 | 1,910533519 | 0,308163054 | 6,199748779 | 5,65534E-10 | 1,77064E-07 | ENSG00000081059 | TCF7 |
| 130,5257104 | -3,139291322 | 0,504823416 | -6,21859292 | 5,01633E-10 | 1,60021E-07 | ENSG00000177508 | IRX3 |
| 255,9683899 | -2,156642431 | 0,343802325 | -6,272914034 | 3,54353E-10 | 1,15212E-07 | ENSG00000163106 | HPGDS |
| 732,1053229 | 1,714281145 | 0,273265642 | 6,273313892 | 3,53443E-10 | 1,15212E-07 | ENSG00000152229 | PSTPIP2 |
| 269,8526606 | -2,414432196 | 0,383416878 | -6,297146359 | 3,03175E-10 | 1,02516E-07 | ENSG00000107968 | MAP3K8 |
| 351,6690275 | -1,81039005 | 0,286189173 | -6,325850937 | 2,51841E-10 | 8,68954E-08 | ENSG00000167851 | CD300A |
| 86,47044927 | -2,743322638 | 0,429905772 | -6,381218429 | 1,75685E-10 | 6,18812E-08 | ENSG00000171954 | CYP4F22 |
| 101,3166725 | 3,086701376 | 0,482427707 | 6,398267203 | 1,5715E-10 | 5,65306E-08 | ENSG00000129226 | CD68 |
| 206,7189641 | 2,206279888 | 0,344244472 | 6,409049587 | 1,4643E-10 | 5,38192E-08 | ENSG00000255026 | ENSG00000255026 |
| 586,8343345 | 1,713143243 | 0,266498683 | 6,428336631 | 1,29008E-10 | 4,84697E-08 | ENSG00000232653 | GOLGA8N |
| 169,1335264 | -2,208187658 | 0,340018603 | -6,494314246 | 8,34125E-11 | 3,20512E-08 | ENSG00000178860 | MSC |
| 2571,16348 | -2,004096257 | 0,304987383 | -6,571079233 | 4,99519E-11 | 1,96404E-08 | ENSG00000105374 | NKG7 |
| 414,8247484 | -1,917472392 | 0,29008411 | -6,610056625 | 3,84173E-11 | 1,54648E-08 | ENSG00000126822 | PLEKHG3 |
| 532,3102739 | 1,865707096 | 0,282207575 | 6,611116286 | 3,81433E-11 | 1,54648E-08 | ENSG00000110841 | PPFIBP1 |
| 2147,057304 | 1,503875777 | 0,222534754 | 6,757936679 | 1,39971E-11 | 5,91621E-09 | ENSG00000137312 | FLOT1 |
| 386,1921476 | 1,758906621 | 0,259545445 | 6,776873389 | 1,22804E-11 | 5,32373E-09 | ENSG00000177374 | HIC1 |
| 85,4527136 | -3,444708381 | 0,50769199 | -6,785035899 | 1,16058E-11 | 5,16365E-09 | ENSG00000111536 | IL26 |
| 242,9566174 | -2,2385652 | 0,329066656 | -6,802771284 | 1,02626E-11 | 4,68943E-09 | ENSG00000186891 | TNFRSF18 |
| 589,2763775 | -2,503353658 | 0,365283989 | -6,853171051 | 7,22306E-12 | 3,39223E-09 | ENSG00000275302 | CCL4 |
| 1144,588789 | -1,645412351 | 0,238608336 | -6,895871187 | 5,35357E-12 | 2,58608E-09 | ENSG00000280800 | ENSG00000280800 |
| 366,9846105 | -2,091130323 | 0,296245362 | -7,058778265 | 1,67973E-12 | 8,3527E-10 | ENSG00000172215 | CXCR6 |
| 623,0514486 | 1,983338834 | 0,27904335 | 7,107636981 | 1,18047E-12 | 6,04792E-10 | ENSG00000077782 | FGFR1 |
| 228,7688076 | 2,317041861 | 0,309576539 | 7,484552515 | 7,17913E-14 | 3,79305E-11 | ENSG00000116574 | RHOU |
| 5350,047252 | -2,11973307 | 0,282896767 | -7,49295614 | 6,73394E-14 | 3,6726E-11 | ENSG00000225783 | MIAT |
| 96,21809033 | -3,925073856 | 0,517902845 | -7,578784121 | 3,48808E-14 | 1,96577E-11 | ENSG00000236481 | LINC02195 |
| 582,1233193 | -2,087418671 | 0,273901252 | -7,621062891 | 2,51595E-14 | 1,4668E-11 | ENSG00000163508 | EOMES |
| 7679,690715 | -1,874776062 | 0,24469277 | -7,661755046 | 1,83409E-14 | 1,10746E-11 | ENSG00000274012 | RN7SL2 |
| 7442,65403 | -2,033249984 | 0,257535997 | -7,895012755 | 2,90286E-15 | 1,81773E-12 | ENSG00000276168 | RN7SL1 |
| 230,0212331 | 2,66915553 | 0,337155622 | 7,916687007 | 2,43923E-15 | 1,58615E-12 | ENSG00000070214 | SLC44A1 |

| | | | | | | | |
|---|---|---|---|---|---|---|---|
| 607,4300004 | 1,948693272 | 0,242998954 | 8,019348382 | 1,06308E-15 | 7,18937E-13 | ENSG00000112303 | VNN2 |
| 1180,443984 | -1,981927712 | 0,238477247 | -8,310762309 | 9,50892E-17 | 6,69864E-14 | ENSG00000172965 | MIR4435-2HG |
| 188,7740081 | 3,208259712 | 0,385070266 | 8,331621511 | 7,97425E-17 | 5,86177E-14 | ENSG00000131016 | AKAP12 |
| 178,3904367 | 2,817755807 | 0,334233266 | 8,430506765 | 3,44171E-17 | 2,64496E-14 | ENSG00000005102 | MEOX1 |
| 144,3503383 | -3,034228512 | 0,358229349 | -8,470072364 | 2,45241E-17 | 2,07038E-14 | ENSG00000175779 | LINC02694 |
| 4275,123991 | -2,525203946 | 0,298327267 | -8,464542886 | 2,57159E-17 | 2,07038E-14 | ENSG00000100453 | GZMB |
| 2558,061645 | -1,893053212 | 0,220728153 | -8,576401267 | 9,78876E-18 | 8,71045E-15 | ENSG00000077984 | CST7 |
| 1129,228599 | -2,093198507 | 0,243298954 | -8,603401185 | 7,73879E-18 | 7,43225E-15 | ENSG00000222041 | CYTOR |
| 221,9885195 | 2,860678999 | 0,332604191 | 8,600850741 | 7,91273E-18 | 7,43225E-15 | ENSG00000133026 | MYH10 |
| 1516,576117 | -2,172955672 | 0,249789023 | -8,699163988 | 3,34338E-18 | 3,53291E-15 | ENSG00000282885 | ENSG00000282885 |
| 470,8812287 | -2,490507832 | 0,280466358 | -8,879880802 | 6,6932E-19 | 7,54413E-16 | ENSG00000113525 | IL5 |
| 141,8038371 | -3,905695074 | 0,427813497 | -9,129433977 | 6,88584E-20 | 8,31563E-17 | ENSG00000116667 | C1orf21 |
| 918,7778703 | -3,155404893 | 0,333068141 | -9,473751784 | 2,69969E-21 | 3,51105E-18 | ENSG00000169194 | IL13 |
| 261,1566841 | -3,125763664 | 0,32552172 | -9,602319819 | 7,8165E-22 | 1,10128E-18 | ENSG00000100450 | GZMH |
| 343,3736583 | -2,775285747 | 0,287165157 | -9,6644237 | 4,27016E-22 | 6,56324E-19 | ENSG00000115956 | PLEK |
| 184,497159 | -4,893457741 | 0,498007446 | -9,826073437 | 8,69428E-23 | 1,46994E-19 | ENSG00000138678 | GPAT3 |
| 696,3438061 | -2,448347483 | 0,24641709 | -9,935786027 | 2,90873E-23 | 5,46421E-20 | ENSG00000136161 | RCBTB2 |
| 3863,496235 | -2,071752222 | 0,208002845 | -9,96021098 | 2,27579E-23 | 4,8096E-20 | ENSG00000132965 | ALOX5AP |
| 438,2147369 | -3,643466036 | 0,343659058 | -10,60197877 | 2,91741E-26 | 7,04638E-23 | ENSG00000205336 | ADGRG1 |
| 988,985833 | -2,478475811 | 0,225423696 | -10,9947439 | 4,05062E-28 | 1,1414E-24 | ENSG00000016391 | CHDH |
| 1189,552284 | -2,624907399 | 0,225234181 | -11,65412544 | 2,18613E-31 | 7,39219E-28 | ENSG00000139679 | LPAR6 |
| 253,8962399 | -3,845317312 | 0,325777966 | -11,80349108 | 3,74448E-32 | 1,5827E-28 | ENSG00000073756 | PTGS2 |
| 16787,49505 | -2,790565533 | 0,210858634 | -13,23429581 | 5,56155E-40 | 3,13431E-36 | ENSG00000115523 | GNLY |
| 1126,192212 | -4,464096168 | 0,243393996 | -18,34102831 | 3,89362E-75 | 3,29147E-71 | ENSG00000056736 | IL17RB |
| 7150,295401 | 6,600984496 | 0,315409651 | 20,92828954 | 2,95926E-97 | 5,00322E-93 | ENSG00000112182 | BACH2 |

LV_STAT5B_vs_GFP_d12

| baseMean | log2FoldChange | lfcSE | stat | pvalue | padj | ENSEMBL | SYMBOL |
|---|---|---|---|---|---|---|---|
| 6,821668232 | -29,99494959 | 3,965159363 | -7,5646265 | 3,88981E-14 | 7,93132E-11 | ENSG00000260966 | ENSG00000260966 |
| 27,15146104 | -29,10442725 | 3,908427719 | -7,446581937 | 9,57896E-14 | 1,59092E-10 | ENSG00000124208 | PEDS1-UBE2V1 |
| 26,18444987 | -28,55442203 | 3,907990036 | -7,306677287 | 2,73829E-13 | 4,11591E-10 | ENSG00000278996 | ENSG00000278996 |
| 10,98906546 | -26,75200464 | 3,911162846 | -6,839910711 | 7,92426E-12 | 7,83397E-09 | ENSG00000261499 | ENSG00000261499 |
| 2,276037345 | -25,68714647 | 2,946268326 | -8,718536004 | 2,81822E-18 | 9,19416E-15 | ENSG00000239945 | ENSG00000239945 |
| 3,854614013 | -25,16556199 | 2,547630456 | -9,878026827 | 5,18437E-23 | 2,81891E-19 | ENSG00000257921 | ENSG00000257921 |
| 0,929991374 | -23,74923453 | 3,989398625 | -5,953086356 | 2,63132E-09 | 1,53293E-06 | ENSG00000131864 | USP29 |
| 1,048146595 | -23,59588053 | 3,80506959 | -6,20116925 | 5,60452E-10 | 3,97482E-07 | ENSG00000104369 | JPH1 |
| 3,976276363 | -23,58220728 | 2,547605417 | -9,256616871 | 2,11012E-20 | 7,64896E-17 | ENSG00000234769 | ENSG00000234769 |
| 1,083258448 | -23,51081387 | 3,803622637 | -6,181163621 | 6,36308E-10 | 4,32477E-07 | ENSG00000138650 | PCDH10 |
| 2,690033816 | -23,3539816 | 2,811864615 | -8,305514241 | 9,93881E-17 | 2,94767E-13 | ENSG00000267827 | ENSG00000267827 |
| 1,627484905 | -23,05601744 | 3,312015674 | -6,961324979 | 3,37087E-12 | 3,92755E-09 | ENSG00000254959 | INMT-MINDY4 |
| 1,112265938 | -23,01130543 | 3,786084119 | -6,077864282 | 1,21794E-09 | 7,49698E-07 | ENSG00000230646 | KLF2P2 |
| 0,802106307 | -22,89282494 | 4,012195142 | -5,705810443 | 1,15791E-08 | 5,39651E-06 | ENSG00000233087 | RAB6D |
| 0,894713277 | -22,6911314 | 3,988919458 | -5,688540879 | 1,28129E-08 | 5,88746E-06 | ENSG00000229111 | MED4-AS1 |
| 0,917865019 | -22,5872205 | 3,989834612 | -5,661192179 | 1,50325E-08 | 6,48594E-06 | ENSG00000265630 | ENSG00000265630 |
| 0,833886928 | -22,51116886 | 3,998047002 | -5,630541325 | 1,79645E-08 | 7,51376E-06 | ENSG00000070731 | ST6GALNAC2 |
| 0,877170861 | -22,47051443 | 3,986186432 | -5,63709571 | 1,72942E-08 | 7,32735E-06 | ENSG00000251660 | ENSG00000251660 |
| 2,324940642 | -20,68559993 | 2,732404609 | -7,570474687 | 3,71863E-14 | 7,93132E-11 | ENSG00000258830 | ENSG00000258830 |
| 0,742249989 | -19,35893544 | 3,974855595 | -4,870349369 | 1,11401E-06 | 0,000306504 | ENSG00000225756 | DBH-AS1 |
| 0,741356787 | -19,20372261 | 3,998388136 | -4,802866044 | 1,56411E-06 | 0,000384517 | ENSG00000225721 | ENSG00000225721 |
| 4,636993713 | -19,1323563 | 3,968356652 | -4,821229033 | 1,42676E-06 | 0,000363647 | ENSG00000267303 | ENSG00000267303 |
| 0,636013357 | -19,01642421 | 3,975514829 | -4,783386563 | 1,72366E-06 | 0,000413476 | ENSG00000253859 | ENSG00000253859 |
| 0,636013357 | -19,01642421 | 3,975514829 | -4,783386563 | 1,72366E-06 | 0,000413476 | ENSG00000264673 | ENSG00000264673 |
| 0,822243517 | -18,87785519 | 3,883843177 | -4,860612112 | 1,17023E-06 | 0,000312932 | ENSG00000231345 | ENSG00000231345 |
| 2,392357119 | -18,85245765 | 2,91141652 | -6,475355732 | 9,45888E-11 | 7,5265E-08 | ENSG00000218336 | TENM3 |
| 3,214481185 | -18,80321708 | 2,681762084 | -7,011515747 | 2,3575E-12 | 3,01056E-09 | ENSG00000249590 | ENSG00000249590 |
| 3,239656049 | -18,15169316 | 2,666967454 | -6,806117236 | 1,00268E-11 | 9,62101E-09 | ENSG00000285628 | ENSG00000285628 |
| 3,232347997 | -18,13469735 | 2,626501535 | -6,904506663 | 5,03782E-12 | 5,47846E-09 | ENSG00000265018 | ENSG00000265018 |
| 1,62147037 | -18,10673747 | 3,277923981 | -5,523843009 | 3,31664E-08 | 1,27296E-05 | ENSG00000272760 | ENSG00000272760 |
| 0,624648438 | -18,07645829 | 3,998186757 | -4,521164064 | 6,15005E-06 | 0,001166507 | ENSG00000182272 | B4GALNT4 |
| 0,624648438 | -18,07645829 | 3,998186757 | -4,521164064 | 6,15005E-06 | 0,001166507 | ENSG00000288570 | LOC128092248 |
| 0,927394141 | -17,91087896 | 4,022500732 | -4,452672641 | 8,4808E-06 | 0,001487514 | ENSG00000019186 | CYP24A1 |
| 0,927394141 | -17,91087896 | 4,022500732 | -4,452672641 | 8,4808E-06 | 0,001487514 | ENSG00000244528 | ENSG00000244528 |
| 1,803608675 | -17,78757126 | 3,191565165 | -5,573306618 | 2,49949E-08 | 1,00671E-05 | ENSG00000186076 | ENSG00000186076 |
| 1,413059517 | -17,7782069 | 3,46903125 | -5,124833309 | 2,97801E-07 | 9,43249E-05 | ENSG00000214914 | RPL23AP3 |
| 1,15196441 | -17,77226146 | 3,721668678 | -4,775347566 | 1,79397E-06 | 0,000424105 | ENSG00000270025 | BMS1P7 |
| 1,02251036 | -17,74740817 | 3,882213062 | -4,571466813 | 4,84322E-06 | 0,000969357 | ENSG00000207417 | ENSG00000207417 |
| 0,833928863 | -17,60753923 | 3,989293525 | -4,413698596 | 1,01619E-05 | 0,001726683 | ENSG00000124116 | WFDC3 |
| 0,931558443 | -17,48435548 | 3,989540577 | -4,382548601 | 1,17299E-05 | 0,001903862 | ENSG00000271554 | ENSG00000271554 |
| 0,895243317 | -17,46487377 | 3,989820971 | -4,377357755 | 1,20127E-05 | 0,001921085 | ENSG00000264350 | ENSG00000264350 |
| 1,287979481 | -17,44174851 | 3,585651896 | -4,86431729 | 1,14853E-06 | 0,000309665 | ENSG00000220392 | FCF1P5 |
| 1,546887581 | -17,43337512 | 3,361220105 | -5,186621102 | 2,14144E-07 | 6,91706E-05 | ENSG00000232702 | ENSG00000232702 |
| 0,73088507 | -17,42214782 | 3,994171967 | -4,361892269 | 1,28942E-05 | 0,002032181 | ENSG00000269082 | ENSG00000269082 |
| 1,366856491 | -17,30430154 | 3,30290887 | -5,239109593 | 1,61353E-07 | 5,37142E-05 | ENSG00000233179 | ENSG00000233179 |
| 1,44021381 | -17,27895768 | 3,276790615 | -5,273134512 | 1,34113E-07 | 4,60559E-05 | ENSG00000167711 | SERPINF2 |
| 0,748122433 | -17,27195464 | 3,996676943 | -4,321578874 | 1,54917E-05 | 0,002339815 | ENSG00000276601 | ENSG00000276601 |
| 0,748122433 | -17,27195464 | 3,996676943 | -4,321578874 | 1,54917E-05 | 0,002339815 | ENSG00000286454 | ENSG00000286454 |
| 0,761002448 | -17,13695168 | 3,994614697 | -4,290013676 | 1,78662E-05 | 0,002613755 | ENSG00000177151 | OR2T35 |
| 0,640992599 | -17,12164074 | 4,002324118 | -4,277924584 | 1,88644E-05 | 0,002711154 | ENSG00000231291 | ENSG00000231291 |
| 0,943358334 | -17,03731663 | 3,990390713 | -4,269586078 | 1,95836E-05 | 0,002760261 | ENSG00000200708 | ENSG00000200708 |
| 0,717713496 | -16,8850921 | 3,999908323 | -4,221369777 | 2,42822E-05 | 0,003301582 | ENSG00000222067 | RNU4-86P |
| 1,185679724 | -16,83928956 | 3,486133036 | -4,830363439 | 1,36284E-06 | 0,000350089 | ENSG00000229806 | RPS15P5 |
| 0,633735404 | -16,83192674 | 4,002628627 | -4,205218198 | 2,6083E-05 | 0,003459078 | ENSG00000235782 | ENSG00000235782 |
| 0,926358818 | -16,70050171 | 4,008158087 | -4,1666275 | 3,09139E-05 | 0,004006105 | ENSG00000234975 | FTH1P2 |
| 0,734077908 | -16,6492799 | 3,999453998 | -4,162888211 | 3,14247E-05 | 0,004052175 | ENSG00000276204 | ENSG00000276204 |
| 0,642896237 | -16,62250076 | 4,013186618 | -4,141970544 | 3,44335E-05 | 0,004354097 | ENSG00000268391 | MTCO3P42 |
| 0,642896237 | -16,62250076 | 4,013186618 | -4,141970544 | 3,44335E-05 | 0,004354097 | ENSG00000273204 | ENSG00000273204 |
| 0,674424544 | -16,50855983 | 4,012027918 | -4,114766938 | 3,87571E-05 | 0,004771362 | ENSG00000257083 | LOC102724421 |
| 0,949744011 | -16,41161147 | 4,008758427 | -4,093938751 | 4,24106E-05 | 0,005068149 | ENSG00000271672 | ENSG00000271672 |
| 0,631034115 | -16,36509037 | 4,013731003 | -4,07727632 | 4,55663E-05 | 0,005405658 | ENSG00000250274 | LOC100128059 |
| 0,595233272 | -16,2426715 | 4,041102813 | -4,019366061 | 5,83549E-05 | 0,006410006 | ENSG00000267046 | ENSG00000267046 |
| 0,639206196 | -16,23224383 | 4,018012536 | -4,039868886 | 5,34811E-05 | 0,006016438 | ENSG00000223561 | LINC03007 |
| 0,639206196 | -16,23224383 | 4,018012536 | -4,039868886 | 5,34811E-05 | 0,006016438 | ENSG00000271916 | ENSG00000271916 |
| 0,639206196 | -16,23224383 | 4,018012536 | -4,039868886 | 5,34811E-05 | 0,006016438 | ENSG00000284640 | LOC105378644 |
| 0,532969564 | -16,2212163 | 4,021658002 | -4,033464879 | 5,49604E-05 | 0,00611955 | ENSG00000200999 | LOC124900179 |
| 0,532969564 | -16,2212163 | 4,021658002 | -4,033464879 | 5,49604E-05 | 0,00611955 | ENSG00000215223 | ENSG00000215223 |
| 0,85167946 | -16,19233321 | 4,013528765 | -4,034438061 | 5,47331E-05 | 0,00611955 | ENSG00000280058 | ENSG00000280058 |
| 1,374690539 | -15,96846169 | 3,507957263 | -4,552068482 | 5,3121E-06 | 0,001025456 | ENSG00000234965 | SHISA8 |
| 56,59434959 | -9,306753492 | 1,246021914 | -7,469173203 | 8,07003E-14 | 1,46265E-10 | ENSG00000256500 | ENSG00000256500 |
| 12,89936403 | -5,893792691 | 1,522983912 | -3,869898195 | 0,000108881 | 0,01066705 | ENSG00000133863 | TEX15 |
| 5,686680258 | -5,859845264 | 1,692878417 | -3,461468471 | 0,000537237 | 0,036669438 | ENSG00000168229 | PTGDR |
| 7,33841842 | -5,813143929 | 1,564102185 | -3,716601117 | 0,000201921 | 0,017566576 | ENSG00000224837 | ENSG00000224837 |

| | | | | | | | |
|---|---|---|---|---|---|---|---|
| 12,46123058 | -5,668777459 | 1,301116395 | -4,356856529 | 1,31944E-05 | 0,002052566 | ENSG00000167768 | KRT1 |
| 7,244904657 | -5,495627423 | 1,575207266 | -3,488828132 | 0,000485143 | 0,034332547 | ENSG00000266770 | RN7SL619P |
| 8,452532214 | -5,389797008 | 1,562253714 | -3,450013887 | 0,000560558 | 0,037551612 | ENSG00000185736 | ADARB2 |
| 12,47356482 | -5,369504892 | 1,451781657 | -3,698562291 | 0,000216824 | 0,018517463 | ENSG00000203761 | ENSG00000203761 |
| 14,90427021 | -4,590600396 | 1,004822323 | -4,568569278 | 4,91065E-06 | 0,00097686 | ENSG00000006468 | ETV1 |
| 48,85807163 | -4,441774283 | 0,633719222 | -7,009057216 | 2,39929E-12 | 3,01056E-09 | ENSG00000041353 | RAB27B |
| 14,40522793 | -4,128196548 | 1,117540262 | -3,694002524 | 0,000220752 | 0,018745192 | ENSG00000198618 | ENSG00000198618 |
| 9,748060079 | -4,025537211 | 1,200904936 | -3,352086488 | 0,00080205 | 0,048913266 | ENSG00000122786 | CALD1 |
| 18,37345045 | -3,680552079 | 1,020095211 | -3,608047601 | 0,00030851 | 0,023535825 | ENSG00000118785 | SPP1 |
| 32,98429971 | -3,175638534 | 0,745775768 | -4,258168031 | 2,06109E-05 | 0,002873547 | ENSG00000082438 | COBLL1 |
| 37,10139095 | -2,693024874 | 0,60098511 | -4,481017634 | 7,4288E-06 | 0,001331632 | ENSG00000255036 | SUGT1P4-STRA6LP-CCDC180 |
| 28,47694252 | -2,676727788 | 0,633133835 | -4,227744026 | 2,36046E-05 | 0,003235617 | ENSG00000114812 | VIPR1 |
| 34,48280921 | -2,626377485 | 0,677312257 | -3,877646474 | 0,000105472 | 0,010364196 | ENSG00000114757 | PEX5L |
| 74,40646456 | -2,553921066 | 0,405708749 | -6,294961768 | 3,07476E-10 | 2,33281E-07 | ENSG00000124212 | PTGIS |
| 43,24462263 | -2,341081419 | 0,547871478 | -4,273048538 | 1,92818E-05 | 0,002758995 | ENSG00000204172 | AGAP9 |
| 257,5150894 | -2,27132054 | 0,346170192 | -6,561282837 | 5,33468E-11 | 4,57997E-08 | ENSG00000137959 | IFI44L |
| 73,08098601 | -2,121903693 | 0,435544702 | -4,871839064 | 1,10564E-06 | 0,000306504 | ENSG00000120885 | CLU |
| 53,40107271 | -2,054453934 | 0,613026378 | -3,351330396 | 0,000804243 | 0,048913266 | ENSG00000179978 | ENSG00000179978 |
| 147,5375274 | -2,000183837 | 0,319615097 | -6,258101879 | 3,89691E-10 | 2,82517E-07 | ENSG00000121989 | ACVR2A |
| 337,3550329 | -1,968341871 | 0,264412675 | -7,444203922 | 9,75307E-14 | 1,59092E-10 | ENSG00000091409 | ITGA6 |
| 58,22938032 | -1,932041371 | 0,475239973 | -4,06540165 | 4,79498E-05 | 0,005640046 | ENSG00000184979 | USP18 |
| 162,6582109 | -1,9217493 | 0,284801814 | -6,747672257 | 1,50236E-11 | 1,40037E-08 | ENSG00000134326 | CMPK2 |
| 121,5641776 | -1,910513192 | 0,359479594 | -5,314663827 | 1,06854E-07 | 3,74841E-05 | ENSG00000162630 | B3GALT2 |
| 255,5326719 | -1,907921073 | 0,333830139 | -5,715245126 | 1,09546E-08 | 5,17946E-06 | ENSG00000245164 | LINC00861 |
| 44,97825579 | -1,867535089 | 0,519426317 | -3,59538019 | 0,000323918 | 0,024459862 | ENSG00000259205 | ENSG00000259205 |
| 35,75267111 | -1,862686158 | 0,555475964 | -3,353315494 | 0,000798496 | 0,048874566 | ENSG00000126368 | NR1D1 |
| 186,472725 | -1,823931423 | 0,316406465 | -5,764520093 | 8,18905E-09 | 4,11015E-06 | ENSG00000137965 | IFI44 |
| 35,65773115 | -1,818827137 | 0,509578559 | -3,569277212 | 0,000357967 | 0,026602122 | ENSG00000055732 | MCOLN3 |
| 105,3467557 | -1,797606796 | 0,419594477 | -4,28415266 | 1,83437E-05 | 0,002654969 | ENSG00000196993 | NPIPB9 |
| 117,1440955 | -1,794289732 | 0,367660358 | -4,88029153 | 1,05929E-06 | 0,000297917 | ENSG00000113319 | RASGRF2 |
| 187,1767333 | -1,73440788 | 0,366492563 | -4,73245041 | 2,21826E-06 | 0,000499092 | ENSG00000185710 | SMG1P4 |
| 210,4749437 | -1,664774925 | 0,254331341 | -6,545693181 | 5,92201E-11 | 4,82999E-08 | ENSG00000171476 | HOPX |
| 655,0763391 | -1,655202087 | 0,231025381 | -7,164589806 | 7,80198E-13 | 1,06055E-09 | ENSG00000145649 | GZMA |
| 110,0004302 | -1,616687242 | 0,378797159 | -4,267949761 | 1,97278E-05 | 0,002762227 | ENSG00000171246 | NPTX1 |
| 59,51607993 | -1,588010127 | 0,414908481 | -3,82737447 | 0,000129517 | 0,012354899 | ENSG00000049246 | PER3 |
| 67,61376696 | -1,530906061 | 0,443454826 | -3,452225508 | 0,000555983 | 0,037476001 | ENSG00000143110 | C1orf162 |
| 127,0212521 | -1,510703121 | 0,416998962 | -3,622798279 | 0,000291433 | 0,022745726 | ENSG00000134321 | RSAD2 |
| 99,74927718 | -1,485522187 | 0,388304358 | -3,825664473 | 0,00013042 | 0,012404712 | ENSG00000153094 | BCL2L11 |
| 188,9234691 | -1,475363301 | 0,271810013 | -5,427921088 | 5,70142E-08 | 2,08992E-05 | ENSG00000184613 | NELL2 |
| 450,5082523 | -1,465088 | 0,233171624 | -6,283303137 | 3,31454E-10 | 2,45758E-07 | ENSG00000137628 | DDX60 |
| 288,0058433 | -1,458585378 | 0,287198198 | -5,078671767 | 3,80083E-07 | 0,000118094 | ENSG00000163629 | PTPN13 |
| 65,76958377 | -1,445861535 | 0,431350909 | -3,351938077 | 0,00080248 | 0,048913266 | ENSG00000081377 | CDC14B |
| 91,73617378 | -1,427413014 | 0,369977495 | -3,858107682 | 0,000114268 | 0,011061986 | ENSG00000111796 | KLRB1 |
| 138,0658351 | -1,424039168 | 0,324592501 | -4,387159796 | 1,1484E-05 | 0,00188269 | ENSG00000161048 | NAPEPLD |
| 99,06767164 | -1,420451151 | 0,345944144 | -4,106012991 | 4,02547E-05 | 0,00491861 | ENSG00000251079 | ENSG00000251079 |
| 412,2720656 | -1,409635889 | 0,253326723 | -5,564497389 | 2,62909E-08 | 1,04599E-05 | ENSG00000196693 | ZNF33B |
| 491,7670419 | -1,402961676 | 0,310598069 | -4,516968446 | 6,27312E-06 | 0,001176175 | ENSG00000172005 | MAL |
| 126,9116117 | -1,369431614 | 0,345020496 | -3,969131203 | 7,21352E-05 | 0,007591411 | ENSG00000246223 | LINC01550 |
| 322,767055 | -1,359212526 | 0,232007202 | -5,858492813 | 4,67087E-09 | 2,53971E-06 | ENSG00000204177 | ENSG00000204177 |
| 103,4663613 | -1,33222045 | 0,388173719 | -3,432021248 | 0,000599101 | 0,039725729 | ENSG00000030582 | GRN |
| 110,6723091 | -1,289253055 | 0,34640124 | -3,721848849 | 0,000197769 | 0,017344164 | ENSG00000204681 | GABBR1 |
| 159,4463666 | -1,247795358 | 0,307195441 | -4,061894125 | 4,86762E-05 | 0,005691795 | ENSG00000128872 | TMOD2 |
| 245,1253346 | -1,246967332 | 0,312377293 | -3,991862919 | 6,55563E-05 | 0,007058441 | ENSG00000116106 | EPHA4 |
| 1863,384783 | -1,193701823 | 0,248640119 | -4,800922024 | 1,57937E-06 | 0,000384517 | ENSG00000215252 | GOLGA8B |
| 392,8642302 | -1,17652169 | 0,2641703 | -4,453648614 | 8,44232E-06 | 0,001487514 | ENSG00000138640 | FAM13A |
| 634,7074948 | -1,167860713 | 0,202762491 | -5,75974731 | 8,424E-09 | 4,16401E-06 | ENSG00000151692 | RNF144A |
| 291,0077772 | -1,165588237 | 0,259017099 | -4,5000436 | 6,79395E-06 | 0,001224563 | ENSG00000267520 | ENSG00000267520 |
| 224,2598457 | -1,152705183 | 0,26211346 | -4,397733655 | 1,09387E-05 | 0,001811495 | ENSG00000013375 | PGM3 |
| 1109,440136 | -1,12715977 | 0,237972644 | -4,736509845 | 2,1743E-06 | 0,0004926 | ENSG00000115738 | ID2 |
| 169,3305011 | -1,105377785 | 0,277090931 | -3,989223976 | 6,62898E-05 | 0,007113943 | ENSG00000168056 | LTBP3 |
| 471,4799458 | -1,09683503 | 0,228109861 | -4,808363057 | 1,52171E-06 | 0,000378964 | ENSG00000181381 | DDX60L |
| 155,7852454 | -1,096396842 | 0,27654665 | -3,964599982 | 7,35191E-05 | 0,007687462 | ENSG00000181523 | SGSH |
| 260,0386579 | -1,079419578 | 0,292596023 | -3,689112271 | 0,000225038 | 0,018955476 | ENSG00000223705 | NSUN5P1 |
| 251,8944091 | -1,076822572 | 0,312069225 | -3,4505888 | 0,000559365 | 0,037548822 | ENSG00000272419 | ENSG00000272419 |
| 209,5142177 | -1,062112034 | 0,312852837 | -3,394925375 | 0,000686473 | 0,044259878 | ENSG00000233369 | ENSG00000233369 |
| 180,7078048 | -1,060707494 | 0,263825775 | -4,020484712 | 5,80785E-05 | 0,006401195 | ENSG00000251474 | ENSG00000251474 |
| 510,007267 | -1,056453705 | 0,28283673 | -3,735206893 | 0,000187561 | 0,016582622 | ENSG00000272333 | KMT2B |
| 424,2139536 | -1,049813632 | 0,286915174 | -3,658968666 | 0,000253232 | 0,020449135 | ENSG00000136111 | TBC1D4 |
| 567,057328 | -1,048468298 | 0,245541552 | -4,270023912 | 1,95452E-05 | 0,002760261 | ENSG00000166326 | TRIM44 |
| 111,190591 | -1,044805818 | 0,308036285 | -3,391827099 | 0,000694282 | 0,044452808 | ENSG00000089041 | P2RX7 |
| 176,4577918 | -1,044408299 | 0,289468962 | -3,608014806 | 0,000308549 | 0,023535825 | ENSG00000117408 | IPO13 |
| 164,8307866 | -1,04345209 | 0,267318318 | -3,903406615 | 9,48482E-05 | 0,009521004 | ENSG00000134539 | KLRD1 |
| 244,0570075 | -1,039798708 | 0,300393926 | -3,461450505 | 0,000537273 | 0,036669438 | ENSG00000136161 | RCBTB2 |
| 675,1558809 | -1,034255212 | 0,262907874 | -3,933907324 | 8,3576E-05 | 0,008574164 | ENSG00000080854 | IGSF9B |
| 492,558526 | -1,029478421 | 0,216792798 | -4,748674461 | 2,04754E-06 | 0,000470416 | ENSG00000073910 | FRY |

| | | | | | | | |
|---|---|---|---|---|---|---|---|
| 425,970656 | -1,028324501 | 0,215691743 | -4,767565455 | 1,86465E-06 | 0,000437644 | ENSG00000184384 | MAML2 |
| 799,348456 | -1,023035254 | 0,195619219 | -5,22972773 | 1,6976E-07 | 5,59419E-05 | ENSG00000106952 | TNFSF8 |
| 222,7600848 | -1,017532415 | 0,299706827 | -3,395092547 | 0,000686054 | 0,044259878 | ENSG00000090554 | FLT3LG |
| 193,0694045 | -1,016794338 | 0,278413625 | -3,652099774 | 0,000260105 | 0,020849284 | ENSG00000119917 | IFIT3 |
| 187,9616692 | -1,000522634 | 0,275975287 | -3,625406626 | 0,000288507 | 0,02259107 | ENSG00000138642 | HERC6 |
| 544,8895279 | -1,000262683 | 0,286132994 | -3,495796376 | 0,000472649 | 0,033594136 | ENSG00000148400 | NOTCH1 |
| 230,1073074 | 1,00540825 | 0,277995219 | 3,616638639 | 0,000298453 | 0,02307286 | ENSG00000066827 | ZFAT |
| 255,7427818 | 1,005533963 | 0,263819084 | 3,811452709 | 0,000138152 | 0,012840703 | ENSG00000280721 | LINC01943 |
| 272,7640764 | 1,008238031 | 0,29739911 | 3,390185103 | 0,000698454 | 0,044591738 | ENSG00000204257 | HLA-DMA |
| 326,9530545 | 1,008590717 | 0,248989166 | 4,050741375 | 5,10556E-05 | 0,005864924 | ENSG00000172893 | DHCR7 |
| 324,4653391 | 1,0127313 | 0,245118682 | 4,131595737 | 3,60254E-05 | 0,004489892 | ENSG00000081913 | PHLPP1 |
| 112,1025985 | 1,014160385 | 0,302378181 | 3,353946971 | 0,000796676 | 0,048854826 | ENSG00000132825 | PPP1R3D |
| 158,6113131 | 1,02580175 | 0,303142184 | 3,383896413 | 0,00071465 | 0,045096187 | ENSG00000115325 | DOK1 |
| 304,5341778 | 1,036651809 | 0,248384391 | 4,173578719 | 2,99852E-05 | 0,003928663 | ENSG00000114023 | FAM162A |
| 693,747533 | 1,054697092 | 0,29324616 | 3,596627126 | 0,00032237 | 0,024401395 | ENSG00000179029 | TMEM107 |
| 491,9344261 | 1,063407413 | 0,250870643 | 4,238867483 | 2,2465E-05 | 0,003118719 | ENSG00000287979 | ENSG00000287979 |
| 366,4634629 | 1,070328405 | 0,256045023 | 4,180235154 | 2,91208E-05 | 0,003830792 | ENSG00000285646 | ENSG00000285646 |
| 466,1263308 | 1,071278911 | 0,228490493 | 4,688505408 | 2,75208E-06 | 0,000594617 | ENSG00000124610 | H1-1 |
| 150,167729 | 1,075206859 | 0,28118803 | 3,823800252 | 0,00013141 | 0,012462595 | ENSG00000143653 | SCCPDH |
| 128,8289446 | 1,090674374 | 0,320201678 | 3,406210673 | 0,000658713 | 0,042893936 | ENSG00000134508 | CABLES1 |
| 423,9274163 | 1,107359407 | 0,221957506 | 4,98906041 | 6,06737E-07 | 0,000178326 | ENSG00000128578 | STRIP2 |
| 468,8932589 | 1,129394196 | 0,273417189 | 4,130662739 | 3,61719E-05 | 0,004489892 | ENSG00000162433 | AK4 |
| 192,825432 | 1,129928389 | 0,3334814 | 3,388280095 | 0,000703324 | 0,04464055 | ENSG00000011590 | ZBTB32 |
| 232,0112192 | 1,132066427 | 0,251523852 | 4,500831304 | 6,76882E-06 | 0,001224563 | ENSG00000125726 | CD70 |
| 220,8552741 | 1,140288171 | 0,320275998 | 3,560329774 | 0,000370389 | 0,027400413 | ENSG00000067208 | EVI5 |
| 537,7861349 | 1,142393028 | 0,249608498 | 4,576739329 | 4,72279E-06 | 0,000951089 | ENSG00000060558 | GNA15 |
| 311,404889 | 1,145540402 | 0,249004486 | 4,600480974 | 4,21517E-06 | 0,000864878 | ENSG00000279602 | ENSG00000279602 |
| 556,9783651 | 1,181820084 | 0,316513146 | 3,733873608 | 0,000188557 | 0,016625644 | ENSG00000207166 | SNORA68 |
| 44225,88388 | 1,190218661 | 0,235597232 | 5,051921235 | 4,37388E-07 | 0,000134616 | ENSG00000265185 | ENSG00000265185 |
| 186,9887959 | 1,192044151 | 0,268946382 | 4,432274356 | 9,32443E-06 | 0,001622877 | ENSG00000141698 | NT5C3B |
| 177,6354658 | 1,201341619 | 0,278725869 | 4,310118832 | 1,63167E-05 | 0,002430664 | ENSG00000161956 | SENP3 |
| 641,7681458 | 1,204601681 | 0,231804199 | 5,196634439 | 2,02929E-07 | 6,62035E-05 | ENSG00000198814 | GK |
| 92,0503494 | 1,21970929 | 0,351510033 | 3,469913158 | 0,000520627 | 0,035833174 | ENSG00000120280 | TASL |
| 392,5587359 | 1,219956303 | 0,284346499 | 4,290386234 | 1,78363E-05 | 0,002613755 | ENSG00000121039 | RDH10 |
| 1495,808756 | 1,222540028 | 0,212946946 | 5,741054537 | 9,40888E-09 | 4,58142E-06 | ENSG00000177606 | JUN |
| 94,35949304 | 1,223993979 | 0,340994143 | 3,589486811 | 0,00033133 | 0,024829874 | ENSG00000243244 | STON1 |
| 263,5043048 | 1,23341168 | 0,272754173 | 4,52206347 | 6,12397E-06 | 0,001166507 | ENSG00000104341 | LAPTM4B |
| 528,1192227 | 1,238202913 | 0,243658128 | 5,081722184 | 3,74028E-07 | 0,00011733 | ENSG00000141655 | TNFRSF11A |
| 276,2400438 | 1,285110621 | 0,375253481 | 3,42464677 | 0,000615599 | 0,040654472 | ENSG00000141526 | SLC16A3 |
| 1060,415925 | 1,288577937 | 0,196826368 | 6,54677496 | 5,87929E-11 | 4,82999E-08 | ENSG00000179583 | CIITA |
| 751,7182729 | 1,294351711 | 0,228628401 | 5,661377599 | 1,50163E-08 | 6,48594E-06 | ENSG00000198502 | HLA-DRB5 |
| 192,7059183 | 1,300138973 | 0,276513742 | 4,701896418 | 2,57756E-06 | 0,000568122 | ENSG00000118777 | ABCG2 |
| 221,6657163 | 1,311158694 | 0,241686546 | 5,425037983 | 5,79422E-08 | 2,10034E-05 | ENSG00000242574 | HLA-DMB |
| 179,4070228 | 1,312468179 | 0,320452158 | 4,095675894 | 4,20938E-05 | 0,005067416 | ENSG00000114013 | CD86 |
| 160,0464324 | 1,314858835 | 0,369683906 | 3,556711057 | 0,000375527 | 0,027717616 | ENSG00000196576 | PLXNB2 |
| 177,2779069 | 1,325949304 | 0,275623844 | 4,8107206 | 1,50387E-06 | 0,000377402 | ENSG00000183780 | SLC35F3 |
| 117,1885835 | 1,32667435 | 0,326959446 | 4,057611317 | 4,95772E-05 | 0,005755894 | ENSG00000204475 | NCR3 |
| 425,3526258 | 1,336797749 | 0,218055547 | 6,130537686 | 8,75826E-10 | 5,83121E-07 | ENSG00000151914 | DST |
| 109,4072141 | 1,338909619 | 0,360617057 | 3,712829417 | 0,000204955 | 0,01778312 | ENSG00000184792 | OSBP2 |
| 98,02173172 | 1,340251168 | 0,346683702 | 3,865919165 | 0,000110672 | 0,01081003 | ENSG00000240350 | ENSG00000240350 |
| 246,4115531 | 1,348320714 | 0,308173142 | 4,375205133 | 1,21318E-05 | 0,001930678 | ENSG00000207205 | RNVU1-15 |
| 129692,7217 | 1,355266096 | 0,218709398 | 6,1966523 | 5,76767E-10 | 4,0035E-07 | ENSG00000264940 | ENSG00000264940 |
| 109,9932 | 1,355588685 | 0,326220703 | 4,155434252 | 3,2467E-05 | 0,004137517 | ENSG00000227630 | LINC01132 |
| 92,22433865 | 1,377023819 | 0,353308577 | 3,897510306 | 9,71867E-05 | 0,009696081 | ENSG00000135750 | KCNK1 |
| 186,767949 | 1,37762308 | 0,37863757 | 3,638368695 | 0,00027437 | 0,021725878 | ENSG00000222724 | RNU2-63P |
| 122,4392336 | 1,39049847 | 0,313766332 | 4,431636951 | 9,35204E-06 | 0,001622877 | ENSG00000129116 | PALLD |
| 1951,936095 | 1,414072388 | 0,170767157 | 8,280704611 | 1,22449E-16 | 3,32897E-13 | ENSG00000204287 | HLA-DRA |
| 930,9010894 | 1,430077876 | 0,252649842 | 5,660315735 | 1,51095E-08 | 6,48594E-06 | ENSG00000262074 | ENSG00000262074 |
| 184,8366998 | 1,438483987 | 0,325521138 | 4,419018666 | 9,91501E-06 | 0,001693546 | ENSG00000214338 | MTCL3 |
| 930,0545913 | 1,455112807 | 0,209358551 | 6,950338563 | 3,64411E-12 | 4,09949E-09 | ENSG00000223865 | HLA-DPB1 |
| 119,5562196 | 1,463880981 | 0,335318288 | 4,365646114 | 1,26748E-05 | 0,002007287 | ENSG00000147852 | VLDLR |
| 62,08323664 | 1,480292791 | 0,402521401 | 3,67755053 | 0,000235484 | 0,01949859 | ENSG00000148225 | WDR31 |
| 133,6418289 | 1,483215753 | 0,311398009 | 4,763086822 | 1,90654E-06 | 0,000444278 | ENSG00000050438 | SLC4A8 |
| 229,3468894 | 1,483519559 | 0,336681571 | 4,406298673 | 1,05152E-05 | 0,001768287 | ENSG00000210194 | ENSG00000210194 |
| 92,7046388 | 1,522979662 | 0,430723858 | 3,535860929 | 0,000406449 | 0,02946662 | ENSG00000200972 | RNU5A-8P |
| 152,0855745 | 1,531283701 | 0,339352871 | 4,512364071 | 6,4109E-06 | 0,001188348 | ENSG00000177494 | ZBED2 |
| 175,9733793 | 1,535148558 | 0,36366605 | 4,221313913 | 2,42882E-05 | 0,003301582 | ENSG00000273338 | ENSG00000273338 |
| 68,40629946 | 1,551924095 | 0,40888397 | 3,795512197 | 0,000147339 | 0,013426776 | ENSG00000146416 | AIG1 |
| 220,7307193 | 1,561010705 | 0,312679021 | 4,992374285 | 5,96415E-07 | 0,000176886 | ENSG00000212195 | LOC124904154 |
| 53,45459676 | 1,56301097 | 0,465970037 | 3,354316473 | 0,000795613 | 0,048854826 | ENSG00000135643 | KCNMB4 |
| 234,909838 | 1,578082934 | 0,300208914 | 5,256615837 | 1,4673E-07 | 4,98639E-05 | ENSG00000277925 | TERC |
| 137,4657702 | 1,579914734 | 0,360862643 | 4,378160952 | 1,19685E-05 | 0,001921085 | ENSG00000286172 | RNVU1-8 |
| 52,7909636 | 1,580879854 | 0,463294824 | 3,412254514 | 0,000644279 | 0,042122179 | ENSG00000103202 | NME4 |
| 62,97784639 | 1,5892044 | 0,468287936 | 3,393647961 | 0,000689683 | 0,044291761 | ENSG00000260035 | ENSG00000260035 |
| 1556,430452 | 1,597210765 | 0,220862147 | 7,231708955 | 4,76954E-13 | 6,76527E-10 | ENSG00000196126 | HLA-DRB1 |

| | | | | | | | |
|---|---|---|---|---|---|---|---|
| 145,8348048 | 1,598295138 | 0,318986096 | 5,010547979 | 5,42753E-07 | 0,000163952 | ENSG00000229391 | ENSG00000229391 |
| 1406,353855 | 1,606249374 | 0,219887781 | 7,304859623 | 2,77556E-13 | 4,11591E-10 | ENSG00000179344 | HLA-DQB1 |
| 172,8644777 | 1,609502198 | 0,270583788 | 5,948258051 | 2,71011E-09 | 1,55113E-06 | ENSG00000005102 | MEOX1 |
| 1009,016618 | 1,627533595 | 0,289422469 | 5,623383705 | 1,87253E-08 | 7,72024E-06 | ENSG00000072110 | ACTN1 |
| 47,5448985 | 1,654027995 | 0,480705219 | 3,440836355 | 0,000579919 | 0,038532143 | ENSG00000154319 | FAM167A |
| 61,07691035 | 1,655462338 | 0,47720298 | 3,469094719 | 0,000522215 | 0,035866847 | ENSG00000135919 | SERPINE2 |
| 262,9659011 | 1,669698846 | 0,28520626 | 5,854355528 | 4,78863E-09 | 2,56105E-06 | ENSG00000270022 | ENSG00000270022 |
| 56,11343782 | 1,679430176 | 0,465450747 | 3,608180215 | 0,000308352 | 0,023535825 | ENSG00000171757 | LRRC34 |
| 176,2727742 | 1,696246896 | 0,358607044 | 4,730099216 | 2,2441E-06 | 0,000501449 | ENSG00000020577 | SAMD4A |
| 69,58648378 | 1,698238052 | 0,410400562 | 4,138001289 | 3,50344E-05 | 0,004412987 | ENSG00000214193 | SH3D21 |
| 138,8417586 | 1,702275747 | 0,395583166 | 4,303205725 | 1,68344E-05 | 0,002496394 | ENSG00000171169 | NAIF1 |
| 96,2689003 | 1,737689829 | 0,398623998 | 4,359220317 | 1,30527E-05 | 0,00204726 | ENSG00000278774 | LOC124904146 |
| 51,91875581 | 1,739429134 | 0,499605707 | 3,481603812 | 0,000498421 | 0,03504412 | ENSG00000275291 | RNVU1-26 |
| 137,4151843 | 1,744590954 | 0,288105623 | 6,055386689 | 1,40081E-09 | 8,46295E-07 | ENSG00000165474 | GJB2 |
| 36,16063462 | 1,760578125 | 0,494309925 | 3,561688806 | 0,000368477 | 0,027320893 | ENSG00000177181 | RIMKLA |
| 51,3767284 | 1,79472854 | 0,461605805 | 3,888011202 | 0,000101069 | 0,009991739 | ENSG00000010278 | CD9 |
| 65,44668777 | 1,826270007 | 0,491200324 | 3,71797395 | 0,000200827 | 0,01754249 | ENSG00000155980 | KIF5A |
| 196,8321352 | 1,83885754 | 0,333217221 | 5,518494908 | 3,41915E-08 | 1,29705E-05 | ENSG00000188761 | BCL2L15 |
| 51,83931417 | 1,853958269 | 0,458256025 | 4,045682251 | 5,2171E-05 | 0,005930402 | ENSG00000150938 | CRIM1 |
| 48,573972 | 1,858086199 | 0,458672693 | 4,051006801 | 5,09977E-05 | 0,005864924 | ENSG00000272768 | ENSG00000272768 |
| 2397,690729 | 1,880084528 | 0,182136174 | 10,32241147 | 5,58037E-25 | 3,64108E-21 | ENSG00000231389 | HLA-DPA1 |
| 41,4173843 | 1,881953196 | 0,531653523 | 3,539811392 | 0,000400413 | 0,029093711 | ENSG00000198910 | L1CAM |
| 119,4982723 | 1,886769619 | 0,419227824 | 4,500583001 | 6,77673E-06 | 0,001224563 | ENSG00000152207 | CYSLTR2 |
| 91,7117972 | 1,895815818 | 0,358574374 | 5,287092323 | 1,24276E-07 | 4,31317E-05 | ENSG00000105383 | CD33 |
| 170,6589022 | 1,912409548 | 0,314196114 | 6,086674732 | 1,1528E-09 | 7,23248E-07 | ENSG00000204252 | HLA-DOA |
| 68,49039728 | 1,917099336 | 0,420625076 | 4,557739053 | 5,17072E-06 | 0,001016203 | ENSG00000133454 | MYO18B |
| 30,47387737 | 1,936912355 | 0,556799616 | 3,478652465 | 0,000503942 | 0,035204696 | ENSG00000140465 | CYP1A1 |
| 95,30937685 | 1,938232116 | 0,341946138 | 5,668238067 | 1,44273E-08 | 6,44764E-06 | ENSG00000119411 | BSPRY |
| 50,71050431 | 1,94336468 | 0,467317488 | 4,158553297 | 3,2027E-05 | 0,00409744 | ENSG00000071909 | MYO3B |
| 71,5275645 | 2,012076774 | 0,480875335 | 4,184196252 | 2,86177E-05 | 0,003779851 | ENSG00000099250 | NRP1 |
| 55,48968186 | 2,018410464 | 0,458647268 | 4,400790329 | 1,07857E-05 | 0,001795274 | ENSG00000179588 | ZFPM1 |
| 42,90154225 | 2,034298365 | 0,51389374 | 3,958597286 | 7,53912E-05 | 0,007808329 | ENSG00000224557 | HLA-DPB2 |
| 27,96615045 | 2,061697662 | 0,607440288 | 3,394074617 | 0,000688609 | 0,044291761 | ENSG00000128683 | GAD1 |
| 160,3157824 | 2,077929155 | 0,297877985 | 6,975772833 | 3,04194E-12 | 3,67556E-09 | ENSG00000133812 | SBF2 |
| 98,77351014 | 2,079644821 | 0,443563175 | 4,688497464 | 2,75218E-06 | 0,000594617 | ENSG00000144645 | OSBPL10 |
| 874,1752952 | 2,142201761 | 0,185017462 | 11,57837609 | 5,30413E-31 | 8,6521E-27 | ENSG00000196735 | HLA-DQA1 |
| 31,7897073 | 2,153410229 | 0,605832809 | 3,554462875 | 0,000378752 | 0,027892561 | ENSG00000108176 | DNAJC12 |
| 55,66479014 | 2,157084311 | 0,454199111 | 4,749204172 | 2,04219E-06 | 0,000470416 | ENSG00000073282 | TP63 |
| 100,1615509 | 2,187821017 | 0,369230417 | 5,925354235 | 3,11625E-09 | 1,75284E-06 | ENSG00000107242 | PIP5K1B |
| 174,8779079 | 2,253630349 | 0,42998513 | 5,241182054 | 1,59551E-07 | 5,36618E-05 | ENSG00000189221 | MAOA |
| 22,30602863 | 2,26900309 | 0,656446682 | 3,456492588 | 0,000547254 | 0,037117719 | ENSG00000100867 | DHRS2 |
| 62,34688647 | 2,321903188 | 0,654641863 | 3,546829678 | 0,000389897 | 0,028456344 | ENSG00000258311 | ENSG00000258311 |
| 22,7524074 | 2,396506705 | 0,715155986 | 3,351026564 | 0,000805126 | 0,048913266 | ENSG00000111249 | CUX2 |
| 29,94066439 | 2,441128083 | 0,679856483 | 3,590652061 | 0,000329852 | 0,024795121 | ENSG00000165025 | SYK |
| 36,70107491 | 2,44355998 | 0,531970701 | 4,593410835 | 4,36059E-06 | 0,000889125 | ENSG00000122861 | PLAU |
| 23,23340603 | 2,475657634 | 0,660431838 | 3,748543744 | 0,000177864 | 0,015941334 | ENSG00000215284 | ENSG00000215284 |
| 37,10846454 | 2,508250678 | 0,656401266 | 3,821215479 | 0,000132796 | 0,012490965 | ENSG00000176014 | TUBB6 |
| 57,62576523 | 2,509699962 | 0,556729745 | 4,507932229 | 6,54625E-06 | 0,001206581 | ENSG00000129226 | CD68 |
| 149,8820457 | 2,521828898 | 0,327573317 | 7,698517452 | 1,37654E-14 | 3,20773E-11 | ENSG00000132170 | PPARG |
| 54,04042451 | 2,548568403 | 0,5164563 | 4,934722267 | 8,02649E-07 | 0,000230847 | ENSG00000141574 | SECTM1 |
| 142,9473423 | 2,576556789 | 0,42134218 | 6,115117146 | 9,64861E-10 | 6,17208E-07 | ENSG00000166025 | AMOTL1 |
| 27,18550658 | 2,591472065 | 0,637457679 | 4,065324099 | 4,79658E-05 | 0,005640046 | ENSG00000162373 | BEND5 |
| 38,98177744 | 2,662886218 | 0,736118965 | 3,617467211 | 0,0002975 | 0,023053777 | ENSG00000255526 | NEDD8-MDP1 |
| 21,81744014 | 2,732000349 | 0,71524291 | 3,819681835 | 0,000133624 | 0,012490965 | ENSG00000143079 | CTTNBP2NL |
| 40,81327389 | 2,818678045 | 0,519111333 | 5,429814124 | 5,64128E-08 | 2,08992E-05 | ENSG00000154734 | ADAMTS1 |
| 24,28376913 | 2,88550598 | 0,772463851 | 3,735457621 | 0,000187374 | 0,016582622 | ENSG00000120658 | ENOX1 |
| 38,25071097 | 2,885992492 | 0,774994115 | 3,723889556 | 0,000196177 | 0,017250865 | ENSG00000172116 | CD8B |
| 25,10266262 | 2,908097991 | 0,827093525 | 3,516044925 | 0,000438027 | 0,031407007 | ENSG00000136237 | RAPGEF5 |
| 29,09502494 | 2,915538499 | 0,820971997 | 3,55132515 | 0,000383297 | 0,028038251 | ENSG00000120549 | KIAA1217 |
| 16,89453334 | 2,933325548 | 0,76120932 | 3,853507135 | 0,000116438 | 0,011238666 | ENSG00000106066 | CPVL |
| 190,6644432 | 2,934941056 | 0,313881455 | 9,350476156 | 8,7254E-21 | 3,55822E-17 | ENSG00000140479 | PCSK6 |
| 20,53844781 | 2,957363382 | 0,808069485 | 3,659788468 | 0,000252424 | 0,020434406 | ENSG00000087303 | NID2 |
| 54,2359054 | 2,963083168 | 0,554718483 | 5,341598051 | 9,21308E-08 | 3,26704E-05 | ENSG00000144642 | RBMS3 |
| 74,49968827 | 2,98179309 | 0,555691585 | 5,365913705 | 8,05404E-08 | 2,88742E-05 | ENSG00000109265 | CRACD |
| 32,5062841 | 3,112026731 | 0,548961976 | 5,668929479 | 1,43692E-08 | 6,44764E-06 | ENSG00000170365 | SMAD1 |
| 82,81672478 | 3,121684697 | 0,416428328 | 7,496331267 | 6,56288E-14 | 1,25946E-10 | ENSG00000115414 | FN1 |
| 22,44370436 | 3,131578713 | 0,703302645 | 4,452675865 | 8,48067E-06 | 0,001487514 | ENSG00000128322 | IGLL1 |
| 19,77976905 | 3,18568103 | 0,794542721 | 4,009452163 | 6,08598E-05 | 0,006618298 | ENSG00000187079 | TEAD1 |
| 20,46169093 | 3,203633369 | 0,824962812 | 3,883367008 | 0,00010302 | 0,01015384 | ENSG00000141404 | GNAL |
| 16,65385637 | 3,305182461 | 0,908830007 | 3,636744426 | 0,000276106 | 0,02181034 | ENSG00000153165 | RGPD3 |
| 22,18372144 | 3,40896467 | 0,849620251 | 4,012339237 | 6,012E-05 | 0,006559718 | ENSG00000147642 | SYBU |
| 12,74192775 | 3,441426344 | 0,999295169 | 3,443853679 | 0,000573486 | 0,038260545 | ENSG00000279970 | ENSG00000279970 |
| 48,74170765 | 3,494455609 | 0,547988853 | 6,376873532 | 1,8074E-10 | 1,40392E-07 | ENSG00000115008 | IL1A |
| 19,35023167 | 3,733162197 | 0,844669473 | 4,419672212 | 9,88507E-06 | 0,001693546 | ENSG00000152463 | OLAH |
| 91,23578866 | 4,082042856 | 0,499506393 | 8,172153372 | 3,02933E-16 | 7,60222E-13 | ENSG00000136099 | PCDH8 |

| | | | | | | | |
|---|---|---|---|---|---|---|---|
| 19,25959287 | 4,090468573 | 0,856406335 | 4,776317507 | 1,78534E-06 | 0,000424105 | ENSG00000196951 | SCOC-AS1 |
| 49,36767519 | 4,26726077 | 0,619653036 | 6,886532498 | 5,71687E-12 | 6,01636E-09 | ENSG00000158125 | XDH |
| 14,8230985 | 4,315803939 | 1,053972851 | 4,094796117 | 4,2254E-05 | 0,005067995 | ENSG00000042832 | TG |
| 12,6952079 | 4,3473591 | 1,196288005 | 3,634040533 | 0,000279017 | 0,021987079 | ENSG00000286132 | ENSG00000286132 |
| 153,2402767 | 4,353579943 | 1,106917174 | 3,933067484 | 8,38687E-05 | 0,008577216 | ENSG00000113721 | PDGFRB |
| 13,02248381 | 4,422029072 | 1,060536723 | 4,169614286 | 3,05116E-05 | 0,003981636 | ENSG00000154096 | THY1 |
| 19,48333292 | 4,456567422 | 1,059299504 | 4,207089125 | 2,58681E-05 | 0,003444577 | ENSG00000187045 | TMPRSS6 |
| 165,6595613 | 4,535385244 | 0,42384555 | 10,70056119 | 1,01164E-26 | 8,25091E-23 | ENSG00000164303 | ENPP6 |
| 331,5243535 | 4,564052757 | 0,664699629 | 6,866338654 | 6,58706E-12 | 6,71551E-09 | ENSG00000169429 | CXCL8 |
| 19015,80497 | 5,009496209 | 0,17719867 | 28,27050682 | 7,9673E-176 | 2,5992E-171 | ENSG00000173757 | STAT5B |
| 29,68023426 | 5,034582359 | 0,82297253 | 6,11755821 | 9,50201E-10 | 6,17208E-07 | ENSG00000157110 | RBPMS |
| 118,1019313 | 5,07589854 | 0,46139898 | 11,00110482 | 3,77478E-28 | 4,10495E-24 | ENSG00000151474 | FRMD4A |
| 10,04937143 | 5,163037264 | 1,356485728 | 3,806186205 | 0,000141126 | 0,013005931 | ENSG00000074047 | GLI2 |
| 7,247876169 | 5,169172304 | 1,489034826 | 3,471491877 | 0,000517575 | 0,035698449 | ENSG00000116990 | MYCL |
| 6,963782903 | 5,430435229 | 1,482184852 | 3,663804297 | 0,000248497 | 0,020369235 | ENSG00000144407 | PTH2R |
| 30,69738857 | 5,500652341 | 0,94757728 | 5,804964362 | 6,43797E-09 | 3,38762E-06 | ENSG00000254126 | CD8B2 |
| 6,389667762 | 5,555703943 | 1,642505963 | 3,382455874 | 0,000718408 | 0,045245847 | ENSG00000091428 | RAPGEF4 |
| 9,672714539 | 5,607165731 | 1,599084535 | 3,506484871 | 0,000454067 | 0,032415371 | ENSG00000283201 | ZNF724 |
| 20,91651503 | 5,665711662 | 1,022450022 | 5,54130915 | 3,00219E-08 | 1,18004E-05 | ENSG00000277481 | PKD1L3 |
| 7,861544642 | 5,92370315 | 1,571478957 | 3,769508414 | 0,000163569 | 0,014781961 | ENSG00000109625 | CPZ |
| 88,12071373 | 5,935848584 | 0,623066375 | 9,526831848 | 1,62156E-21 | 7,55741E-18 | ENSG00000152377 | SPOCK1 |
| 39,88575751 | 8,544168361 | 1,291091353 | 6,61778761 | 3,64614E-11 | 3,21491E-08 | ENSG00000274049 | INO80B-WBP1 |
| 0,630221196 | 14,07076406 | 4,05003825 | 3,47422992 | 0,000512322 | 0,035440393 | ENSG00000171759 | PAH |
| 0,630221196 | 14,07076406 | 4,05003825 | 3,47422992 | 0,000512322 | 0,035440393 | ENSG00000233741 | RPL27P7 |
| 0,676189846 | 14,40680667 | 4,014755342 | 3,588464412 | 0,000332631 | 0,024832417 | ENSG00000142871 | CCN1 |
| 0,613172142 | 14,45598968 | 4,016313042 | 3,599318461 | 0,000319052 | 0,024206422 | ENSG00000115598 | IL1RL2 |
| 0,613172142 | 14,45598968 | 4,016313042 | 3,599318461 | 0,000319052 | 0,024206422 | ENSG00000250306 | ENSG00000250306 |
| 0,812656632 | 14,63889284 | 4,012710756 | 3,648130584 | 0,000264155 | 0,021070428 | ENSG00000182489 | XKRX |
| 0,808159552 | 14,76563044 | 4,008139268 | 3,683911525 | 0,000229682 | 0,019213193 | ENSG00000259992 | ENSG00000259992 |
| 0,808159552 | 14,76563044 | 4,008139268 | 3,683911525 | 0,000229682 | 0,019213193 | ENSG00000271286 | CYTH1P1 |
| 1,14671293 | 14,76887014 | 3,44125596 | 4,29170928 | 1,77303E-05 | 0,002613755 | ENSG00000270038 | ENSG00000270038 |
| 0,725577138 | 14,77913104 | 4,001421453 | 3,693470236 | 0,000221214 | 0,018745192 | ENSG00000242255 | ENSG00000242255 |
| 0,725577138 | 14,77913104 | 4,001421453 | 3,693470236 | 0,000221214 | 0,018745192 | ENSG00000257335 | MGAM |
| 1,148568997 | 14,83581483 | 3,729451055 | 3,978015696 | 6,94928E-05 | 0,007354531 | ENSG00000205704 | SMIM45 |
| 0,925061628 | 14,84287422 | 3,998772495 | 3,711857636 | 0,000205744 | 0,017804198 | ENSG00000269524 | ENSG00000269524 |
| 0,914396184 | 14,84537455 | 4,006676595 | 3,705159176 | 0,000211258 | 0,018207834 | ENSG00000288704 | ENSG00000288704 |
| 1,31652518 | 14,95374978 | 3,54998439 | 4,212342404 | 2,52736E-05 | 0,003379205 | ENSG00000243721 | ENSG00000243721 |
| 6,528662471 | 14,98985627 | 3,980701974 | 3,765631381 | 0,000166129 | 0,014971779 | ENSG00000281039 | ENSG00000281039 |
| 0,837982134 | 15,08365384 | 3,993683073 | 3,776878025 | 0,000158806 | 0,014431481 | ENSG00000201498 | ENSG00000201498 |
| 0,967436185 | 15,18177458 | 3,974332761 | 3,819955572 | 0,000133476 | 0,012490965 | ENSG00000261239 | ENSG00000261239 |
| 0,919097516 | 15,18896193 | 3,795438819 | 4,001898767 | 6,28362E-05 | 0,006810524 | ENSG00000239699 | HNRNPA3P8 |
| 0,82093308 | 15,20306778 | 3,992179816 | 3,808212175 | 0,000139975 | 0,0129364 | ENSG00000284250 | MIR6516 |
| 0,87266268 | 15,20769663 | 3,990897328 | 3,810595807 | 0,000138632 | 0,012848691 | ENSG00000243136 | RN7SL22P |
| 1,596886359 | 15,28181281 | 3,312451444 | 4,613445077 | 3,96049E-06 | 0,000822975 | ENSG00000261771 | DNAAF4-CCPG1 |
| 0,813959254 | 15,28836404 | 3,9931321 | 3,828664731 | 0,00012884 | 0,012326359 | ENSG00000225563 | ENSG00000225563 |
| 0,946818191 | 15,29439887 | 3,990045827 | 3,833138648 | 0,000126519 | 0,012139831 | ENSG00000166763 | ENSG00000166763 |
| 0,794792592 | 15,32583053 | 4,009152923 | 3,82271039 | 0,000131993 | 0,012481543 | ENSG00000148926 | ADM |
| 0,674372688 | 15,4647101 | 4,001198854 | 3,865019125 | 0,00011108 | 0,010817582 | ENSG00000269481 | ENSG00000269481 |
| 0,838399001 | 15,54648115 | 3,99632188 | 3,89019744 | 0,000100163 | 0,009945867 | ENSG00000283886 | ENSG00000283886 |
| 0,75442091 | 15,55661205 | 3,999268069 | 3,88986479 | 0,0001003 | 0,009945867 | ENSG00000231362 | ENSG00000231362 |
| 1,140604926 | 15,55788647 | 3,531926579 | 4,404929187 | 1,05818E-05 | 0,00177037 | ENSG00000224814 | ENSG00000224814 |
| 1,094320433 | 15,59096082 | 3,786596329 | 4,117407685 | 3,83158E-05 | 0,004734903 | ENSG00000180834 | MAP6D1 |
| 1,510082472 | 15,59544874 | 3,399369662 | 4,587747227 | 4,48055E-06 | 0,000907909 | ENSG00000204196 | ENSG00000204196 |
| 1,038769246 | 15,59987597 | 3,85025898 | 4,051643293 | 5,08592E-05 | 0,005864924 | ENSG00000187726 | DNAJB13 |
| 1,091824877 | 15,60298457 | 3,77254741 | 4,135928028 | 3,53523E-05 | 0,004435898 | ENSG00000230953 | ENSG00000230953 |
| 0,933280788 | 15,62961893 | 3,995520615 | 3,911785331 | 9,16163E-05 | 0,009224973 | ENSG00000285068 | ENSG00000285068 |
| 0,922377093 | 15,64219414 | 3,994443881 | 3,91598796 | 9,00347E-05 | 0,009122029 | ENSG00000241002 | RPL32P13 |
| 1,206725429 | 15,76096853 | 3,650325818 | 4,317688151 | 1,57672E-05 | 0,002370457 | ENSG00000262117 | BCAR4 |
| 0,921686137 | 15,91323265 | 3,994191011 | 3,984094052 | 6,7738E-05 | 0,007221849 | ENSG00000200741 | ENSG00000200741 |
| 0,921686137 | 15,91323265 | 3,994191011 | 3,984094052 | 6,7738E-05 | 0,007221849 | ENSG00000285629 | ENSG00000285629 |
| 1,34755733 | 15,93509041 | 3,499660939 | 4,553324074 | 5,28048E-06 | 0,00102542 | ENSG00000288000 | ENSG00000288000 |
| 1,34755733 | 15,93509041 | 3,499660939 | 4,553324074 | 5,28048E-06 | 0,00102542 | ENSG00000288635 | ENSG00000288635 |
| 1,593510868 | 15,93565074 | 3,309416512 | 4,815244826 | 1,4702E-06 | 0,000371811 | ENSG00000170788 | DYDC1 |
| 2,348622734 | 16,14637285 | 2,916061722 | 5,5370477 | 3,07613E-08 | 1,19471E-05 | ENSG00000254633 | ENSG00000254633 |
| 1,449832882 | 16,15795177 | 3,261917551 | 4,953513236 | 7,28854E-07 | 0,000212305 | ENSG00000272482 | ENSG00000272482 |
| 0,966203218 | 16,17620401 | 3,954219657 | 4,090871376 | 4,29755E-05 | 0,005116913 | ENSG00000254641 | ENSG00000254641 |
| 1,823987246 | 16,43778427 | 3,02221121 | 5,438992555 | 5,35827E-08 | 2,00929E-05 | ENSG00000130751 | NPAS1 |
| 2,467699422 | 16,64841393 | 2,880541993 | 5,779611605 | 7,48733E-09 | 3,87725E-06 | ENSG00000277599 | ENSG00000277599 |
| 2,299247083 | 16,6746696 | 2,914251565 | 5,721767401 | 1,05422E-08 | 5,05775E-06 | ENSG00000272104 | ENSG00000272104 |
| 1,25621659 | 16,78036425 | 3,851747487 | 4,356558759 | 1,32123E-05 | 0,002052566 | ENSG00000282780 | TRBJ1-6 |
| 3,620947309 | 17,22455994 | 2,576948003 | 6,684092936 | 2,32359E-11 | 2,10569E-08 | ENSG00000260371 | ENSG00000260371 |
| 2,62746998 | 17,23950351 | 2,849226165 | 6,050591463 | 1,44315E-09 | 8,56024E-07 | ENSG00000273167 | ENSG00000273167 |
| 5,390229318 | 17,92378205 | 3,965995741 | 4,519364926 | 6,20254E-06 | 0,001169663 | ENSG00000288534 | TMX2-CTNND1 |
| 1,739020882 | 18,09538365 | 3,218964143 | 5,621492768 | 1,89314E-08 | 7,72024E-06 | ENSG00000285547 | ENSG00000285547 |
| 0,898163443 | 18,23420304 | 3,761376617 | 4,847747222 | 1,24871E-06 | 0,000328533 | ENSG00000254012 | RPL10P8 |

| | | | | | | | |
|---|---|---|---|---|---|---|---|
| 6,125240702 | 18,43304913 | 3,965558421 | 4,648285859 | 3,34705E-06 | 0,000705391 | ENSG00000228589 | ENSG00000228589 |
| 4,093012685 | 19,0591378 | 3,969282659 | 4,801657992 | 1,57357E-06 | 0,000384517 | ENSG00000284976 | LOC122513141 |
| 6,352708034 | 23,02758232 | 3,920661032 | 5,873392811 | 4,26965E-09 | 2,3609E-06 | ENSG00000144785 | ENSG00000144785 |

E10 BACH2.SAM_vs_non.act

| ensembl_gene_id | baseMean | log2FoldChange | lfcSE | stat | pvalue | padj | hgnc_symbol | gene_biotype |
|---|---|---|---|---|---|---|---|---|
| ENSG00000003400 | 38,17543049 | 2,202415806 | 0,379333404 | 5,806015984 | 6,39769E-09 | 1,60274E-07 | CASP10 | protein_coding |
| ENSG00000005884 | 600,0823205 | 2,256944337 | 0,103228629 | 21,86355044 | 5,7771E-106 | 4,0726E-102 | ITGA3 | protein_coding |
| ENSG00000006283 | 99,62152007 | 2,136434505 | 0,267058625 | 7,999870843 | 1,2455E-15 | 8,85107E-14 | CACNA1G | protein_coding |
| ENSG00000006611 | 112,5025588 | 3,955490463 | 0,34007289 | 11,63130194 | 2,85706E-31 | 9,32461E-29 | USH1C | protein_coding |
| ENSG00000007516 | 219,3636063 | 2,291245152 | 0,179551037 | 12,76096862 | 2,70812E-37 | 1,59824E-34 | BAIAP3 | protein_coding |
| ENSG00000009709 | 16,26917718 | 6,010632709 | 1,068387076 | 5,625894251 | 1,84549E-08 | 4,12196E-07 | PAX7 | protein_coding |
| ENSG00000010310 | 139,6725342 | 2,06053162 | 0,202525395 | 10,17418889 | 2,58545E-24 | 4,57949E-22 | GIPR | protein_coding |
| ENSG00000013016 | 46,74533462 | 2,763343201 | 0,493459387 | 5,599940484 | 2,14425E-08 | 4,70322E-07 | EHD3 | protein_coding |
| ENSG00000015520 | 21,63789286 | 4,003009538 | 0,591879133 | 6,763221266 | 1,34957E-11 | 5,69014E-10 | NPC1L1 | protein_coding |
| ENSG00000019505 | 29,27880986 | 4,946770973 | 0,878543817 | 5,630647987 | 1,79534E-08 | 4,02302E-07 | SYT13 | protein_coding |
| ENSG00000021300 | 207,9633419 | 2,171349857 | 0,189154581 | 11,47923483 | 1,67759E-30 | 5,14187E-28 | PLEKHB1 | protein_coding |
| ENSG00000023445 | 41,60082792 | 2,301370939 | 0,372203082 | 6,183105549 | 6,28527E-10 | 1,92982E-08 | BIRC3 | protein_coding |
| ENSG00000027644 | 80,42153086 | 2,785695267 | 0,292407775 | 9,52674827 | 1,62287E-21 | 2,15049E-19 | INSRR | protein_coding |
| ENSG00000029534 | 158,9072004 | 2,320318483 | 0,245555836 | 9,449250016 | 3,41267E-21 | 4,35833E-19 | ANK1 | protein_coding |
| ENSG00000036530 | 55,10944722 | 2,87528239 | 0,324345948 | 8,864862992 | 7,65982E-19 | 7,7739E-17 | CYP46A1 | protein_coding |
| ENSG00000054179 | 78,94974214 | 2,704872889 | 0,269383671 | 10,04096825 | 1,00681E-23 | 1,68991E-21 | ENTPD2 | protein_coding |
| ENSG00000062038 | 123,7972813 | 2,335281723 | 0,213675182 | 10,92912007 | 8,36557E-28 | 2,16816E-25 | CDH3 | protein_coding |
| ENSG00000063127 | 104,0879335 | 2,087275013 | 0,264915092 | 7,87903398 | 3,29922E-15 | 2,23207E-13 | SLC6A16 | protein_coding |
| ENSG00000064300 | 429,5951477 | 3,382387612 | 0,17634418 | 19,18060242 | 5,3754E-82 | 1,89472E-78 | NGFR | protein_coding |
| ENSG00000065361 | 671,8656728 | 2,368899701 | 0,110530187 | 21,43215134 | 6,7011E-102 | 3,93669E-98 | ERBB3 | protein_coding |
| ENSG00000066923 | 196,013419 | 2,325605086 | 0,187229659 | 12,42113618 | 2,00691E-35 | 9,68732E-33 | STAG3 | protein_coding |
| ENSG00000067842 | 72,01482231 | 3,168467221 | 0,246159201 | 12,87161809 | 6,50255E-38 | 3,95175E-35 | ATP2B3 | protein_coding |
| ENSG00000072163 | 170,1484635 | 2,361081118 | 0,18023359 | 13,10011699 | 3,28751E-39 | 2,14589E-36 | LIMS2 | protein_coding |
| ENSG00000072818 | 63,80343939 | 2,732443982 | 0,276022348 | 9,899357803 | 4,18957E-23 | 6,77403E-21 | ACAP1 | protein_coding |
| ENSG00000073861 | 48,81304673 | 3,449996688 | 0,406771102 | 8,481420313 | 2,22461E-17 | 1,97018E-15 | TBX21 | protein_coding |
| ENSG00000074660 | 70,96936509 | 2,066043906 | 0,251207343 | 8,22445667 | 1,96078E-16 | 1,55661E-14 | SCARF1 | protein_coding |
| ENSG00000075673 | 44,15268178 | 3,329279145 | 0,375791891 | 8,859369315 | 8,04684E-19 | 8,12708E-17 | ATP12A | protein_coding |
| ENSG00000079308 | 651,7935611 | 3,539756081 | 0,130029878 | 27,22263624 | 3,5051E-163 | 1,2355E-158 | TNS1 | protein_coding |
| ENSG00000080031 | 21,58864789 | 3,492209398 | 0,602725552 | 5,794029118 | 6,87175E-09 | 1,70695E-07 | PTPRH | protein_coding |
| ENSG00000081479 | 86,53195937 | 2,479594674 | 0,276635991 | 8,963384196 | 3,14863E-19 | 3,36312E-17 | LRP2 | protein_coding |
| ENSG00000084636 | 35,074958 | 4,453977769 | 0,593376805 | 7,506154156 | 6,08897E-14 | 3,61929E-12 | COL16A1 | protein_coding |
| ENSG00000085117 | 117,9585234 | 2,100700562 | 0,220487174 | 9,527540877 | 1,61053E-21 | 2,14219E-19 | CD82 | protein_coding |
| ENSG00000087245 | 30,5706812 | 3,080989982 | 0,530530381 | 5,80737709 | 6,34591E-09 | 1,59585E-07 | MMP2 | protein_coding |
| ENSG00000088340 | 342,2642788 | 2,478367376 | 0,186977109 | 13,25492406 | 4,22544E-40 | 3,10288E-37 | FER1L4 | transcribed_unitary_pseudogene |
| ENSG00000088827 | 14,58811672 | 6,051051442 | 1,074193288 | 5,633112316 | 1,76986E-08 | 3,97604E-07 | SIGLEC1 | protein_coding |
| ENSG00000090339 | 181,7975527 | 3,549463772 | 0,199145895 | 17,82343426 | 4,64926E-71 | 1,09251E-67 | ICAM1 | protein_coding |
| ENSG00000091536 | 86,52467451 | 4,141895044 | 0,344943791 | 12,00744917 | 3,24707E-33 | 1,2717E-30 | MYO15A | protein_coding |
| ENSG00000092068 | 84,38062302 | 2,87903615 | 0,275368455 | 10,45521408 | 1,38685E-25 | 2,90975E-23 | SLC7A8 | protein_coding |
| ENSG00000095303 | 109,9574369 | 3,34181956 | 0,267628065 | 12,4868054 | 8,81215E-36 | 4,5016E-33 | PTGS1 | protein_coding |
| ENSG00000099994 | 228,0496234 | 2,459542349 | 0,182594138 | 13,46999623 | 2,34878E-41 | 1,83977E-38 | SUSD2 | protein_coding |
| ENSG00000100346 | 58,1955877 | 2,852836886 | 0,301269965 | 9,469370379 | 2,81536E-21 | 3,65318E-19 | CACNA1I | protein_coding |
| ENSG00000101198 | 13,8500675 | 6,87444816 | 1,223708596 | 5,617716656 | 1,93497E-08 | 4,27611E-07 | NKAIN4 | protein_coding |
| ENSG00000101203 | 419,0435605 | 7,36258565 | 0,377341824 | 19,51171372 | 8,73057E-85 | 3,84669E-81 | COL20A1 | protein_coding |
| ENSG00000101204 | 61,10926766 | 4,061232006 | 0,394428597 | 10,29649481 | 7,30756E-25 | 1,37742E-22 | CHRNA4 | protein_coding |
| ENSG00000101850 | 65,044158 | 2,124677125 | 0,229899154 | 9,241778776 | 2,42433E-20 | 2,84843E-18 | GPR143 | protein_coding |
| ENSG00000102445 | 28,03047282 | 4,50837597 | 0,605322046 | 7,447896536 | 9,48401E-14 | 5,4623E-12 | RUBCNL | protein_coding |
| ENSG00000103196 | 47,2828225 | 3,302203143 | 0,403098995 | 8,192040134 | 2,56835E-16 | 1,98965E-14 | CRISPLD2 | protein_coding |
| ENSG00000103489 | 21,05397307 | 4,335201265 | 0,659043865 | 6,578016268 | 4,76766E-11 | 1,84064E-09 | XYLT1 | protein_coding |
| ENSG00000104848 | 63,94631845 | 2,655014283 | 0,298528905 | 8,893659012 | 5,9129E-19 | 6,11196E-17 | KCNA7 | protein_coding |
| ENSG00000104856 | 141,0278826 | 2,293915202 | 0,219216497 | 10,46415407 | 1,26199E-25 | 2,67967E-23 | RELB | protein_coding |
| ENSG00000104870 | 33,0219089 | 2,613720695 | 0,449391476 | 5,816133228 | 6,02245E-09 | 1,52646E-07 | FCGRT | protein_coding |
| ENSG00000104888 | 160,2314889 | 4,175629337 | 0,288741219 | 14,46149374 | 2,12143E-47 | 2,49254E-44 | SLC17A7 | protein_coding |
| ENSG00000104967 | 260,3824365 | 2,039122793 | 0,178127978 | 11,4475155 | 2,41982E-30 | 7,35291E-28 | NOVA2 | protein_coding |
| ENSG00000105122 | 56,06307414 | 2,71742321 | 0,378283137 | 7,18356951 | 6,79145E-13 | 3,52037E-11 | RASAL3 | protein_coding |
| ENSG00000105357 | 53,33016608 | 7,824863674 | 1,034519691 | 7,563764849 | 3,91567E-14 | 2,3674E-12 | MYH14 | protein_coding |
| ENSG00000105559 | 170,0018099 | 2,1230631 | 0,199831475 | 10,62426776 | 2,29809E-26 | 5,25993E-24 | PLEKHA4 | protein_coding |
| ENSG00000105639 | 32,80124725 | 3,851014428 | 0,574956659 | 6,697921253 | 2,11405E-11 | 8,5947E-10 | JAK3 | protein_coding |
| ENSG00000107147 | 29,21054287 | 2,560727146 | 0,410155088 | 6,243314349 | 4,28395E-10 | 1,36405E-08 | KCNT1 | protein_coding |
| ENSG00000107738 | 81,07975052 | 3,411428862 | 0,315871602 | 10,80004925 | 3,44019E-27 | 8,42082E-25 | VSIR | protein_coding |
| ENSG00000107796 | 38,61148429 | 2,160539682 | 0,383224569 | 5,637790097 | 1,72246E-08 | 3,8869E-07 | ACTA2 | protein_coding |
| ENSG00000107859 | 20,53411021 | 3,323089213 | 0,540931762 | 6,143268794 | 8,08403E-10 | 2,43752E-08 | PITX3 | protein_coding |
| ENSG00000108176 | 59,84432933 | 2,414334286 | 0,334224508 | 7,223690143 | 5,05955E-13 | 2,66575E-11 | DNAJC12 | protein_coding |
| ENSG00000108370 | 44,34048549 | 5,137635414 | 0,632216361 | 8,12638794 | 4,42272E-16 | 3,30981E-14 | RGS9 | protein_coding |
| ENSG00000108387 | 149,3357064 | 2,286052287 | 0,206485901 | 11,071227 | 1,73015E-28 | 4,69111E-26 | SEPTIN4 | protein_coding |
| ENSG00000108846 | 62,03154257 | 2,78091928 | 0,293737345 | 9,467367125 | 2,86986E-21 | 3,70538E-19 | ABCC3 | protein_coding |
| ENSG00000109339 | 32,47301188 | 2,942550949 | 0,421489991 | 6,981306818 | 2,92447E-12 | 1,38365E-10 | MAPK10 | protein_coding |
| ENSG00000110169 | 75,48935649 | 2,224711865 | 0,253266698 | 8,784067867 | 1,57666E-18 | 1,56108E-16 | HPX | protein_coding |
| ENSG00000110436 | 51,50091762 | 3,29795318 | 0,385936066 | 8,545335541 | 1,28163E-17 | 1,16131E-15 | SLC1A2 | protein_coding |
| ENSG00000110786 | 19,75259327 | 4,598245908 | 0,791793114 | 5,807383051 | 6,34568E-09 | 1,59585E-07 | PTPN5 | protein_coding |
| ENSG00000112139 | 320,0026817 | 2,160074045 | 0,180603514 | 11,9603102 | 5,73473E-33 | 2,19715E-30 | MDGA1 | protein_coding |
| ENSG00000112182 | 5821,12613 | 5,452496993 | 0,235814357 | 23,12198911 | 2,7827E-118 | 4,9042E-114 | BACH2 | protein_coding |
| ENSG00000113140 | 75,49491918 | 2,869546255 | 0,411171226 | 6,978956874 | 2,9738E-12 | 1,4051E-10 | SPARC | protein_coding |
| ENSG00000115380 | 19,86409129 | 4,481210768 | 0,584332593 | 7,668938578 | 1,73425E-14 | 1,07621E-12 | EFEMP1 | protein_coding |
| ENSG00000115423 | 52,50804759 | 2,633369363 | 0,317148321 | 8,303273859 | 1,01281E-16 | 8,2257E-15 | DNAH6 | protein_coding |
| ENSG00000115596 | 42,17552152 | 4,170244239 | 0,463490356 | 8,997477915 | 2,30962E-19 | 2,48959E-17 | WNT6 | protein_coding |
| ENSG00000115828 | 65,09475549 | 4,104362053 | 0,430531675 | 9,533240631 | 1,52449E-21 | 2,03542E-19 | QPCT | protein_coding |
| ENSG00000116299 | 49,06309292 | 2,819049515 | 0,29801575 | 9,459397761 | 3,09722E-21 | 3,98433E-19 | ELAPOR1 | protein_coding |
| ENSG00000116981 | 66,65257918 | 2,186445564 | 0,270281098 | 8,089524492 | 5,98981E-16 | 4,4077E-14 | NT5C1A | protein_coding |
| ENSG00000116983 | 186,6042251 | 2,819039646 | 0,221373415 | 12,73431882 | 3,81158E-37 | 2,20247E-34 | HPCAL4 | protein_coding |
| ENSG00000117115 | 241,0472994 | 2,426481542 | 0,204868169 | 11,844112 | 2,30851E-32 | 8,21921E-30 | PADI2 | protein_coding |
| ENSG00000117971 | 62,58754707 | 2,140271363 | 0,287084125 | 7,455206243 | 8,97274E-14 | 5,18477E-12 | CHRNB4 | protein_coding |
| ENSG00000118160 | 44,13915242 | 4,234385389 | 0,484709584 | 8,735922561 | 2,41676E-18 | 2,33386E-16 | SLC8A2 | protein_coding |
| ENSG00000119922 | 62,44794168 | 2,57072432 | 0,321089664 | 8,006250626 | 1,18259E-15 | 8,47231E-14 | IFIT2 | protein_coding |
| ENSG00000121101 | 59,48247272 | 2,04453132 | 0,288116865 | 7,096187575 | 1,28245E-12 | 6,38474E-11 | TEX14 | protein_coding |
| ENSG00000122176 | 14,38398387 | 3,864665466 | 0,694587473 | 5,563972311 | 2,63702E-08 | 5,67112E-07 | FMOD | protein_coding |
| ENSG00000122733 | 60,94914948 | 3,242032923 | 0,318527024 | 10,17820365 | 2,48099E-24 | 4,41666E-22 | PHF24 | protein_coding |
| ENSG00000122735 | 81,85834923 | 2,006889823 | 0,219633124 | 9,137464262 | 6,39338E-20 | 7,22288E-18 | DNAI1 | protein_coding |

| | | | | | | | | |
|---|---|---|---|---|---|---|---|---|
| ENSG00000122986 | 85,49434103 | 2,558348301 | 0,21873288 | 11,69622188 | 1,33255E-31 | 4,47332E-29 | HVCN1 | protein_coding |
| ENSG00000123095 | 22,18488543 | 2,337644463 | 0,428474154 | 5,455742054 | 4,87687E-08 | 9,82848E-07 | BHLHE41 | protein_coding |
| ENSG00000123453 | 57,41692086 | 4,801004613 | 0,480447389 | 9,992779073 | 1,6392E-23 | 2,72541E-21 | SARDH | protein_coding |
| ENSG00000124466 | 18,20793617 | 3,411682165 | 0,58659353 | 5,816092389 | 6,02392E-09 | 1,52646E-07 | LYPD3 | protein_coding |
| ENSG00000124772 | 128,4547558 | 3,411899468 | 0,252788317 | 13,49706155 | 1,62738E-41 | 1,30368E-38 | CPNE5 | protein_coding |
| ENSG00000125531 | 24,09089 | 3,000233406 | 0,449970157 | 6,667627532 | 2,59972E-11 | 1,04487E-09 | FNDC11 | protein_coding |
| ENSG00000125637 | 149,1336634 | 2,202100609 | 0,196809143 | 11,18901578 | 4,61528E-29 | 1,3226E-26 | PSD4 | protein_coding |
| ENSG00000126856 | 11,21499846 | 3,443046483 | 0,607090515 | 5,67138902 | 1,41644E-08 | 3,26319E-07 | PRDM7 | protein_coding |
| ENSG00000128340 | 38,41452422 | 2,423618161 | 0,359640857 | 6,738995618 | 1,59485E-11 | 6,62055E-10 | RAC2 | protein_coding |
| ENSG00000129009 | 21,70462098 | 2,076069469 | 0,378533207 | 5,48451082 | 4,14615E-08 | 8,48192E-07 | ISLR | protein_coding |
| ENSG00000129244 | 195,248251 | 2,777970826 | 0,187847728 | 14,78841853 | 1,73994E-49 | 2,2594E-46 | ATP1B2 | protein_coding |
| ENSG00000129910 | 32,90903158 | 4,371036325 | 0,603617095 | 7,241405792 | 4,44058E-13 | 2,36795E-11 | CDH15 | protein_coding |
| ENSG00000129993 | 41,92189405 | 2,572761723 | 0,318981593 | 8,065549161 | 7,29074E-16 | 5,28774E-14 | CBFA2T3 | protein_coding |
| ENSG00000130635 | 16,67456414 | 5,282305956 | 0,659995872 | 8,003543928 | 1,20889E-15 | 8,60824E-14 | COL5A1 | protein_coding |
| ENSG00000130720 | 19,73590948 | 4,092769823 | 0,726609883 | 5,632692204 | 1,77418E-08 | 3,97833E-07 | FIBCD1 | protein_coding |
| ENSG00000131044 | 38,10114478 | 2,077194772 | 0,305304643 | 6,803678941 | 1,01981E-11 | 4,3677E-10 | TTLL9 | protein_coding |
| ENSG00000131095 | 25,39295638 | 2,631123852 | 0,466416161 | 5,641150703 | 1,68917E-08 | 3,81912E-07 | GFAP | protein_coding |
| ENSG00000131398 | 84,5904858 | 2,012327611 | 0,219293593 | 9,176408585 | 4,45709E-20 | 5,10076E-18 | KCNC3 | protein_coding |
| ENSG00000131409 | 93,08016815 | 2,631330833 | 0,211842142 | 12,42118687 | 2,00564E-35 | 9,68732E-33 | LRRC4B | protein_coding |
| ENSG00000131620 | 14,81946826 | 4,530828167 | 0,805392206 | 5,625617098 | 1,84846E-08 | 4,1222E-07 | ANO1 | protein_coding |
| ENSG00000131831 | 30,39697986 | 2,430437012 | 0,396808593 | 6,124960636 | 9,0706E-10 | 2,7072E-08 | RAI2 | protein_coding |
| ENSG00000132031 | 24,177042 | 2,409677244 | 0,435489933 | 5,533255908 | 3,1434E-08 | 6,62275E-07 | MATN3 | protein_coding |
| ENSG00000132470 | 325,6261233 | 2,345054367 | 0,151808677 | 15,44743297 | 7,85084E-54 | 1,25785E-50 | ITGB4 | protein_coding |
| ENSG00000132669 | 91,33079032 | 2,164025587 | 0,273809731 | 7,9033918 | 2,71415E-15 | 1,85764E-13 | RIN2 | protein_coding |
| ENSG00000132821 | 134,3298519 | 3,129205862 | 0,211628317 | 14,78632872 | 1,7948E-49 | 2,2594E-46 | VSTM2L | protein_coding |
| ENSG00000134202 | 133,3099966 | 2,402494154 | 0,220375095 | 10,9018406 | 1,12949E-27 | 2,90601E-25 | GSTM3 | protein_coding |
| ENSG00000134668 | 29,04793169 | 2,194553771 | 0,394346039 | 5,565045806 | 2,62084E-08 | 5,6432E-07 | SPOCD1 | protein_coding |
| ENSG00000135253 | 52,53387293 | 2,709942009 | 0,384833786 | 7,041850557 | 1,89703E-12 | 9,18497E-11 | KCP | protein_coding |
| ENSG00000135472 | 151,1634624 | 2,860748949 | 0,222008377 | 12,88577029 | 5,41323E-38 | 3,40724E-35 | FAIM2 | protein_coding |
| ENSG00000135636 | 111,6216189 | 2,971010278 | 0,246311222 | 12,06201751 | 1,67626E-33 | 6,79134E-31 | DYSF | protein_coding |
| ENSG00000135929 | 30,65981492 | 2,070995547 | 0,37027617 | 5,593110529 | 2,23037E-08 | 4,86787E-07 | CYP27A1 | protein_coding |
| ENSG00000136931 | 70,29593293 | 2,996504268 | 0,322551675 | 9,289997544 | 1,54292E-20 | 1,84355E-18 | NR5A1 | protein_coding |
| ENSG00000137474 | 33,00823762 | 5,769736127 | 0,878854659 | 6,56506291 | 5,20109E-11 | 1,98837E-09 | MYO7A | protein_coding |
| ENSG00000139567 | 43,82437808 | 5,901797735 | 0,86906817 | 6,790949133 | 1,11398E-11 | 4,73651E-10 | ACVRL1 | protein_coding |
| ENSG00000139973 | 32,67936719 | 3,053674618 | 0,521551739 | 5,85497927 | 4,7707E-09 | 1,23736E-07 | SYT16 | protein_coding |
| ENSG00000140279 | 23,80177759 | 2,425229181 | 0,430966238 | 5,627422677 | 1,82922E-08 | 4,09114E-07 | DUOX2 | protein_coding |
| ENSG00000141314 | 83,81559369 | 3,149352249 | 0,25741412 | 12,23457457 | 2,03167E-34 | 8,95152E-32 | RHBDL3 | protein_coding |
| ENSG00000141485 | 62,41351389 | 3,528032386 | 0,39680897 | 8,891009655 | 6,05561E-19 | 6,24118E-17 | SLC13A5 | protein_coding |
| ENSG00000141505 | 65,37748526 | 2,393533849 | 0,279673948 | 8,558301083 | 1,14543E-17 | 1,04326E-15 | ASGR1 | protein_coding |
| ENSG00000141837 | 71,50100902 | 6,080258481 | 0,585243373 | 10,38928206 | 2,77434E-25 | 5,55624E-23 | CACNA1A | protein_coding |
| ENSG00000142233 | 39,12327797 | 4,753762628 | 0,519450066 | 9,15152954 | 5,61331E-20 | 6,36199E-18 | NTN5 | protein_coding |
| ENSG00000142606 | 24,77043762 | 2,995854299 | 0,481097636 | 6,227123298 | 4,75077E-10 | 1,49781E-08 | MMEL1 | protein_coding |
| ENSG00000142609 | 171,8405596 | 3,602534436 | 0,196920377 | 18,29437104 | 9,1754E-75 | 2,3101E-71 | CFAP74 | protein_coding |
| ENSG00000142621 | 18,54460559 | 4,737013954 | 0,799157796 | 5,927507657 | 3,07567E-09 | 8,30101E-08 | FHAD1 | protein_coding |
| ENSG00000142677 | 24,77013325 | 3,104270328 | 0,565475273 | 5,489665911 | 4,02695E-08 | 8,26204E-07 | IL22RA1 | protein_coding |
| ENSG00000143195 | 87,79582677 | 3,720088151 | 0,36384425 | 10,22439725 | 1,54184E-24 | 2,78702E-22 | ILDR2 | protein_coding |
| ENSG00000143382 | 753,1059867 | 2,374698799 | 0,12369987 | 19,19726195 | 3,90122E-82 | 1,52789E-78 | ADAMTSL4 | protein_coding |
| ENSG00000143416 | 476,5570321 | 10,78321163 | 1,026142257 | 10,50849582 | 7,89426E-26 | 1,7071E-23 | SELENBP1 | protein_coding |
| ENSG00000143502 | 120,4820064 | 2,313474435 | 0,194945725 | 11,86727451 | 1,75079E-32 | 6,36204E-30 | SUSD4 | protein_coding |
| ENSG00000143603 | 306,6005563 | 2,489798093 | 0,176803097 | 14,08232176 | 4,87804E-45 | 4,91261E-42 | KCNN3 | protein_coding |
| ENSG00000143850 | 242,2820346 | 4,161801037 | 0,236454755 | 17,60083461 | 2,42721E-69 | 5,34714E-66 | PLEKHA6 | protein_coding |
| ENSG00000143851 | 51,14985771 | 2,796359557 | 0,326850353 | 8,555473569 | 1,17386E-17 | 1,0664E-15 | PTPN7 | protein_coding |
| ENSG00000144891 | 30,93196936 | 3,771750098 | 0,538276182 | 7,007090831 | 2,43324E-12 | 1,16531E-10 | AGTR1 | protein_coding |
| ENSG00000145526 | 34,16233112 | 3,573614759 | 0,562159657 | 6,356939204 | 2,05813E-10 | 6,94212E-09 | CDH18 | protein_coding |
| ENSG00000145794 | 42,54085402 | 3,174030809 | 0,392310179 | 8,09061549 | 5,9364E-16 | 4,37753E-14 | MEGF10 | protein_coding |
| ENSG00000146005 | 53,5039062 | 2,136830151 | 0,299669073 | 7,130632902 | 9,99085E-13 | 5,03082E-11 | PSD2 | protein_coding |
| ENSG00000148357 | 138,5209186 | 4,849398242 | 0,341523686 | 14,19930285 | 9,25283E-46 | 9,59246E-43 | HMCN2 | protein_coding |
| ENSG00000148848 | 47,87005454 | 2,76191806 | 0,288724645 | 9,565924185 | 1,11203E-21 | 1,49607E-19 | ADAM12 | protein_coding |
| ENSG00000149260 | 516,7088159 | 2,744313424 | 0,14039464 | 19,54713822 | 4,3632E-85 | 2,19706E-81 | CAPN5 | protein_coding |
| ENSG00000149654 | 35,15108547 | 3,088246025 | 0,455428484 | 6,780968109 | 1,19373E-11 | 5,06338E-10 | CDH22 | protein_coding |
| ENSG00000154783 | 27,56954341 | 2,666753044 | 0,42821097 | 6,227661666 | 4,73448E-10 | 1,49401E-08 | FGD5 | protein_coding |
| ENSG00000155980 | 35,76339107 | 2,852147539 | 0,494656665 | 5,765913497 | 8,12167E-09 | 1,988E-07 | KIF5A | protein_coding |
| ENSG00000156042 | 198,0241579 | 2,031687069 | 0,171841387 | 11,82303692 | 2,96759E-32 | 1,04602E-29 | CFAP70 | protein_coding |
| ENSG00000156804 | 117,5358332 | 2,056392013 | 0,197953505 | 10,38825768 | 2,80429E-25 | 5,5845E-23 | FBXO32 | protein_coding |
| ENSG00000156966 | 22,45594777 | 2,668768562 | 0,42402053 | 6,293960725 | 3,09466E-10 | 1,00814E-08 | B3GNT7 | protein_coding |
| ENSG00000157087 | 55,70403505 | 3,58587652 | 0,347008794 | 10,33367622 | 4,96216E-25 | 9,82619E-23 | ATP2B2 | protein_coding |
| ENSG00000157103 | 37,79162273 | 3,825217268 | 0,422739609 | 9,048637003 | 1,44764E-19 | 1,59957E-17 | SLC6A1 | protein_coding |
| ENSG00000157423 | 33,53042305 | 2,458923103 | 0,432528421 | 5,684997754 | 1,30814E-08 | 3,03352E-07 | HYDIN | protein_coding |
| ENSG00000158445 | 34,6049428 | 2,523877315 | 0,354837737 | 7,112764667 | 1,13741E-12 | 5,70291E-11 | KCNB1 | protein_coding |
| ENSG00000158825 | 29,59030788 | 2,512611895 | 0,434597786 | 5,781465018 | 7,40529E-09 | 1,83045E-07 | CDA | protein_coding |
| ENSG00000159708 | 30,24040437 | 2,542361031 | 0,421263871 | 6,035079689 | 1,58884E-09 | 4,52371E-08 | LRRC36 | protein_coding |
| ENSG00000160111 | 47,09603347 | 2,787886487 | 0,42616192 | 6,541847956 | 6,07633E-11 | 2,29805E-09 | CPAMD8 | protein_coding |
| ENSG00000160179 | 18,30693674 | 5,839399871 | 1,060334219 | 5,507131401 | 3,64728E-08 | 7,57568E-07 | ABCG1 | protein_coding |
| ENSG00000160460 | 164,1581496 | 2,647382151 | 0,181731204 | 14,56757064 | 4,51669E-48 | 5,48981E-45 | SPTBN4 | protein_coding |
| ENSG00000160539 | 69,62110148 | 3,497247072 | 0,352059602 | 9,933678999 | 2,97087E-23 | 4,84803E-21 | PLPP7 | protein_coding |
| ENSG00000160712 | 338,4667966 | 2,125738946 | 0,181830115 | 11,69079692 | 1,42047E-31 | 4,72348E-29 | IL6R | protein_coding |
| ENSG00000160808 | 19,47482869 | 4,114420318 | 0,607901337 | 6,768236998 | 1,30361E-11 | 5,50296E-10 | MYL3 | protein_coding |
| ENSG00000161082 | 65,2355563 | 2,194723399 | 0,267515912 | 8,204085442 | 2,32353E-16 | 1,81194E-14 | CELF5 | protein_coding |
| ENSG00000161270 | 21,53136647 | 2,893310331 | 0,526813653 | 5,492094437 | 3,97195E-08 | 8,16346E-07 | NPHS1 | protein_coding |
| ENSG00000161405 | 36,42927008 | 2,054285137 | 0,344112344 | 5,969809483 | 2,37531E-09 | 6,50038E-08 | IKZF3 | protein_coding |
| ENSG00000161642 | 254,3086608 | 2,846307918 | 0,19922249 | 14,28708134 | 2,6342E-46 | 2,81365E-43 | ZNF385A | protein_coding |
| ENSG00000161681 | 243,6822011 | 2,223060011 | 0,164295383 | 13,53087331 | 1,02795E-41 | 8,42633E-39 | SHANK1 | protein_coding |
| ENSG00000162426 | 33,17659478 | 2,001647087 | 0,326005392 | 6,139920194 | 8,25629E-10 | 2,48309E-08 | SLC45A1 | protein_coding |
| ENSG00000162745 | 129,9929893 | 2,022525653 | 0,219396273 | 9,218596213 | 3,01015E-20 | 3,49019E-18 | OLFML2B | protein_coding |
| ENSG00000163075 | 48,0094885 | 2,672755525 | 0,338910475 | 7,886317252 | 3,11235E-15 | 2,10969E-13 | CFAP221 | protein_coding |
| ENSG00000163235 | 64,59063083 | 3,436996966 | 0,33293067 | 10,32346153 | 5,51965E-25 | 1,0749E-22 | TGFA | protein_coding |
| ENSG00000163239 | 40,12742841 | 2,901404644 | 0,367246883 | 7,90042007 | 2,77965E-15 | 1,89878E-13 | TDRD10 | protein_coding |
| ENSG00000163462 | 62,29061386 | 2,003002317 | 0,268600886 | 7,45716943 | 8,8401E-14 | 5,11652E-12 | TRIM46 | protein_coding |
| ENSG00000163520 | 107,4344374 | 3,575883754 | 0,300855962 | 11,88570015 | 1,40453E-32 | 5,21126E-30 | FBLN2 | protein_coding |

| | | | | | | | | |
|---|---|---|---|---|---|---|---|---|
| ENSG00000163637 | 26,71897658 | 2,723782185 | 0,462182837 | 5,89330015 | 3,78558E-09 | 1,00607E-07 | PRICKLE2 | protein_coding |
| ENSG00000166428 | 43,44224647 | 2,358073317 | 0,315694963 | 7,469467666 | 8,05199E-14 | 4,70674E-12 | PLD4 | protein_coding |
| ENSG00000166501 | 77,64726001 | 2,037440344 | 0,264559702 | 7,701249763 | 1,34742E-14 | 8,46592E-13 | PRKCB | protein_coding |
| ENSG00000166546 | 78,01003981 | 2,330563403 | 0,263757894 | 8,835994882 | 9,92113E-19 | 9,93466E-17 | BEAN1 | protein_coding |
| ENSG00000166897 | 143,5448493 | 2,026360409 | 0,193018853 | 10,49825124 | 8,79952E-26 | 1,89125E-23 | ELFN2 | protein_coding |
| ENSG00000167080 | 36,11620095 | 3,367076363 | 0,436946269 | 7,705927709 | 1,29896E-14 | 8,19066E-13 | B4GALNT2 | protein_coding |
| ENSG00000168016 | 95,5553875 | 2,660591828 | 0,233781091 | 11,38069728 | 5,21808E-30 | 1,54561E-27 | TRANK1 | protein_coding |
| ENSG00000168070 | 21,82112092 | 2,84036902 | 0,463295514 | 6,130793265 | 8,7442E-10 | 2,62397E-08 | MAJIN | protein_coding |
| ENSG00000168071 | 207,1734025 | 3,12440749 | 0,200884831 | 15,55322758 | 1,51286E-54 | 2,66626E-51 | CCDC88B | protein_coding |
| ENSG00000168477 | 126,1223942 | 2,979552736 | 0,236481822 | 12,59950009 | 2,12487E-36 | 1,10143E-33 | TNXB | protein_coding |
| ENSG00000168546 | 17,14167412 | 4,378271266 | 0,783467605 | 5,588324566 | 2,29271E-08 | 4,98233E-07 | GFRA2 | protein_coding |
| ENSG00000169064 | 23,59455979 | 2,775817646 | 0,454208034 | 6,111335418 | 9,88009E-10 | 2,9265E-08 | ZBBX | protein_coding |
| ENSG00000169181 | 61,97330386 | 8,032831351 | 1,035675427 | 7,75612817 | 8,75615E-15 | 5,6218E-13 | GSG1L | protein_coding |
| ENSG00000169220 | 95,60908272 | 2,141163065 | 0,236672167 | 9,046957634 | 1,47007E-19 | 1,61928E-17 | RGS14 | protein_coding |
| ENSG00000169291 | 128,2900061 | 2,273287914 | 0,201389202 | 11,28803278 | 1,50364E-29 | 4,3443E-27 | SHE | protein_coding |
| ENSG00000169994 | 24,28834931 | 3,469514574 | 0,519942655 | 6,67287929 | 2,50833E-11 | 1,01044E-09 | MYO7B | protein_coding |
| ENSG00000170421 | 222,2503444 | 2,091213236 | 0,164285819 | 12,72911596 | 4,07426E-37 | 2,31628E-34 | KRT8 | protein_coding |
| ENSG00000170442 | 13,23024352 | 4,135022446 | 0,74379836 | 5,559332564 | 2,70808E-08 | 5,80977E-07 | KRT86 | protein_coding |
| ENSG00000170703 | 37,55692612 | 2,450991776 | 0,366680262 | 6,68427519 | 2,3207E-11 | 9,41314E-10 | TTLL6 | protein_coding |
| ENSG00000171992 | 168,4756059 | 3,061249373 | 0,217501498 | 14,07461285 | 5,4402E-45 | 5,32656E-42 | SYNPO | protein_coding |
| ENSG00000172005 | 30,09470292 | 3,832327389 | 0,556265259 | 6,889388346 | 5,60328E-12 | 2,52241E-10 | MAL | protein_coding |
| ENSG00000172350 | 158,7075642 | 2,325155126 | 0,192877267 | 12,05510201 | 1,82307E-33 | 7,30221E-31 | ABCG4 | protein_coding |
| ENSG00000172794 | 24,48212037 | 3,162237377 | 0,554980562 | 5,697924565 | 1,21275E-08 | 2,82905E-07 | RAB37 | protein_coding |
| ENSG00000172818 | 25,52282361 | 4,292720897 | 0,615617974 | 6,973027233 | 3,10193E-12 | 1,45782E-10 | OVOL1 | protein_coding |
| ENSG00000173210 | 50,74359587 | 2,391762013 | 0,332832725 | 7,186078273 | 6,66788E-13 | 3,46141E-11 | ABLIM3 | protein_coding |
| ENSG00000173546 | 642,8241115 | 2,650119553 | 0,144106667 | 18,3899857 | 1,58022E-75 | 4,64162E-72 | CSPG4 | protein_coding |
| ENSG00000173641 | 19,88210274 | 3,042468476 | 0,506108368 | 6,011496092 | 1,83819E-09 | 5,15864E-08 | HSPB7 | protein_coding |
| ENSG00000173698 | 181,8311626 | 2,140466654 | 0,178817024 | 11,97015033 | 5,09363E-33 | 1,97297E-30 | ADGRG2 | protein_coding |
| ENSG00000173838 | 14,62799024 | 3,637006617 | 0,588119527 | 6,184128313 | 6,24466E-10 | 1,91902E-08 | MARCHF10 | protein_coding |
| ENSG00000174292 | 197,7667965 | 2,061497789 | 0,168758262 | 12,21568509 | 2,56333E-34 | 1,10186E-31 | TNK1 | protein_coding |
| ENSG00000174348 | 31,01562375 | 3,351055361 | 0,488434033 | 6,86081464 | 6,8469E-12 | 3,03677E-10 | PODN | protein_coding |
| ENSG00000174358 | 13,61722985 | 4,924339147 | 0,795556715 | 6,189802759 | 6,02395E-10 | 1,86093E-08 | SLC6A19 | protein_coding |
| ENSG00000174403 | 52,80696112 | 2,318961429 | 0,321166461 | 7,22043461 | 5,18217E-13 | 2,72628E-11 | MIR1-1HG-AS1 | lncRNA |
| ENSG00000174460 | 258,9302181 | 2,993599606 | 0,286918205 | 10,43363422 | 1,74099E-25 | 3,6098E-23 | ZCCHC12 | protein_coding |
| ENSG00000174844 | 32,17621975 | 2,915135856 | 0,411850772 | 7,078136187 | 1,46106E-12 | 7,19266E-11 | DNAH12 | protein_coding |
| ENSG00000175147 | 23,23196597 | 2,359904644 | 0,406168003 | 5,810168759 | 6,24099E-09 | 1,57581E-07 | TMEM51-AS1 | lncRNA |
| ENSG00000175170 | 90,79893133 | 6,299254891 | 0,534672306 | 11,78152454 | 4,86059E-32 | 1,67967E-29 | FAM182B | lncRNA |
| ENSG00000175264 | 31,28823814 | 3,907579572 | 0,550317817 | 7,100587067 | 1,24228E-12 | 6,20225E-11 | CHST1 | protein_coding |
| ENSG00000175591 | 169,0977583 | 3,068620641 | 0,17862118 | 17,17948926 | 3,78226E-66 | 7,84219E-63 | P2RY2 | protein_coding |
| ENSG00000175985 | 32,51268074 | 2,543085268 | 0,390629882 | 6,510216916 | 7,50424E-11 | 2,76683E-09 | PLEKHD1 | protein_coding |
| ENSG00000177098 | 244,1807548 | 3,63987941 | 0,19586921 | 18,58321384 | 4,39386E-77 | 1,40795E-73 | SCN4B | protein_coding |
| ENSG00000177103 | 29,69341852 | 6,994040771 | 1,0479583 | 6,673968584 | 2,48977E-11 | 1,00411E-09 | DSCAML1 | protein_coding |
| ENSG00000177108 | 49,09942928 | 2,436205782 | 0,363490185 | 6,702260158 | 2,0522E-11 | 8,37266E-10 | ZDHHC22 | protein_coding |
| ENSG00000177301 | 72,10759455 | 3,551932312 | 0,356095188 | 9,97467091 | 1,96753E-23 | 3,22564E-21 | KCNA2 | protein_coding |
| ENSG00000177692 | 29,7365375 | 2,440023199 | 0,412199211 | 5,919524188 | 3,22874E-09 | 8,66769E-08 | DNAJC28 | protein_coding |
| ENSG00000179057 | 36,24220304 | 2,491883578 | 0,320385416 | 7,777768442 | 7,3815E-15 | 4,81975E-13 | IGSF22 | protein_coding |
| ENSG00000179388 | 105,1878836 | 2,638353058 | 0,472650694 | 5,582035717 | 2,3772E-08 | 5,15322E-07 | EGR3 | protein_coding |
| ENSG00000179630 | 24,25067493 | 2,918884113 | 0,47066245 | 6,2016507 | 5,5874E-10 | 1,73826E-08 | LACC1 | protein_coding |
| ENSG00000179930 | 20,99924027 | 4,653652501 | 0,773757974 | 6,01435159 | 1,80608E-09 | 5,08067E-08 | ZNF648 | protein_coding |
| ENSG00000180638 | 160,3207462 | 3,08426784 | 0,168086664 | 18,34927149 | 3,34568E-75 | 9,07143E-72 | SLC47A2 | protein_coding |
| ENSG00000180712 | 25,56282506 | 2,989756353 | 0,416524329 | 7,177867282 | 7,08072E-13 | 3,64352E-11 | LINC02363 | lncRNA |
| ENSG00000182040 | 63,42443178 | 2,993999423 | 0,337594888 | 8,868615982 | 7,40606E-19 | 7,56663E-17 | USH1G | protein_coding |
| ENSG00000182256 | 45,64411068 | 2,012563127 | 0,260150848 | 7,736139023 | 1,02481E-14 | 6,50858E-13 | GABRG3 | protein_coding |
| ENSG00000183508 | 71,91176717 | 3,003318322 | 0,320002813 | 9,385287254 | 6,27452E-21 | 7,73301E-19 | TENT5C | protein_coding |
| ENSG00000183682 | 37,10800662 | 2,393408291 | 0,356915335 | 6,705815239 | 2,00285E-11 | 8,18984E-10 | BMP8A | protein_coding |
| ENSG00000183914 | 93,0181299 | 3,959403709 | 0,317280862 | 12,47917599 | 9,69851E-36 | 4,88362E-33 | DNAH2 | protein_coding |
| ENSG00000184185 | 27,59986107 | 2,528171434 | 0,419883017 | 6,021132863 | 1,73201E-09 | 4,8918E-08 | KCNJ12 | protein_coding |
| ENSG00000184454 | 33,60086497 | 4,016985311 | 0,493216492 | 8,144466731 | 3,80957E-16 | 2,89396E-14 | NCMAP | protein_coding |
| ENSG00000184497 | 46,24431071 | 2,205949215 | 0,311077271 | 7,091322375 | 1,32836E-12 | 6,59468E-11 | TMEM255B | protein_coding |
| ENSG00000184574 | 16,78578139 | 3,268315488 | 0,546879055 | 5,976304012 | 2,28257E-09 | 6,27093E-08 | LPAR5 | protein_coding |
| ENSG00000184584 | 98,85294963 | 2,110536288 | 0,243234584 | 8,676958064 | 4,06495E-18 | 3,84132E-16 | STING1 | protein_coding |
| ENSG00000184828 | 31,81840414 | 2,015821644 | 0,365602111 | 5,513703507 | 3,5136E-08 | 7,31527E-07 | ZBTB7C | protein_coding |
| ENSG00000184985 | 37,11313988 | 2,295689725 | 0,317999869 | 7,219153043 | 5,23124E-13 | 2,74391E-11 | SORCS2 | protein_coding |
| ENSG00000185101 | 130,0980583 | 2,622916342 | 0,238404716 | 11,00194822 | 3,73964E-28 | 9,83692E-26 | ANO9 | protein_coding |
| ENSG00000185513 | 80,55294616 | 2,333153177 | 0,298964514 | 7,804114083 | 5,9921E-15 | 3,9701E-13 | L3MBTL1 | protein_coding |
| ENSG00000185567 | 1345,784646 | 3,199699886 | 0,141480443 | 22,61584586 | 3,0267E-113 | 3,5562E-109 | AHNAK2 | protein_coding |
| ENSG00000185974 | 21,45522505 | 4,176730859 | 0,554469902 | 7,532836036 | 4,96499E-14 | 2,96621E-12 | GRK1 | protein_coding |
| ENSG00000186409 | 75,52118562 | 2,130411217 | 0,245652257 | 8,67246752 | 4,22855E-18 | 3,98524E-16 | CCDC30 | protein_coding |
| ENSG00000186510 | 37,96503529 | 3,859651121 | 0,468521162 | 8,23794405 | 1,75195E-16 | 1,40347E-14 | CLCNKA | protein_coding |
| ENSG00000186642 | 110,1477549 | 2,967938996 | 0,254912815 | 11,6429572 | 2,49222E-31 | 8,20988E-29 | PDE2A | protein_coding |
| ENSG00000187122 | 491,2863301 | 2,116482603 | 0,132510331 | 15,97220822 | 1,99593E-57 | 3,90847E-54 | SLIT1 | protein_coding |
| ENSG00000187416 | 8,367705779 | 7,107808684 | 1,256044286 | 5,658883817 | 1,52361E-08 | 3,48448E-07 | LHFPL3 | protein_coding |
| ENSG00000187554 | 32,48806765 | 2,012393583 | 0,350402445 | 5,743092304 | 9,2963E-09 | 2,23455E-07 | TLR5 | protein_coding |
| ENSG00000187678 | 25,59153587 | 2,556005721 | 0,46704808 | 5,472682212 | 4,43275E-08 | 9,0211E-07 | SPRY4 | protein_coding |
| ENSG00000187800 | 90,42599617 | 3,146125146 | 0,28915623 | 10,88036438 | 1,4299E-27 | 3,65226E-25 | PEAR1 | protein_coding |
| ENSG00000187902 | 24,21213633 | 2,811543247 | 0,477654959 | 5,886138511 | 3,95323E-09 | 1,04691E-07 | SHISA7 | protein_coding |
| ENSG00000187997 | 30,01185762 | 2,289284462 | 0,37407856 | 6,119795969 | 9,36952E-10 | 2,78698E-08 | C17orf99 | protein_coding |
| ENSG00000188162 | 80,73670584 | 6,733100716 | 0,621025859 | 10,84190073 | 2,17896E-27 | 5,4471E-25 | OTOG | protein_coding |
| ENSG00000188263 | 17,9402011 | 3,53330525 | 0,61707504 | 5,72589235 | 1,02891E-08 | 2,44059E-07 | IL17REL | protein_coding |
| ENSG00000188389 | 42,5837011 | 3,469377005 | 0,381331048 | 9,098071135 | 9,19503E-20 | 1,02891E-17 | PDCD1 | protein_coding |
| ENSG00000188596 | 88,37659924 | 3,10021843 | 0,287091944 | 10,79869531 | 3,49129E-27 | 8,48697E-25 | CFAP54 | protein_coding |
| ENSG00000188672 | 32,22698711 | 2,64334468 | 0,374219867 | 7,063613973 | 1,62227E-12 | 7,90896E-11 | RHCE | protein_coding |
| ENSG00000188783 | 19,84793134 | 3,1527466 | 0,528105196 | 5,969921569 | 2,37368E-09 | 6,50038E-08 | PRELP | protein_coding |
| ENSG00000189120 | 32,01089699 | 4,502199812 | 0,567013057 | 7,940204828 | 2,01848E-15 | 1,40054E-13 | SP6 | protein_coding |
| ENSG00000196092 | 60,21736558 | 3,49793104 | 0,379108869 | 9,226719082 | 2,79048E-20 | 3,25692E-18 | PAX5 | protein_coding |
| ENSG00000196169 | 81,5569693 | 3,328853495 | 0,3125313 | 10,65126434 | 1,72013E-26 | 3,98888E-24 | KIF19 | protein_coding |
| ENSG00000196337 | 29,55256542 | 2,104197249 | 0,377075743 | 5,580303927 | 2,40099E-08 | 5,2016E-07 | CGB7 | protein_coding |
| ENSG00000196668 | 25,42454525 | 2,957507746 | 0,45408702 | 6,513085852 | 7,36224E-11 | 2,72588E-09 | LINC00173 | lncRNA |
| ENSG00000197177 | 46,26780958 | 3,935316276 | 0,383788543 | 10,25386595 | 1,13703E-24 | 2,08739E-22 | ADGRA1 | protein_coding |

| | | | | | | | | |
|---|---|---|---|---|---|---|---|---|
| ENSG00000197467 | 75,4480125 | 2,061762126 | 0,252657772 | 8,160295689 | 3,34205E-16 | 2,5498E-14 | COL13A1 | protein_coding |
| ENSG00000197558 | 349,4068344 | 2,779064944 | 0,193122942 | 14,39013365 | 5,96805E-47 | 6,5738E-44 | SSPOP | transcribed_unitary_pseudogene |
| ENSG00000197748 | 67,56042686 | 2,894608221 | 0,280980206 | 10,30182255 | 6,91388E-25 | 1,3173E-22 | CFAP43 | protein_coding |
| ENSG00000198771 | 22,9188964 | 5,277256158 | 0,871750716 | 6,053629853 | 1,41618E-09 | 4,07157E-08 | RCSD1 | protein_coding |
| ENSG00000198959 | 107,8962046 | 3,386032897 | 0,267625653 | 12,65212382 | 1,08891E-36 | 5,81543E-34 | TGM2 | protein_coding |
| ENSG00000203867 | 22,8819153 | 3,271347379 | 0,476663082 | 6,86301814 | 6,74207E-12 | 3,00056E-10 | RBM20 | protein_coding |
| ENSG00000204176 | 35,07021404 | 2,51050708 | 0,364786893 | 6,882119756 | 5,89684E-12 | 2,64107E-10 | SYT15 | protein_coding |
| ENSG00000204653 | 26,26243523 | 3,633971514 | 0,617525101 | 5,884734904 | 3,98693E-09 | 1,05425E-07 | ASPDH | protein_coding |
| ENSG00000205108 | 32,88877491 | 2,382852051 | 0,36119825 | 6,597075286 | 4,19348E-11 | 1,63677E-09 | FAM205A | protein_coding |
| ENSG00000205181 | 18,15922783 | 3,133030233 | 0,55316168 | 5,663859853 | 1,48005E-08 | 3,3942E-07 | LINC00654 | lncRNA |
| ENSG00000205336 | 39,367077 | 7,923189195 | 1,198386172 | 6,611549247 | 3,80319E-11 | 1,50117E-09 | ADGRG1 | protein_coding |
| ENSG00000205517 | 49,01626637 | 2,216051061 | 0,268296644 | 8,25970473 | 1,46033E-16 | 1,1752E-14 | RGL3 | protein_coding |
| ENSG00000205832 | 33,43511939 | 2,231853655 | 0,367653914 | 6,070528756 | 1,2749E-09 | 3,70161E-08 | C16orf96 | protein_coding |
| ENSG00000205838 | 77,77825065 | 2,065922821 | 0,24521406 | 8,424977033 | 3,60821E-17 | 3,06463E-15 | TTC23L | protein_coding |
| ENSG00000206549 | 19,44217915 | 2,45744865 | 0,420152563 | 5,848943614 | 4,94705E-09 | 1,27466E-07 | ENSG00000206549 | protein_coding |
| ENSG00000213937 | 26,47689174 | 2,28291849 | 0,379884673 | 6,009504074 | 1,86092E-09 | 5,20171E-08 | CLDN9 | protein_coding |
| ENSG00000215644 | 20,4788848 | 4,025102957 | 0,670828734 | 6,000194618 | 1,97081E-09 | 5,47417E-08 | GCGR | protein_coding |
| ENSG00000218730 | 46,08102436 | 6,499117837 | 0,674089326 | 9,641330294 | 5,34902E-22 | 7,51165E-20 | ENSG00000218730 | processed_pseudogene |
| ENSG00000222078 | 80,39645476 | 5,690516843 | 0,618220766 | 9,204667907 | 3,42729E-20 | 3,96083E-18 | RN7SKP110 | misc_RNA |
| ENSG00000224514 | 33,00729954 | 2,294087065 | 0,399328395 | 5,744863366 | 9,19952E-09 | 2,21644E-07 | LINC00620 | lncRNA |
| ENSG00000224940 | 146,5391305 | 2,917684712 | 0,208619918 | 13,98564787 | 1,90732E-44 | 1,81701E-41 | PRRT4 | protein_coding |
| ENSG00000225950 | 31,75173152 | 2,573480861 | 0,413446096 | 6,22446526 | 4,83201E-10 | 1,51799E-08 | NTF4 | protein_coding |
| ENSG00000226455 | 57,5752517 | 4,256836848 | 0,452845282 | 9,40020139 | 5,44587E-21 | 6,7829E-19 | ENSG00000226455 | lncRNA |
| ENSG00000226869 | 8,572941445 | 6,953322716 | 1,251334228 | 5,556727019 | 2,7488E-08 | 5,88638E-07 | LHFPL3-AS1 | lncRNA |
| ENSG00000228013 | 13,71514211 | 4,472452832 | 0,802785732 | 5,57116632 | 2,5304E-08 | 5,46181E-07 | IL6R-AS1 | lncRNA |
| ENSG00000228140 | 21,66856158 | 3,322861695 | 0,485046648 | 6,850602329 | 7,35397E-12 | 3,25235E-10 | EFHD2-AS1 | lncRNA |
| ENSG00000229089 | 23,66742858 | 2,382681054 | 0,414982069 | 5,741648205 | 9,37594E-09 | 2,25125E-07 | ANKRD20A8P | transcribed_unprocessed_pseudogene |
| ENSG00000229162 | 30,30952364 | 2,870365382 | 0,389612968 | 7,367222384 | 1,7422E-13 | 9,6707E-12 | RUNX3-AS1 | lncRNA |
| ENSG00000229474 | 91,7226923 | 2,123204946 | 0,245213096 | 8,658611539 | 4,77545E-18 | 4,46486E-16 | PATL2 | protein_coding |
| ENSG00000231301 | 21,90509235 | 3,170080547 | 0,513944794 | 6,168134366 | 6,91004E-10 | 2,10871E-08 | RPL13AP | processed_pseudogene |
| ENSG00000232022 | 28,74561382 | 2,984153235 | 0,460376677 | 6,481981782 | 9,05256E-11 | 3,29258E-09 | FAAHP1 | transcribed_unprocessed_pseudogene |
| ENSG00000232871 | 24,69194789 | 3,739012067 | 0,51540012 | 7,254581289 | 4,02906E-13 | 2,1583E-11 | SEC1P | transcribed_unitary_pseudogene |
| ENSG00000233217 | 18,93548325 | 6,609360087 | 1,064253864 | 6,210322847 | 5,28759E-10 | 1,65228E-08 | MROH3P | transcribed_unitary_pseudogene |
| ENSG00000235151 | 21,6640103 | 2,613909429 | 0,402453537 | 6,494934669 | 8,30695E-11 | 3,05003E-09 | ENSG00000235151 | lncRNA |
| ENSG00000242866 | 31,34185633 | 2,094557801 | 0,338765961 | 6,182905142 | 6,29325E-10 | 1,93059E-08 | STRC | protein_coding |
| ENSG00000243323 | 86,20875275 | 4,488438975 | 0,386709503 | 11,60674598 | 3,80834E-31 | 1,22033E-28 | PTPRVP | unitary_pseudogene |
| ENSG00000244509 | 58,21111876 | 4,502990313 | 0,572599401 | 7,864119845 | 3,71702E-15 | 2,50991E-13 | APOBEC3C | protein_coding |
| ENSG00000246323 | 22,47311337 | 4,253814335 | 0,57150999 | 7,443114578 | 9,83387E-14 | 5,64535E-12 | FAM13B-AS1 | lncRNA |
| ENSG00000248485 | 112,6022887 | 2,637713267 | 0,259105451 | 10,18007632 | 2,4337E-24 | 4,35448E-22 | PCP4L1 | protein_coding |
| ENSG00000248596 | 17,33702581 | 3,82849952 | 0,61668159 | 6,208227362 | 5,35856E-10 | 1,67297E-08 | ENSG00000248596 | lncRNA |
| ENSG00000255346 | 25,71613081 | 3,744713585 | 0,584234548 | 6,409606548 | 1,45896E-10 | 5,06653E-09 | NOX5 | protein_coding |
| ENSG00000255367 | 29,46971315 | 2,043506493 | 0,361131197 | 5,658626304 | 1,52589E-08 | 3,48448E-07 | ENSG00000255367 | lncRNA |
| ENSG00000255974 | 34,91623082 | 2,19715149 | 0,369745242 | 5,94233878 | 2,80984E-09 | 7,61856E-08 | CYP2A6 | protein_coding |
| ENSG00000256235 | 642,0764804 | 3,340880664 | 0,148953 | 22,42909287 | 2,0477E-111 | 1,8044E-107 | SMIM3 | protein_coding |
| ENSG00000257335 | 35,76907282 | 5,877396836 | 1,063382641 | 5,527076153 | 3,25612E-08 | 6,81946E-07 | MGAM | protein_coding |
| ENSG00000257743 | 25,63764666 | 8,042341122 | 1,222685177 | 6,577605806 | 4,78084E-11 | 1,84371E-09 | MGAM2 | protein_coding |
| ENSG00000258811 | 34,24427425 | 2,148021059 | 0,372984803 | 5,759004232 | 8,46116E-09 | 2,05824E-07 | ENSG00000258811 | lncRNA |
| ENSG00000259417 | 88,11571761 | 3,728702463 | 0,257957352 | 14,45472455 | 2,34064E-47 | 2,66139E-44 | CTXND1 | protein_coding |
| ENSG00000260220 | 20,42165955 | 7,147777275 | 1,221161199 | 5,853262681 | 4,82022E-09 | 1,24654E-07 | CCDC187 | protein_coding |
| ENSG00000260401 | 117,6464787 | 3,741282948 | 0,241335248 | 15,50243068 | 3,34013E-54 | 5,60633E-51 | ENSG00000260401 | lncRNA |
| ENSG00000262155 | 39,3924242 | 2,163409509 | 0,2993129 | 7,227919365 | 4,9045E-13 | 2,5957E-11 | LINC02175 | lncRNA |
| ENSG00000262943 | 40,98324488 | 2,95029401 | 0,38648345 | 7,633687827 | 2,28132E-14 | 1,40335E-12 | ALOX12P2 | transcribed_unprocessed_pseudogene |
| ENSG00000264589 | 20,93237171 | 3,514637024 | 0,521549081 | 6,738842335 | 1,59654E-11 | 6,62055E-10 | MAPT-AS1 | lncRNA |
| ENSG00000267226 | 50,87650999 | 4,330072888 | 0,393387133 | 11,00715433 | 3,52977E-28 | 9,35469E-26 | MALT1-AS1 | lncRNA |
| ENSG00000267750 | 38,643259 | 2,373381114 | 0,350652785 | 6,768465032 | 1,30156E-11 | 5,50088E-10 | RUNDC3A-AS1 | lncRNA |
| ENSG00000271447 | 23,20734697 | 3,635510158 | 0,550677972 | 6,601880491 | 4,05975E-11 | 1,59175E-09 | MMP28 | protein_coding |
| ENSG00000275395 | 73,76799311 | 2,520035206 | 0,281746599 | 8,944332304 | 3,74205E-19 | 3,97289E-17 | FCGBP | protein_coding |
| ENSG00000275620 | 16,31415814 | 5,542590334 | 0,887147132 | 6,247656261 | 4,16657E-10 | 1,33391E-08 | ENSG00000275620 | lncRNA |
| ENSG00000277152 | 62,17946397 | 3,19192774 | 0,327886788 | 9,734847087 | 2,1414E-22 | 3,145E-20 | ENSG00000277152 | lncRNA |
| ENSG00000278195 | 15,68168754 | 7,642219623 | 1,23526411 | 6,186709031 | 6,14332E-10 | 1,89449E-08 | SSTR3 | protein_coding |
| ENSG00000279108 | 39,06601539 | 2,037281754 | 0,347754297 | 5,858394193 | 4,67364E-09 | 1,21398E-07 | ENSG00000279108 | TEC |

E10 LTR.SAM_vs_non.act

| ensembl_gene_id | baseMean | log2FoldChange | lfcSE | stat | pvalue | padj | hgnc_symbol | gene_biotype |
|---|---|---|---|---|---|---|---|---|
| ENSG00000167105 | 16,5699056 | 3,05142164 | 0,561817749 | 5,431337202 | 5,59333E-08 | 9,52143E-07 | TMEM92 | protein_coding |
| ENSG00000140519 | 19,8612697 | 2,308395501 | 0,42477098 | 5,434447296 | 5,49665E-08 | 9,37081E-07 | RHCG | protein_coding |
| ENSG00000229162 | 30,30952364 | 2,145013456 | 0,39464533 | 5,435294152 | 5,47061E-08 | 9,33105E-07 | RUNX3-AS1 | lncRNA |
| ENSG00000260220 | 20,42165955 | 6,648535718 | 1,222960126 | 5,436428853 | 5,4359E-08 | 9,28108E-07 | CCDC187 | protein_coding |
| ENSG00000072952 | 17,32678809 | 2,954767301 | 0,543261304 | 5,438943068 | 5,35976E-08 | 9,15563E-07 | IRAG1 | protein_coding |
| ENSG00000145708 | 29,59814114 | -2,175234574 | 0,399017526 | -5,451476271 | 4,99534E-08 | 8,58442E-07 | CRHBP | protein_coding |
| ENSG00000249006 | 20,55744169 | 2,425928515 | 0,442906549 | 5,477292043 | 4,31884E-08 | 7,50459E-07 | ENSG00000249006 | processed_pseudogene |
| ENSG00000153237 | 26,44299374 | 2,144403645 | 0,391100672 | 5,482996574 | 4,18181E-08 | 7,28493E-07 | CCDC148 | protein_coding |
| ENSG00000112539 | 14,60348547 | 3,317000643 | 0,604259403 | 5,489365368 | 4,0338E-08 | 7,05936E-07 | C6orf118 | protein_coding |
| ENSG00000152760 | 11,83620777 | 6,846355588 | 1,241061881 | 5,516530393 | 3,45758E-08 | 6,12595E-07 | DYNLT5 | protein_coding |
| ENSG00000111181 | 21,6110929 | 2,415913161 | 0,437687742 | 5,519718573 | 3,39543E-08 | 6,02206E-07 | SLC6A12 | protein_coding |
| ENSG00000074966 | 16,67694599 | 3,204910722 | 0,579950181 | 5,52618281 | 3,27273E-08 | 5,82553E-07 | TXK | protein_coding |
| ENSG00000215644 | 20,4788848 | 3,722484835 | 0,671166272 | 5,546293055 | 2,9179E-08 | 5,24839E-07 | GCGR | protein_coding |
| ENSG00000080031 | 21,58864789 | 3,335405106 | 0,601288983 | 5,547091662 | 2,9046E-08 | 5,22746E-07 | PTPRH | protein_coding |
| ENSG00000257335 | 35,76907282 | 5,90371626 | 1,061865502 | 5,55975898 | 2,70147E-08 | 4,8873E-07 | MGAM | protein_coding |
| ENSG00000188282 | 12,41113603 | 4,486919045 | 0,804979687 | 5,573953128 | 2,49023E-08 | 4,53623E-07 | RUFY4 | protein_coding |
| ENSG00000123243 | 15,74736214 | 3,240223437 | 0,579326464 | 5,593087209 | 2,23067E-08 | 4,09825E-07 | ITIH5 | protein_coding |
| ENSG00000204176 | 35,07021404 | 2,063911268 | 0,366806944 | 5,626696288 | 1,83694E-08 | 3,42439E-07 | SYT15 | protein_coding |
| ENSG00000240219 | 24,12641831 | 2,273074943 | 0,400217772 | 5,679595219 | 1,35014E-08 | 2,58288E-07 | ENSG00000240219 | lncRNA |
| ENSG00000162692 | 31,61464606 | -2,186781968 | 0,384784854 | -5,683129007 | 1,32252E-08 | 2,53147E-07 | VCAM1 | protein_coding |
| ENSG00000137699 | 10,51172075 | 3,852859881 | 0,677335939 | 5,688255503 | 1,28344E-08 | 2,46215E-07 | TRIM29 | protein_coding |
| ENSG00000141506 | 18,25342339 | 2,876826089 | 0,505579704 | 5,690153431 | 1,26925E-08 | 2,43766E-07 | PIK3R5 | protein_coding |
| ENSG00000175985 | 32,51268074 | 2,240974467 | 0,391029319 | 5,730962757 | 9,98622E-09 | 1,94845E-07 | PLEKHD1 | protein_coding |
| ENSG00000271447 | 23,20734697 | 3,17909176 | 0,55272673 | 5,751651922 | 8,83756E-09 | 1,74417E-07 | MMP28 | protein_coding |
| ENSG00000131831 | 30,39697986 | 2,278353168 | 0,395138281 | 5,76596417 | 8,11923E-09 | 1,61917E-07 | RAI2 | protein_coding |
| ENSG00000260776 | 11,88361601 | 7,155727604 | 1,240053796 | 5,770497723 | 7,90377E-09 | 1,57804E-07 | GOLGA6FP | transcribed_unprocessed_pseudogene |
| ENSG00000104870 | 33,0219089 | 2,574801528 | 0,445905977 | 5,774314908 | 7,72668E-09 | 1,54719E-07 | FCGRT | protein_coding |
| ENSG00000155980 | 35,76339107 | 2,840586109 | 0,490272132 | 5,793896746 | 6,87717E-09 | 1,38896E-07 | KIF5A | protein_coding |
| ENSG00000154783 | 27,56954341 | 2,475854858 | 0,426969882 | 5,798663945 | 6,68453E-09 | 1,35512E-07 | FGD5 | protein_coding |
| ENSG00000267121 | 25,14055091 | 2,223573857 | 0,383080439 | 5,804456798 | 6,4575E-09 | 1,31297E-07 | FMNL1-DT | lncRNA |
| ENSG00000102445 | 28,03047282 | 3,554024171 | 0,61168754 | 5,810195462 | 6,23999E-09 | 1,27101E-07 | RUBCNL | protein_coding |
| ENSG00000187554 | 32,48806765 | 2,016792845 | 0,346689306 | 5,817291763 | 5,98087E-09 | 1,22186E-07 | TLR5 | protein_coding |
| ENSG00000003400 | 38,17543049 | 2,186048532 | 0,375690583 | 5,818747213 | 5,92903E-09 | 1,21199E-07 | CASP10 | protein_coding |
| ENSG00000200818 | 19,18190101 | -2,447862845 | 0,420251146 | -5,824761849 | 5,71941E-09 | 1,17263E-07 | RNU6-1204P | snRNA |
| ENSG00000142606 | 24,77043762 | 2,800158511 | 0,480271172 | 5,830369745 | 5,53047E-09 | 1,13525E-07 | MMEL1 | protein_coding |
| ENSG00000205832 | 33,43511939 | 2,136078662 | 0,365475783 | 5,844651717 | 5,07629E-09 | 1,04956E-07 | C16orf96 | protein_coding |
| ENSG00000198846 | 62,51102885 | -2,131027594 | 0,36310069 | -5,868971481 | 4,38507E-09 | 9,1436E-08 | TOX | protein_coding |
| ENSG00000159208 | 27,57291636 | 2,678047587 | 0,456020258 | 5,87265048 | 4,28882E-09 | 8,97017E-08 | CIART | protein_coding |
| ENSG00000164342 | 24,19455123 | 2,448036172 | 0,41648268 | 5,877882298 | 4,15548E-09 | 8,71787E-08 | TLR3 | protein_coding |
| ENSG00000198771 | 22,9188964 | 5,133466346 | 0,870894063 | 5,894478518 | 3,75866E-09 | 7,92903E-08 | RCSD1 | protein_coding |
| ENSG00000257743 | 25,63764666 | 7,235305628 | 1,225256094 | 5,905137434 | 3,52353E-09 | 7,46975E-08 | MGAM2 | protein_coding |
| ENSG00000187688 | 35,83748957 | 2,090544382 | 0,353616802 | 5,91189211 | 3,382E-09 | 7,18748E-08 | TRPV2 | protein_coding |
| ENSG00000107859 | 20,53411021 | 3,195993551 | 0,53912272 | 5,928137387 | 3,0639E-09 | 6,57254E-08 | PITX3 | protein_coding |
| ENSG00000248596 | 17,33702581 | 3,650101571 | 0,615624398 | 5,929104795 | 3,04591E-09 | 6,53803E-08 | ENSG00000248596 | lncRNA |
| ENSG00000160808 | 19,47482869 | 3,620305935 | 0,610181725 | 5,933160216 | 2,97159E-09 | 6,3945E-08 | MYL3 | protein_coding |
| ENSG00000122176 | 14,38398387 | 4,094687547 | 0,689641463 | 5,93741497 | 2,89551E-09 | 6,23863E-08 | FMOD | protein_coding |
| ENSG00000158825 | 29,59030788 | 2,569064319 | 0,429921985 | 5,97565235 | 2,29171E-09 | 5,03575E-08 | CDA | protein_coding |
| ENSG00000142910 | 17,9936797 | 3,234368991 | 0,540854041 | 5,980114311 | 2,22981E-09 | 4,91231E-08 | TINAGL1 | protein_coding |
| ENSG00000118292 | 19,7771347 | 3,662715016 | 0,611423038 | 5,990475972 | 2,09228E-09 | 4,63613E-08 | C1orf54 | protein_coding |
| ENSG00000183876 | 18,97958051 | 3,181173314 | 0,526726621 | 6,03951497 | 1,54578E-09 | 3,5021E-08 | ARSI | protein_coding |
| ENSG00000225950 | 31,75173152 | 2,486946931 | 0,411469186 | 6,044066032 | 1,50278E-09 | 3,40918E-08 | NTF4 | protein_coding |
| ENSG00000128040 | 30,1991571 | 2,226160489 | 0,368114199 | 6,047472488 | 1,47136E-09 | 3,34454E-08 | SPINK2 | protein_coding |
| ENSG00000179930 | 20,99924027 | 4,671474148 | 0,771378328 | 6,056009069 | 1,3954E-09 | 3,18454E-08 | ZNF648 | protein_coding |
| ENSG00000153253 | 30,5396611 | 2,447278575 | 0,403251735 | 6,068860627 | 1,28821E-09 | 2,95367E-08 | SCN3A | protein_coding |
| ENSG00000224514 | 33,00729954 | 2,403601531 | 0,395526937 | 6,076960397 | 1,22482E-09 | 2,81208E-08 | LINC00620 | lncRNA |
| ENSG00000105357 | 53,33016608 | 6,320839759 | 1,037444198 | 6,092703365 | 1,1102E-09 | 2,56956E-08 | MYH14 | protein_coding |
| ENSG00000226435 | 27,62106964 | 2,37403373 | 0,389187697 | 6,099971162 | 1,06088E-09 | 2,46039E-08 | ANKRD18DP | transcribed_unprocessed_pseudogene |
| ENSG00000205181 | 18,15922783 | 3,345558989 | 0,547176579 | 6,114221836 | 9,70293E-10 | 2,26254E-08 | LINC00654 | lncRNA |
| ENSG00000174844 | 32,17621975 | 2,533471207 | 0,413137599 | 6,132269771 | 8,6634E-10 | 2,03956E-08 | DNAH12 | protein_coding |
| ENSG00000172005 | 30,09470292 | 3,423594163 | 0,557682858 | 6,138962514 | 8,30622E-10 | 1,96491E-08 | MAL | protein_coding |
| ENSG00000250423 | 62,20598494 | 3,535176284 | 0,575606692 | 6,141652513 | 8,16674E-10 | 1,93325E-08 | KIAA1210 | protein_coding |
| ENSG00000179630 | 24,25067493 | 2,875149307 | 0,468061933 | 6,142668537 | 8,11465E-10 | 1,92224E-08 | LACC1 | protein_coding |
| ENSG00000177108 | 49,09942928 | 2,22888631 | 0,362644047 | 6,146209563 | 7,93564E-10 | 1,88374E-08 | ZDHHC22 | protein_coding |
| ENSG00000157423 | 33,53042305 | 2,629069551 | 0,426194857 | 6,168703134 | 6,88524E-10 | 1,65617E-08 | HYDIN | protein_coding |
| ENSG00000183287 | 37,4782 | 2,25986459 | 0,366058944 | 6,173499172 | 6,67949E-10 | 1,61082E-08 | CCBE1 | protein_coding |
| ENSG00000174348 | 31,01562375 | 3,03586635 | 0,488937114 | 6,209114142 | 5,32841E-10 | 1,30657E-08 | PODN | protein_coding |
| ENSG00000196337 | 29,55256542 | 2,308557256 | 0,370924758 | 6,223788541 | 4,85291E-10 | 1,19412E-08 | CGB7 | protein_coding |
| ENSG00000163637 | 26,71897658 | 2,848918894 | 0,456782315 | 6,236929055 | 4,46244E-10 | 1,1026E-08 | PRICKLE2 | protein_coding |
| ENSG00000089163 | 29,83367976 | 2,256434927 | 0,361137145 | 6,248138581 | 4,15373E-10 | 1,03244E-08 | SIRT4 | protein_coding |
| ENSG00000163239 | 40,12742841 | 2,318318971 | 0,370345744 | 6,259877442 | 3,8528E-10 | 9,61826E-09 | TDRD10 | protein_coding |
| ENSG00000130635 | 16,67456414 | 4,174641189 | 0,665164044 | 6,276107714 | 3,47154E-10 | 8,74282E-09 | COL5A1 | protein_coding |
| ENSG00000232871 | 24,69194789 | 3,25068531 | 0,517590818 | 6,280415327 | 3,3767E-10 | 8,53531E-09 | SEC1P | transcribed_unitary_pseudogene |
| ENSG00000142408 | 46,81587608 | 2,078603726 | 0,330494686 | 6,289371101 | 3,18755E-10 | 8,09297E-09 | CACNG8 | protein_coding |
| ENSG00000229089 | 23,66742858 | 2,576836226 | 0,409033399 | 6,299818631 | 2,97994E-10 | 7,59398E-09 | ANKRD20A8P | transcribed_unprocessed_pseudogene |
| ENSG00000188672 | 32,22698711 | 2,358844349 | 0,374366831 | 6,300890332 | 2,95941E-10 | 7,55287E-09 | RHCE | protein_coding |
| ENSG00000169064 | 23,59455979 | 2,836400718 | 0,449979143 | 6,303404858 | 2,91177E-10 | 7,44792E-09 | ZBBX | protein_coding |
| ENSG00000205108 | 32,88877491 | 2,265385342 | 0,359149372 | 6,307641113 | 2,8332E-10 | 7,2632E-09 | FAM205A | protein_coding |
| ENSG00000177692 | 29,7365375 | 2,572899243 | 0,406838963 | 6,324122022 | 2,54676E-10 | 6,59295E-09 | DNAJC28 | protein_coding |
| ENSG00000204653 | 26,26243523 | 3,876316782 | 0,612519535 | 6,328478623 | 2,4759E-10 | 6,42405E-09 | ASPDH | protein_coding |
| ENSG00000154262 | 18,28891883 | 3,275074675 | 0,516690676 | 6,338559665 | 2,31923E-10 | 6,03117E-09 | ABCA6 | protein_coding |
| ENSG00000258711 | 38,85073234 | -2,517145869 | 0,393113474 | -6,403102498 | 1,52251E-10 | 4,08316E-09 | ENSG00000258711 | lncRNA |
| ENSG00000042832 | 19,46992073 | 3,587572472 | 0,55983762 | 6,408237569 | 1,47211E-10 | 3,95729E-09 | TG | protein_coding |
| ENSG00000013016 | 46,74533462 | 3,101474137 | 0,483739558 | 6,411454435 | 1,44138E-10 | 3,8838E-09 | EHD3 | protein_coding |
| ENSG00000206549 | 19,44217915 | 2,659672145 | 0,414497876 | 6,416612241 | 1,3934E-10 | 3,76434E-09 | ENSG00000206549 | protein_coding |
| ENSG00000143851 | 51,14985771 | 2,122691485 | 0,330610953 | 6,420511683 | 1,35817E-10 | 3,67402E-09 | PTPN7 | protein_coding |
| ENSG00000215910 | 36,05932083 | 2,498271087 | 0,387446998 | 6,448033157 | 1,13311E-10 | 3,08957E-09 | C1orf167 | protein_coding |
| ENSG00000019505 | 29,27880986 | 5,627970038 | 0,872441931 | 6,450824789 | 1,11243E-10 | 3,04044E-09 | SYT13 | protein_coding |

| | | | | | | | | |
|---|---|---|---|---|---|---|---|---|
| ENSG00000178031 | 156,5166925 | -2,30641963 | 0,356432444 | -6,470846489 | 9,74554E-11 | 2,69582E-09 | ADAMTSL1 | protein_coding |
| ENSG00000137474 | 33,00823762 | 5,682035206 | 0,878008357 | 6,471504699 | 9,70317E-11 | 2,68844E-09 | MYO7A | protein_coding |
| ENSG00000132031 | 24,177042 | 2,776869864 | 0,427939423 | 6,488932112 | 8,64469E-11 | 2,40683E-09 | MATN3 | protein_coding |
| ENSG00000142677 | 24,77013325 | 3,649354879 | 0,555325753 | 6,571557067 | 4,97918E-11 | 1,44493E-09 | IL22RA1 | protein_coding |
| ENSG00000188783 | 19,84793134 | 3,431846448 | 0,521756146 | 6,577491179 | 4,78452E-11 | 1,3908E-09 | PRELP | protein_coding |
| ENSG00000145794 | 42,54085402 | 2,608152973 | 0,394501135 | 6,611268623 | 3,8104E-11 | 1,13455E-09 | MEGF10 | protein_coding |
| ENSG00000173210 | 50,74359587 | 2,195734189 | 0,33202277 | 6,613203635 | 3,76091E-11 | 1,12079E-09 | ABLIM3 | protein_coding |
| ENSG00000174403 | 52,80696112 | 2,120065638 | 0,320548618 | 6,613866093 | 3,7441E-11 | 1,11675E-09 | MIR1-1HG-AS1 | lncRNA |
| ENSG00000105639 | 32,80124725 | 3,792212082 | 0,572849163 | 6,619913802 | 3,59408E-11 | 1,07294E-09 | JAK3 | protein_coding |
| ENSG00000075673 | 44,15268178 | 2,530963501 | 0,380916712 | 6,644401315 | 3,04452E-11 | 9,20623E-10 | ATP12A | protein_coding |
| ENSG00000229739 | 45,05494248 | 2,158228046 | 0,319918885 | 6,746172697 | 1,51796E-11 | 4,77684E-10 | PDC-AS1 | lncRNA |
| ENSG00000232022 | 28,74561382 | 3,089777989 | 0,456065476 | 6,774856142 | 1,2453E-11 | 3,95876E-10 | FAAHP1 | transcribed_unprocessed_pseudogene |
| ENSG00000134668 | 29,04793169 | 2,633077506 | 0,386128064 | 6,819181895 | 9,15604E-12 | 2,94893E-10 | SPOCD1 | protein_coding |
| ENSG00000128052 | 19,90832369 | 3,72599596 | 0,545385178 | 6,831861427 | 8,38198E-12 | 2,72189E-10 | KDR | protein_coding |
| ENSG00000214456 | 41,97462221 | 2,276393454 | 0,332734201 | 6,84147721 | 7,83807E-12 | 2,55564E-10 | PLIN5 | protein_coding |
| ENSG00000267750 | 38,643259 | 2,377843084 | 0,347270488 | 6,847236289 | 7,52903E-12 | 2,45956E-10 | RUNDC3A-AS1 | lncRNA |
| ENSG00000118640 | 22,29578137 | 3,348281509 | 0,488915114 | 6,848390269 | 7,46856E-12 | 2,44213E-10 | VAMP8 | protein_coding |
| ENSG00000229015 | 35,85310318 | -2,457735915 | 0,356718377 | -6,889849455 | 5,58515E-12 | 1,85277E-10 | ENSG00000229015 | transcribed_unprocessed_pseudogene |
| ENSG00000159708 | 30,24040437 | 2,861135037 | 0,413866219 | 6,913188144 | 4,73881E-12 | 1,58427E-10 | LRRC36 | protein_coding |
| ENSG00000087245 | 30,5706812 | 3,609154426 | 0,520719629 | 6,931089644 | 4,17611E-12 | 1,4085E-10 | MMP2 | protein_coding |
| ENSG00000158683 | 62,092147 | 2,074747056 | 0,299173823 | 6,934921766 | 4,06446E-12 | 1,37355E-10 | PKD1L1 | protein_coding |
| ENSG00000277152 | 62,17946397 | 2,328145934 | 0,331878581 | 7,015053299 | 2,29861E-12 | 7,98789E-11 | ENSG00000277152 | lncRNA |
| ENSG00000184497 | 46,24431071 | 2,174648442 | 0,308477826 | 7,049610251 | 1,7942E-12 | 6,31856E-11 | TMEM255B | protein_coding |
| ENSG00000139567 | 43,82437808 | 6,136665628 | 0,867134407 | 7,076948597 | 1,47363E-12 | 5,27631E-11 | ACVRL1 | protein_coding |
| ENSG00000139973 | 32,67936719 | 3,630305908 | 0,511917692 | 7,091581247 | 1,32588E-12 | 4,80751E-11 | SYT16 | protein_coding |
| ENSG00000084636 | 35,074958 | 4,216819058 | 0,59310313 | 7,109756883 | 1,16248E-12 | 4,24644E-11 | COL16A1 | protein_coding |
| ENSG00000189120 | 32,01089699 | 4,041323034 | 0,568401858 | 7,10997506 | 1,16064E-12 | 4,24425E-11 | SP6 | protein_coding |
| ENSG00000061337 | 59,85536919 | 2,012587946 | 0,282797878 | 7,116701019 | 1,10541E-12 | 4,06831E-11 | LZTS1 | protein_coding |
| ENSG00000141505 | 65,37748526 | 2,003898209 | 0,280775017 | 7,13702463 | 9,53728E-13 | 3,54429E-11 | ASGR1 | protein_coding |
| ENSG00000178662 | 120,6374984 | 2,274315823 | 0,316595645 | 7,183661106 | 6,7869E-13 | 2,56951E-11 | CSRNP3 | protein_coding |
| ENSG00000141485 | 62,41351389 | 2,885228186 | 0,399894408 | 7,214975072 | 5,39439E-13 | 2,07902E-11 | SLC13A5 | protein_coding |
| ENSG00000205336 | 39,367077 | 8,643357402 | 1,195233956 | 7,231519286 | 4,7762E-13 | 1,85536E-11 | ADGRG1 | protein_coding |
| ENSG00000157103 | 37,79162273 | 3,085147316 | 0,426342255 | 7,236316079 | 4,61036E-13 | 1,79681E-11 | SLC6A1 | protein_coding |
| ENSG00000197177 | 46,26780958 | 2,832246733 | 0,390472211 | 7,253388734 | 4,06471E-13 | 1,5916E-11 | ADGRA1 | protein_coding |
| ENSG00000118514 | 50,0537567 | 2,332366495 | 0,319588064 | 7,298040053 | 2,9199E-13 | 1,1619E-11 | ALDH8A1 | protein_coding |
| ENSG00000036530 | 55,10944722 | 2,399103953 | 0,326364736 | 7,350990124 | 1,96744E-13 | 8,01486E-12 | CYP46A1 | protein_coding |
| ENSG00000023445 | 41,60082792 | 2,675423053 | 0,363755797 | 7,354997699 | 1,9093E-13 | 7,79655E-12 | BIRC3 | protein_coding |
| ENSG00000255346 | 25,71613081 | 4,246980377 | 0,57741436 | 7,35516931 | 1,90685E-13 | 7,79582E-12 | NOX5 | protein_coding |
| ENSG00000090530 | 30,38554495 | 2,707073656 | 0,366867784 | 7,378880829 | 1,59626E-13 | 6,60473E-12 | P3H2 | protein_coding |
| ENSG00000121101 | 59,48247272 | 2,12178204 | 0,284865959 | 7,448352356 | 9,45131E-14 | 4,01225E-12 | TEX14 | protein_coding |
| ENSG00000160111 | 47,09603347 | 3,184110344 | 0,419372427 | 7,592560078 | 3,13646E-14 | 1,39719E-12 | CPAMD8 | protein_coding |
| ENSG00000166546 | 78,01003981 | 2,014608618 | 0,264120507 | 7,627611502 | 2,39143E-14 | 1,07649E-12 | BEAN1 | protein_coding |
| ENSG00000163075 | 48,0094885 | 2,574812736 | 0,337350599 | 7,632453438 | 2,30328E-14 | 1,03954E-12 | CFAP221 | protein_coding |
| ENSG00000142233 | 39,12327797 | 4,006043343 | 0,522300358 | 7,669999231 | 1,71997E-14 | 7,87675E-13 | NTN5 | protein_coding |
| ENSG00000263528 | 56,46283572 | 2,492935332 | 0,324464016 | 7,683241323 | 1,55113E-14 | 7,12253E-13 | IKBKE | protein_coding |
| ENSG00000244509 | 58,21111876 | 4,393357226 | 0,571694319 | 7,684801261 | 1,53235E-14 | 7,05517E-13 | APOBEC3C | protein_coding |
| ENSG00000115596 | 42,17552152 | 3,591400354 | 0,465872164 | 7,708982485 | 1,26825E-14 | 5,87867E-13 | WNT6 | protein_coding |
| ENSG00000006283 | 99,62152007 | 2,052991387 | 0,265801707 | 7,723770507 | 1,12938E-14 | 5,27778E-13 | CACNA1G | protein_coding |
| ENSG00000154263 | 43,91788027 | 2,176112449 | 0,281102497 | 7,741348696 | 9,83677E-15 | 4,62207E-13 | ABCA10 | protein_coding |
| ENSG00000136449 | 87,9712079 | 2,069204587 | 0,265157945 | 7,803668064 | 6,01332E-15 | 2,88884E-13 | MYCBPAP | protein_coding |
| ENSG00000172782 | 67,74159269 | 2,066561662 | 0,264472388 | 7,813903293 | 5,54436E-15 | 2,67855E-13 | FADS6 | protein_coding |
| ENSG00000262943 | 40,98324488 | 3,000035329 | 0,383640936 | 7,819904091 | 5,28636E-15 | 2,5867E-13 | ALOX12P2 | transcribed_unprocessed_pseudogene |
| ENSG00000073861 | 48,81304673 | 3,186451202 | 0,406683726 | 7,835207059 | 4,68071E-15 | 2,31678E-13 | TBX21 | protein_coding |
| ENSG00000174460 | 258,9302181 | 2,274716395 | 0,28823912 | 7,89176845 | 2,97934E-15 | 1,50507E-13 | ZCCHC12 | protein_coding |
| ENSG00000184454 | 33,60086497 | 3,8885764 | 0,492123414 | 7,901628509 | 2,75283E-15 | 1,39889E-13 | NCMAP | protein_coding |
| ENSG00000146005 | 53,5039062 | 2,333193181 | 0,295018561 | 7,908631819 | 2,60233E-15 | 1,32437E-13 | PSD2 | protein_coding |
| ENSG00000163898 | 58,26773528 | 2,141459533 | 0,270717716 | 7,910304361 | 2,5676E-15 | 1,30864E-13 | LIPH | protein_coding |
| ENSG00000170703 | 37,55692612 | 2,847294868 | 0,359933514 | 7,910613375 | 2,56124E-15 | 1,30734E-13 | TTLL6 | protein_coding |
| ENSG00000105122 | 56,06307414 | 2,954920951 | 0,372939427 | 7,92332679 | 2,31239E-15 | 1,18561E-13 | RASAL3 | protein_coding |
| ENSG00000115828 | 65,09475549 | 3,448507315 | 0,433158271 | 7,961310096 | 1,70227E-15 | 8,83859E-14 | QPCT | protein_coding |
| ENSG00000161082 | 65,2355563 | 2,118344933 | 0,265904405 | 7,966565766 | 1,63145E-15 | 8,54355E-14 | CELF5 | protein_coding |
| ENSG00000136931 | 70,29593293 | 2,59263914 | 0,323430767 | 8,016055993 | 1,09195E-15 | 5,83409E-14 | NR5A1 | protein_coding |
| ENSG00000175264 | 31,28823814 | 4,376628374 | 0,544840302 | 8,032864598 | 9,52227E-16 | 5,11148E-14 | CHST1 | protein_coding |
| ENSG00000175793 | 42,4702083 | 2,348392001 | 0,292054197 | 8,040945911 | 8,91476E-16 | 4,82311E-14 | SFN | protein_coding |
| ENSG00000118160 | 44,13915242 | 3,908683984 | 0,485094291 | 8,057575726 | 7,78224E-16 | 4,24386E-14 | SLC8A2 | protein_coding |
| ENSG00000119922 | 62,44794168 | 2,575103514 | 0,31858024 | 8,083061008 | 6,3161E-16 | 3,46638E-14 | IFIT2 | protein_coding |
| ENSG00000175170 | 90,79893133 | 4,378174196 | 0,541085588 | 8,091463338 | 5,89521E-16 | 3,25099E-14 | FAM182B | lncRNA |
| ENSG00000117971 | 62,58754707 | 2,298642488 | 0,28336857 | 8,111847001 | 4,98561E-16 | 2,77166E-14 | CHRNB4 | protein_coding |
| ENSG00000205413 | 53,57885864 | 2,426725106 | 0,296809104 | 8,176046724 | 2,93308E-16 | 1,66569E-14 | SAMD9 | protein_coding |
| ENSG00000042493 | 56,3573933 | 2,493501655 | 0,303811505 | 8,207397073 | 2,26035E-16 | 1,2987E-14 | CAPG | protein_coding |
| ENSG00000116299 | 49,06309292 | 2,464356645 | 0,298745904 | 8,249005644 | 1,59718E-16 | 9,2624E-15 | ELAPOR1 | protein_coding |
| ENSG00000027644 | 80,42153086 | 2,427474037 | 0,293058553 | 8,283239009 | 1,1987E-16 | 6,99259E-15 | INSRR | protein_coding |
| ENSG00000188162 | 80,73670584 | 5,211160142 | 0,624576291 | 8,343512578 | 7,21158E-17 | 4,30199E-15 | OTOG | protein_coding |
| ENSG00000175463 | 132,4470049 | 2,029254217 | 0,241673832 | 8,396665062 | 4,59338E-17 | 2,76902E-15 | TBC1D10C | protein_coding |
| ENSG00000129910 | 32,90903158 | 5,05385563 | 0,597932831 | 8,452212973 | 2,85833E-17 | 1,74455E-15 | CDH15 | protein_coding |
| ENSG00000128340 | 38,41452422 | 2,971875477 | 0,350361145 | 8,482320382 | 2,20747E-17 | 1,35696E-15 | RAC2 | protein_coding |
| ENSG00000108370 | 44,34048549 | 5,373654525 | 0,629951826 | 8,530262644 | 1,46016E-17 | 9,13958E-16 | RGS9 | protein_coding |
| ENSG00000108176 | 59,84432933 | 2,823835656 | 0,327617327 | 8,619311085 | 6,73581E-18 | 4,36758E-16 | DNAJC12 | protein_coding |
| ENSG00000104848 | 63,94631845 | 2,569859534 | 0,297204573 | 8,646769813 | 5,29777E-18 | 3,47455E-16 | KCNA7 | protein_coding |
| ENSG00000186510 | 37,96503529 | 4,026114825 | 0,465497822 | 8,649051903 | 5,1929E-18 | 3,41249E-16 | CLCNKA | protein_coding |
| ENSG00000109339 | 32,47301188 | 3,582111902 | 0,412714612 | 8,679391993 | 3,9789E-18 | 2,66043E-16 | MAPK10 | protein_coding |
| ENSG00000157087 | 55,70403505 | 3,040025041 | 0,349050416 | 8,709415318 | 3,05446E-18 | 2,08292E-16 | ATP2B2 | protein_coding |
| ENSG00000158859 | 106,1566088 | 2,216733428 | 0,253842512 | 8,732711506 | 2,48639E-18 | 1,70571E-16 | ADAMTS4 | protein_coding |
| ENSG00000141837 | 71,50100902 | 5,150190721 | 0,587370409 | 8,768216177 | 1,81519E-18 | 1,2553E-16 | CACNA1A | protein_coding |
| ENSG00000238230 | 92,84575109 | -2,038251617 | 0,229206673 | -8,892636452 | 5,96758E-19 | 4,2823E-17 | LINC00391 | lncRNA |
| ENSG00000205795 | 71,71236047 | 2,03222982 | 0,228521424 | 8,892950974 | 5,95071E-19 | 4,27914E-17 | CYS1 | protein_coding |
| ENSG00000110169 | 75,48935649 | 2,243991299 | 0,250835067 | 8,946082871 | 3,68321E-19 | 2,68803E-17 | HPX | protein_coding |
| ENSG00000229474 | 91,7226923 | 2,191501898 | 0,242887139 | 9,022716927 | 1,83481E-19 | 1,38321E-17 | PATL2 | protein_coding |
| ENSG00000122733 | 60,94914948 | 2,896918763 | 0,319097649 | 9,078471042 | 1,10111E-19 | 8,4684E-18 | PHF24 | protein_coding |

| | | | | | | | | |
|---|---|---|---|---|---|---|---|---|
| ENSG00000103196 | 47,2828225 | 3,634050896 | 0,399446526 | 9,097715621 | 9,22517E-20 | 7,14295E-18 | CRISPLD2 | protein_coding |
| ENSG00000141314 | 83,81559369 | 2,381454263 | 0,260909195 | 9,127521408 | 7,00855E-20 | 5,51376E-18 | RHBDL3 | protein_coding |
| ENSG00000100346 | 58,1955877 | 2,746432243 | 0,300036384 | 9,153663978 | 5,50346E-20 | 4,37991E-18 | CACNA1I | protein_coding |
| ENSG00000184908 | 67,12193238 | 2,455078391 | 0,268060027 | 9,158688904 | 5,25316E-20 | 4,21988E-18 | CLCNKB | protein_coding |
| ENSG00000177301 | 72,10759455 | 3,301459697 | 0,355989768 | 9,274029746 | 1,79243E-20 | 1,52183E-18 | KCNA2 | protein_coding |
| ENSG00000127084 | 69,7869279 | 2,303190383 | 0,247352174 | 9,311381184 | 1,26181E-20 | 1,07933E-18 | FGD3 | protein_coding |
| ENSG00000101204 | 61,10926766 | 3,680833138 | 0,394994193 | 9,318701899 | 1,17772E-20 | 1,015E-18 | CHRNA4 | protein_coding |
| ENSG00000163235 | 64,59063083 | 3,1174664 | 0,333268121 | 9,35422924 | 8,42115E-21 | 7,34997E-19 | TGFA | protein_coding |
| ENSG00000123453 | 57,41692086 | 4,510769368 | 0,480543688 | 9,386803912 | 6,18485E-21 | 5,43966E-19 | SARDH | protein_coding |
| ENSG00000260401 | 117,6464787 | 2,344138508 | 0,247588632 | 9,467876154 | 2,85591E-21 | 2,58471E-19 | ENSG00000260401 | lncRNA |
| ENSG00000160539 | 69,62110148 | 3,343389441 | 0,351386951 | 9,514836653 | 1,81999E-21 | 1,68268E-19 | PLPP7 | protein_coding |
| ENSG00000183508 | 71,91176717 | 3,056928579 | 0,318052399 | 9,611399232 | 7,15697E-22 | 6,78153E-20 | TENT5C | protein_coding |
| ENSG00000169129 | 58,87727685 | 2,307683002 | 0,239093304 | 9,651809391 | 4,82959E-22 | 4,64033E-20 | AFAP1L2 | protein_coding |
| ENSG00000076641 | 119,3526687 | 2,159381515 | 0,223676631 | 9,654032719 | 4,72598E-22 | 4,55354E-20 | PAG1 | protein_coding |
| ENSG00000184584 | 98,85294963 | 2,315996469 | 0,239517367 | 9,669430243 | 4,06636E-22 | 3,94011E-20 | STING1 | protein_coding |
| ENSG00000205517 | 49,01626637 | 2,549030129 | 0,263342878 | 9,679510418 | 3,68476E-22 | 3,61117E-20 | RGL3 | protein_coding |
| ENSG00000135253 | 52,53387293 | 3,628811857 | 0,374293643 | 9,695093468 | 3,16348E-22 | 3,15437E-20 | KCP | protein_coding |
| ENSG00000182040 | 63,42443178 | 3,230019295 | 0,333048161 | 9,698354999 | 3,06399E-22 | 3,06408E-20 | USH1G | protein_coding |
| ENSG00000063127 | 104,0879335 | 2,54890574 | 0,260132968 | 9,798472517 | 1,14301E-22 | 1,17034E-20 | SLC6A16 | protein_coding |
| ENSG00000168477 | 126,1223942 | 2,347965363 | 0,238473553 | 9,84581028 | 7,14611E-23 | 7,47313E-21 | TNXB | protein_coding |
| ENSG00000062038 | 123,7972813 | 2,103502487 | 0,213404981 | 9,856857488 | 6,40218E-23 | 6,75696E-21 | CDH3 | protein_coding |
| ENSG00000115423 | 52,50804759 | 3,102894842 | 0,310754538 | 9,985034687 | 1,77239E-23 | 1,94233E-21 | DNAH6 | protein_coding |
| ENSG00000075651 | 61,36425676 | 2,682893992 | 0,266343924 | 10,07304374 | 7,26929E-24 | 8,25643E-22 | PLD1 | protein_coding |
| ENSG00000010310 | 139,6725342 | 2,047699798 | 0,200814744 | 10,19695943 | 2,04577E-24 | 2,45356E-22 | GIPR | protein_coding |
| ENSG00000148344 | 95,33202855 | 2,274173045 | 0,221916395 | 10,24788208 | 1,20965E-24 | 1,47135E-22 | PTGES | protein_coding |
| ENSG00000146216 | 135,7642231 | 2,130170321 | 0,207806167 | 10,25075603 | 1,17421E-24 | 1,43334E-22 | TTBK1 | protein_coding |
| ENSG00000110324 | 94,82902807 | 2,108118091 | 0,205046013 | 10,28119521 | 8,56587E-25 | 1,0569E-22 | IL10RA | protein_coding |
| ENSG00000149599 | 145,4656593 | 2,049862656 | 0,19901013 | 10,30029304 | 7,0247E-25 | 8,76197E-23 | DUSP15 | protein_coding |
| ENSG00000258776 | 162,1454258 | -2,112017907 | 0,20475864 | -10,3146705 | 6,04884E-25 | 7,62799E-23 | ENSG00000258776 | lncRNA |
| ENSG00000108846 | 62,03154257 | 2,997976157 | 0,290089836 | 10,33464734 | 4,91216E-25 | 6,24044E-23 | ABCC3 | protein_coding |
| ENSG00000185513 | 80,55294616 | 3,022677991 | 0,292193965 | 10,34476531 | 4,41998E-25 | 5,67827E-23 | L3MBTL1 | protein_coding |
| ENSG00000131409 | 93,08016815 | 2,211547836 | 0,212870168 | 10,38918634 | 2,77712E-25 | 3,62198E-23 | LRRC4B | protein_coding |
| ENSG00000188596 | 88,37659924 | 2,989369638 | 0,286157144 | 10,44660145 | 1,5187E-25 | 2,03488E-23 | CFAP54 | protein_coding |
| ENSG00000028137 | 104,3543289 | 2,171457006 | 0,206955663 | 10,4923778 | 9,36417E-26 | 1,27968E-23 | TNFRSF1B | protein_coding |
| ENSG00000162745 | 129,9929893 | 2,274184197 | 0,216496065 | 10,5045059 | 8,23529E-26 | 1,13027E-23 | OLFML2B | protein_coding |
| ENSG00000159307 | 139,8992222 | 2,064467058 | 0,193792288 | 10,6529887 | 1,68855E-26 | 2,49651E-24 | SCUBE1 | protein_coding |
| ENSG00000243323 | 86,20875275 | 4,126911333 | 0,387117265 | 10,66062329 | 1,55551E-26 | 2,31981E-24 | PTPRVP | unitary_pseudogene |
| ENSG00000054179 | 78,94974214 | 2,868752134 | 0,266408306 | 10,76825334 | 4,86136E-27 | 7,64906E-25 | ENTPD2 | protein_coding |
| ENSG00000134202 | 133,3099966 | 2,361082266 | 0,218872765 | 10,78746486 | 3,94523E-27 | 6,26506E-25 | GSTM3 | protein_coding |
| ENSG00000197748 | 67,56042686 | 3,008532314 | 0,278376884 | 10,80740712 | 3,17518E-27 | 5,08934E-25 | CFAP43 | protein_coding |
| ENSG00000184988 | 129,3448816 | 2,053583328 | 0,189414581 | 10,84173834 | 2,18283E-27 | 3,5654E-25 | TMEM106A | protein_coding |
| ENSG00000116981 | 66,65257918 | 2,917527792 | 0,261657391 | 11,150183 | 7,14593E-29 | 1,27001E-26 | NT5C1A | protein_coding |
| ENSG00000224511 | 155,5899012 | -2,124868248 | 0,190522683 | -11,15283604 | 6,93598E-29 | 1,23912E-26 | LINC00365 | lncRNA |
| ENSG00000187800 | 90,42599617 | 3,215107194 | 0,287288139 | 11,19122845 | 4,50153E-29 | 8,08414E-27 | PEAR1 | protein_coding |
| ENSG00000143195 | 87,79582677 | 4,058208833 | 0,360497252 | 11,25725316 | 2,13304E-29 | 3,99811E-27 | ILDR2 | protein_coding |
| ENSG00000275395 | 73,76799311 | 3,117960185 | 0,276320043 | 11,28387268 | 1,5765E-29 | 3,03794E-27 | FCGBP | protein_coding |
| ENSG00000196169 | 81,5569693 | 3,515211555 | 0,310141353 | 11,33422397 | 8,88158E-30 | 1,76097E-27 | KIF19 | protein_coding |
| ENSG00000088340 | 342,2642788 | 2,144194724 | 0,187139294 | 11,45774724 | 2,15035E-30 | 4,49752E-28 | FER1L4 | transcribed_unitary_pseudogene |
| ENSG00000108387 | 149,3357064 | 2,346521464 | 0,204663963 | 11,46524005 | 1,97212E-30 | 4,17566E-28 | SEPTIN4 | protein_coding |
| ENSG00000167207 | 87,68723697 | 2,520513097 | 0,219146861 | 11,50147937 | 1,29674E-30 | 2,81515E-28 | NOD2 | protein_coding |
| ENSG00000169291 | 128,2900061 | 2,301029214 | 0,199808718 | 11,51616022 | 1,09377E-30 | 2,42049E-28 | SHE | protein_coding |
| ENSG00000104967 | 260,3824365 | 2,040583623 | 0,17702903 | 11,52683053 | 9,66362E-31 | 2,16648E-28 | NOVA2 | protein_coding |
| ENSG00000092068 | 84,38062302 | 3,143740537 | 0,271623796 | 11,57387748 | 5,58981E-31 | 1,26978E-28 | SLC7A8 | protein_coding |
| ENSG00000222078 | 80,39645476 | 7,2013185 | 0,614424975 | 11,72041957 | 1,00175E-31 | 2,41978E-29 | RN7SKP110 | misc_RNA |
| ENSG00000007516 | 219,3636063 | 2,11187215 | 0,179152029 | 11,78815645 | 4,49268E-32 | 1,12484E-29 | BAIAP3 | protein_coding |
| ENSG00000091536 | 86,52467451 | 4,059151241 | 0,344165709 | 11,79417687 | 4,1827E-32 | 1,05493E-29 | MYO15A | protein_coding |
| ENSG00000183914 | 93,0181299 | 3,768417214 | 0,316930359 | 11,89036363 | 1,32827E-32 | 3,53186E-30 | DNAH2 | protein_coding |
| ENSG00000226455 | 57,5752517 | 5,343384667 | 0,447301657 | 11,94581909 | 6,82744E-33 | 1,82959E-30 | ENSG00000226455 | lncRNA |
| ENSG00000224940 | 146,5391305 | 2,512517422 | 0,209387553 | 11,9993638 | 3,58038E-33 | 9,74687E-31 | PRRT4 | protein_coding |
| ENSG00000148357 | 138,5209186 | 4,125761119 | 0,343057913 | 12,0264275 | 2,58089E-33 | 7,13927E-31 | HMCN2 | protein_coding |
| ENSG00000132692 | 133,439112 | 2,29347351 | 0,189976463 | 12,07240873 | 1,47745E-33 | 4,15395E-31 | BCAN | protein_coding |
| ENSG00000072163 | 170,1484635 | 2,181652915 | 0,179714967 | 12,13951711 | 6,52097E-34 | 1,91176E-31 | LIMS2 | protein_coding |
| ENSG00000124772 | 128,4547558 | 3,097003512 | 0,252996243 | 12,24130238 | 1,87008E-34 | 5,57786E-32 | CPNE5 | protein_coding |
| ENSG00000173175 | 162,3054381 | 2,021347012 | 0,165113207 | 12,24218856 | 1,84977E-34 | 5,56568E-32 | ADCY5 | protein_coding |
| ENSG00000107738 | 81,07975052 | 3,823598001 | 0,312292897 | 12,24362781 | 1,81725E-34 | 5,51623E-32 | VSIR | protein_coding |
| ENSG00000132561 | 193,3244932 | 2,136843112 | 0,174030233 | 12,27857412 | 1,18061E-34 | 3,64829E-32 | MATN2 | protein_coding |
| ENSG00000166750 | 165,4849974 | 2,155172181 | 0,175508998 | 12,27955378 | 1,1664E-34 | 3,63715E-32 | SLFN5 | protein_coding |
| ENSG00000161681 | 243,6822011 | 2,020059557 | 0,164054526 | 12,31334241 | 7,67764E-35 | 2,48444E-32 | SHANK1 | protein_coding |
| ENSG00000135636 | 111,6216189 | 3,042833254 | 0,24471757 | 12,43406127 | 1,70735E-35 | 5,74154E-33 | DYSF | protein_coding |
| ENSG00000152527 | 187,3898911 | 2,478505771 | 0,197000348 | 12,58122536 | 2,67843E-36 | 9,28007E-34 | PLEKHH2 | protein_coding |
| ENSG00000095303 | 109,9574369 | 3,371878681 | 0,26629336 | 12,66227095 | 9,56906E-37 | 3,49179E-34 | PTGS1 | protein_coding |
| ENSG00000116016 | 132,9139965 | 2,458495127 | 0,194127513 | 12,66433123 | 9,32113E-37 | 3,45753E-34 | EPAS1 | protein_coding |
| ENSG00000186642 | 110,1477549 | 3,205346315 | 0,252309297 | 12,70403572 | 5,61574E-37 | 2,11676E-34 | PDE2A | protein_coding |
| ENSG00000165272 | 226,5041508 | 2,005955201 | 0,156919261 | 12,78335872 | 2,03105E-37 | 7,82777E-35 | AQP3 | protein_coding |
| ENSG00000168016 | 95,5553875 | 2,95922304 | 0,230309133 | 12,84891746 | 8,72223E-38 | 3,47885E-35 | TRANK1 | protein_coding |
| ENSG00000156042 | 198,0241579 | 2,189970516 | 0,170020114 | 12,88065552 | 5,78422E-38 | 2,39042E-35 | CFAP70 | protein_coding |
| ENSG00000248485 | 112,6022887 | 3,278538902 | 0,253965086 | 12,90940797 | 3,98352E-38 | 1,66633E-35 | PCP4L1 | protein_coding |
| ENSG00000185101 | 130,0980583 | 3,048258803 | 0,235213233 | 12,95955489 | 2,07448E-38 | 9,00717E-36 | ANO9 | protein_coding |
| ENSG00000117115 | 241,0472994 | 2,640119836 | 0,203642983 | 12,96445275 | 1,94614E-38 | 8,55829E-36 | PADI2 | protein_coding |
| ENSG00000105559 | 170,0018099 | 2,543318296 | 0,195699781 | 12,99602017 | 1,28878E-38 | 5,89418E-36 | PLEKHA4 | protein_coding |
| ENSG00000154188 | 230,9354495 | -2,223247022 | 0,170204496 | -13,06221087 | 5,41325E-39 | 2,57889E-36 | ANGPT1 | protein_coding |
| ENSG00000006611 | 112,5025588 | 4,477258488 | 0,337052996 | 13,28354454 | 2,88412E-40 | 1,57029E-37 | USH1C | protein_coding |
| ENSG00000142609 | 171,8405596 | 2,669427171 | 0,199698659 | 13,36727638 | 9,39172E-41 | 5,28107E-38 | CFAP74 | protein_coding |
| ENSG00000197558 | 349,4068344 | 2,588262449 | 0,192966383 | 13,41302259 | 5,07284E-41 | 2,94921E-38 | SSPOP | transcribed_unitary_pseudogene |
| ENSG00000125637 | 149,1336634 | 2,617862982 | 0,194118702 | 13,48588754 | 1,89373E-41 | 1,15994E-38 | PSD4 | protein_coding |
| ENSG00000103485 | 233,7215818 | 2,314452459 | 0,171448505 | 13,49940299 | 1,57648E-41 | 9,83182E-39 | QPRT | protein_coding |
| ENSG00000163520 | 107,4344374 | 4,032282863 | 0,297710991 | 13,5442862 | 8,56423E-42 | 5,54268E-39 | FBLN2 | protein_coding |
| ENSG00000178038 | 272,5129616 | 2,029702527 | 0,14940886 | 13,58488735 | 4,92289E-42 | 3,24731E-39 | ALS2CL | protein_coding |
| ENSG00000111424 | 145,6025348 | 2,423576183 | 0,178388718 | 13,58592744 | 4,85345E-42 | 3,24731E-39 | VDR | protein_coding |

| | | | | | | | | | |
|---|---|---|---|---|---|---|---|---|---|
| ENSG00000172350 | 158,7075642 | 2,583557558 | 0,189645388 | 13,62309722 | 2,9192E-42 | 2,0435E-39 | ABCG4 | | protein_coding |
| ENSG00000132470 | 325,6261233 | 2,093802101 | 0,151748719 | 13,79782391 | 2,62662E-43 | 2,09525E-40 | ITGB4 | | protein_coding |
| ENSG00000135472 | 151,1634624 | 3,084655129 | 0,21944599 | 14,05655728 | 7,02201E-45 | 6,17594E-42 | FAIM2 | | protein_coding |
| ENSG00000198959 | 107,8962046 | 3,779685009 | 0,264530156 | 14,28829537 | 2,58869E-46 | 2,6116E-43 | TGM2 | | protein_coding |
| ENSG00000143603 | 306,6005563 | 2,546642217 | 0,176011297 | 14,46862939 | 1,91245E-47 | 1,98785E-44 | KCNN3 | | protein_coding |
| ENSG00000143850 | 242,2820346 | 3,460383699 | 0,237623719 | 14,56245071 | 4,86808E-48 | 5,566E-45 | PLEKHA6 | | protein_coding |
| ENSG00000171992 | 168,4756059 | 3,246330062 | 0,215949596 | 15,03281374 | 4,47583E-51 | 6,67502E-48 | SYNPO | | protein_coding |
| ENSG00000116983 | 186,6042251 | 3,378122749 | 0,218315325 | 15,47359418 | 5,23034E-54 | 9,44242E-51 | HPCAL4 | | protein_coding |
| ENSG00000185567 | 1345,784646 | 2,199629236 | 0,14206726 | 15,48301303 | 4,51807E-54 | 8,60969E-51 | AHNAK2 | | protein_coding |
| ENSG00000104888 | 160,2314889 | 4,518038669 | 0,286691555 | 15,75923182 | 5,935E-56 | 1,19751E-52 | SLC17A7 | | protein_coding |
| ENSG00000129244 | 195,248251 | 2,969516861 | 0,185951966 | 15,96926843 | 2,09225E-57 | 5,12617E-54 | ATP1B2 | | protein_coding |
| ENSG00000021300 | 207,9633419 | 3,012957121 | 0,185106114 | 16,27691842 | 1,43937E-59 | 3,79783E-56 | PLEKHB1 | | protein_coding |
| ENSG00000161642 | 254,3086608 | 3,213360662 | 0,197355457 | 16,2820968 | 1,32258E-59 | 3,7805E-56 | ZNF385A | | protein_coding |
| ENSG00000168071 | 207,1734025 | 3,258224501 | 0,199433799 | 16,33737366 | 5,35086E-60 | 1,66854E-56 | CCDC88B | | protein_coding |
| ENSG00000177098 | 244,1807548 | 3,416420959 | 0,195737513 | 17,45409407 | 3,20421E-68 | 1,22119E-64 | SCN4B | | protein_coding |
| ENSG00000079308 | 651,7935611 | 2,377366245 | 0,131680861 | 18,05399985 | 7,3379E-73 | 3,59568E-69 | TNS1 | | protein_coding |
| ENSG00000064300 | 429,5951477 | 3,204947199 | 0,176178998 | 18,19142598 | 6,03494E-74 | 3,45008E-70 | NGFR | | protein_coding |
| ENSG00000065361 | 671,8656728 | 2,042016772 | 0,110659112 | 18,45321849 | 4,913E-76 | 4,21302E-72 | ERBB3 | | protein_coding |
| ENSG00000149260 | 516,7088159 | 2,615986706 | 0,140111298 | 18,67077626 | 8,55995E-78 | 9,78717E-74 | CAPN5 | | protein_coding |
| ENSG00000173546 | 642,8241115 | 3,048026416 | 0,142992049 | 21,31605524 | 8,0571E-101 | 1,38183E-96 | CSPG4 | | protein_coding |
| ENSG00000112182 | 5821,12613 | 6,543670056 | 0,235710853 | 27,76142881 | 1,2683E-169 | 4,3504E-165 | BACH2 | | protein_coding |
