## Supplementary material for "Selective persistence of HIV-1-infected T cell clones can occur through immune reprogramming driven by defective, transcriptionally active proviruses": STable2

| name | 5'-3' sequence |
|---|---|
| oligo 1 | gcggacaataggatccctctctccttctagcctc |
| oligo 2 | taacttcgtatagcatacattatacgaagttattggaagggctaatttgg |
| oligo 3 | atgtatgctatacgaagttatcggaccgcccgggctagcatgcggccttgacggccttc |
| oligo 4 | ggatcctattgtccgcggtcgacctgggcctcatg |
| oligo 5 | gctcgctcatctctctccttctagcctccg |
| oligo 6 | tctctagcaggatcctattgtccgcgg |
| oligo 7 | aaggagagagatgagcgagctgattaagg |
| oligo 8 | ccttccactattccttcgggcctgtc |
| oligo 9 | aataggatcctgctagagattttccacac |
| oligo 10 | cccgaaggaatagtggaagggctaatttggtc |
| oligo 11 | gatccataacttcgtatagcatacattatacgaagttatccgc |
| oligo 12 | ggataacttcgtataatgtatgctatacgaagttatg |
| oligo 13 | tccccgcggcacaatctcctgcctcagcc |
| oligo 14 | tccccgcggggagacagggtatcactttg |
| oligo 15 | tcccccgggcttcctcagctcaaaaggag |
| oligo 16 | gtacggaccgcagcaggaagaggttgcagt |
| oligo 17 | gtccgnnnnnnnncgtatcgatatatctcg |
| oligo 18 | gaccgagatatatcgatacgnnnnnnnncg |
| oligo 19 | agcgagctgattaaggagaac |
| oligo 20 | ctctagtctaggatctactggc |
| oligo 21 | tggtaacaggattagcagagc |
| oligo 22 | tcgttttctggtgctgtatgg |
| oligo 23 | agcgagctgattaaggagaac |
| oligo 24 | cagccgagtgttttcttctgc |
| oligo 25 | tcctactttctttttggctttc |
| oligo 26 | aagttacgctatatttatac |
| oligo 27 | cagtccactgcaccaacatc |
| oligo 28 | tctccacgatcaaagtcacg |
| oligo 29 | tgaccagcggaaaaaggata |
| oligo 30 | tgagggctccaggaatgag |
| oligo 31 | ttaggatgcagggaactggg |
| oligo 32 | tccatgtctgaatggattgccagagcc |
| oligo 33 | ACCCAGCGCAGGCAA |
| oligo 34 | GGTTTACAATCTCGCTCG |
| oligo 35 | AAGCCAGACAGCTGGGCC |
| oligo 36 | GCGTCATGGAGTACCAC |
| oligo 37 | CACTGAACTGGGATTCAAAC |
| oligo 38 | CTTCTGTCACCGACTCTGC |
| oligo 39 | CTGAGCTTGTATCCACACAG |
| oligo 40 | TCTCTAGCAGTGGCGCCC |
| oligo 41 | TCAGCAAGCCGAGTCCTGCGTCGAGA |

| description |
|---|
| Fwd HIV reporter 5' LTR-leader Gibson |
| Rev HIV reporter 5' LTR-leader Gibson |
| Fwd pMK backbone_step1 Gibson |
| Rev pMK backbone_step1 Gibson |
| Fwd pMK backbone_step2 Gibson |
| Rev pMK backbone_step2 Gibson |
| Fwd HIV reporter bfp-2A-tat Gibson |
| Rev HIV reporter bfp-2A-tat Gibson |
| Fwd HIV reporter 3' LTR Gibson |
| Rev HIV reporter 3' LTR Gibson |
| Fwd HIV reporter 3' loxP Gibson |
| Rev HIV reporter 3' loxP Gibson |
| Fwd 3' BACH2 homology arm |
| Rev 3' BACH2 homology arm |
| Fwd 5' BACH2 homology arm |
| Rev 5' BACH2 homology arm |
| Fwd HIV reporter 8bp random barcode |
| Rev HIV reporter 8bp random barcode |
| Fwd BACH2 clone validation HIV reporter |
| Rev BACH2 clone validation HIV reporter |
| Fwd BACH2 clone validation pMK backbone |
| Rev BACH2 clone validation pMK backbone |
| Fwd Southern bfp-specific probe |
| Rev Southern bfp-specific probe |
| Fwd Southern genomic BACH2-specific probe |
| Rev Southern genomic BACH2-specific probe |
| Fwd qPCR BACH2 exon6 |
| Rev qPCR BACH2 exon6 |
| Probe qPCR BACH2 exon6 (FAM/BHQ1) |
| Fwd qPCR BACH2 exon4/5 |
| Rev qPCR BACH2 exon4/5 |
| Probe qPCR BACH2 exon4/5 (FAM/BHQ1) |
| Fwd qPCR STAT5B exon1 |
| Rev qPCR STAT5B exon2 |
| Probe qPCR STAT5B exon1/2 (FAM/BHQ1) |
| Fwd qPCR STAT5B exon10 |
| Rev qPCR STAT5B exon11 |
| Probe qPCR STAT5B exon10/11 (HEX/BHQ1) |
| Rev STAT5B qPCR exon 2 |
| Fwd qPCR HIV LTR |
| Probe qPCR LTR/STAT5B chimeric transcript |
